# Small intestinal microbiota of undernourished women perturbs placental development in mice

**DOI:** 10.64898/2026.08.17.745217

**Authors:** Reyan Coskun, ZeNan L. Chang, Kali M. Pruss, Haoxin Liu, Athziri Marcial Rodríguez, Evan M. Lee, Michael S. Diamond, Tahmeed Ahmed, Michael J. Barratt, Jeffrey I. Gordon

**Affiliations:** The Edison Family Center for Genome Sciences and Systems Biology, Washington University School of Medicine; St. Louis, MO 63110, USA; The Newman Center for Gut Microbiome and Nutrition Research, Washington University School of Medicine; St. Louis, MO 63110, USA; Division of Gastroenterology, Department of Medicine, Washington University School of Medicine; St. Louis, MO 63110, USA; Departments of Medicine, Molecular Microbiology and Pathology and Immunology, Washington University School of Medicine; St. Louis, MO 63110, USA; International Center for Diarrhoeal Disease Research, Bangladesh (icddr,b); Dhaka 1212, Bangladesh

## Abstract

Children of undernourished women have impaired pre-and postnatal growth. Undernourished women and children have a high incidence of environmental enteric dysfunction (EED), an enteropathy characterized by gut barrier dysfunction and systemic inflammation.

Here, we employ gnotobiotic mice to compare the effects of bacterial consortia cultured from the duodenal microbiota of Bangladeshi women with EED and their healthy counterparts. Female mice harboring the EED-derived consortium exhibited fetal and placental growth restriction.

Transcriptomic and proteomic analyses disclosed pronounced effects of the EED-derived consortium on the decidual component of the maternal-fetal interface involving tissue-resident uterine natural killer (uNK) cells and disruption of TGF-β signaling between uNK and decidual stromal cells. Co-housing mice with EED and healthy consortia ameliorated these effects, disclosing bacterial targets to improve prenatal development.

## INTRODUCTION

Undernutrition, which can manifest as either wasting (defined by anthropometry as low weight-for-length Z score (WLZ)), stunting (low length-for-age Z-score (LAZ)), or both conditions together, affects approximately 200 million children worldwide, with the greatest burden of disease in Africa and South Asia (*1*). The long-term sequelae of undernutrition include, but are not limited to, impaired cognitive development and immune dysfunction, which impose significant societal burdens (*2*, *3*). Stunting has an intergenerational component; women who were undernourished are more likely to have stunted children (*4*). Maternal weight before pregnancy is also a strong predictor of intrauterine growth restriction (IUGR) and low birth weight (*5*, *6*).

While epidemiologic studies have emphasized food insecurity as a key contributing factor to undernutrition, current nutritional interventions have had limited success in alleviating stunting, suggesting other factors contribute to disease pathogenesis (*7*, *8*). Several studies have demonstrated an association between stunting and ‘environmental enteric dysfunction’ (EED) – an enteropathy associated with blunting of small intestinal (SI) villi, reduced surface area for nutrient absorption, and disruption of the gut mucosal barrier resulting in local and systemic inflammation (*9–12*). The Bangladeshi EED (BEED) study performed endoscopy (esophagogastroduodenoscopy, EGD) to obtain duodenal mucosal biopsies and duodenal aspirates from (i) stunted children and (ii) undernourished women (BMI <18.5 kg/m^2^) of childbearing age (18-45 years old) who failed a standard nutritional intervention. The results revealed that the majority of stunted children and undernourished women had EED (*11*, *13*).

Comparisons of pregnant germ-free (GF) C57BL/6J mice and their conventionally-raised (CONV-R) counterparts harboring a normal mouse microbiota have shown that placental weights are significantly reduced in GF animals at mid-gestation and fetal weights are reduced in late gestation (*14–17*). Angiogenesis was mainly affected in the fetal-derived labyrinth zone (LZ) and the junctional zone (JZ) of the placenta, but not in the maternal-derived decidua and mesometrial lymphoid aggregate of pregnancy (MLAp). The deficit in placental vasculature in GF mice was rescued by the introduction of the gut microbiota of CONV-R mice (*i.e.* yielding conventionalized animals, CONV-D) (*14*). These findings raise the possibility that an EED-associated microbiota may affect pre-and postnatal development.

We have used culture-independent methods to characterize bacterial composition in the duodenal microbiota of undernourished women with EED and their healthy counterparts. The absolute abundances of several taxa in the EED-associated microbial community were significantly correlated with levels of duodenal mucosal protein biomarkers/mediators of immunoinflammatory responses (*18*). Bacterial strains were subsequently cultured from the duodenal aspirates of these women [to generate the <u>a</u>dult <u>S</u>mall <u>I</u>ntestinal-<u>L</u>ow BMI (aSI-L) and <u>a</u>dult <u>S</u>mall <u>I</u>ntestinal-<u>H</u>ealthy BMI (aSI-H) consortia]. The bacterial consortium from undernourished women compromised gut barrier integrity and elicited systemic inflammation in non-pregnant dams. In addition, dam-to-pup transmission of the EED-derived consortium impaired postnatal offspring development (*18*). Several of these changes were previously documented in mice harboring a bacterial consortium from children with EED (*18*).

In the current study, we use gnotobiotic mice colonized with either the EED-donor-derived aSI-L or healthy-donor-derived aSI-H consortia to model the effects of an EED-associated SI microbiota on prenatal development. Compared to dams harboring the healthy women’s aSI-H consortium, dams colonized with the undernourished women’s aSI-L consortium exhibited reductions in placental and fetal weights during the course of pregnancy. The results reveal that the effects of the aSI-L consortium largely manifest in the maternal rather than fetal component of the maternal-fetal interface (MFI), with alterations in both tissue-resident uterine and conventional natural killer (NK) cells and decidual stromal cells. Specifically, signaling pathways involving TGF-β, and also FGF-2 and Wnt, that normally mediate NK-decidual stromal cell communications for programmed MFI remodeling, are reduced in the aSI-L colonized dams. These effects are ameliorated by co-housing aSI-L mice with aSI-H animals before pregnancy.

## RESULTS

### Impaired fetal and placental growth in aSI-L compared to aSI-H mice

To evaluate the impact of an EED-associated SI microbiota on prenatal development, we switched C57BL/6J GF adult female mice from a standard mouse chow to a diet representative of that consumed by Bangladeshi adults (see *Methods*). Three days after the diet switch, mice were gavaged with either the aSI-L or aSI-H bacterial consortium [details of the cultured consortia are described in Pruss, Chang *et al* (*18*)]. Timed matings were initiated 17 days post-gavage (**Fig. 1A**).

**Figure 1.**
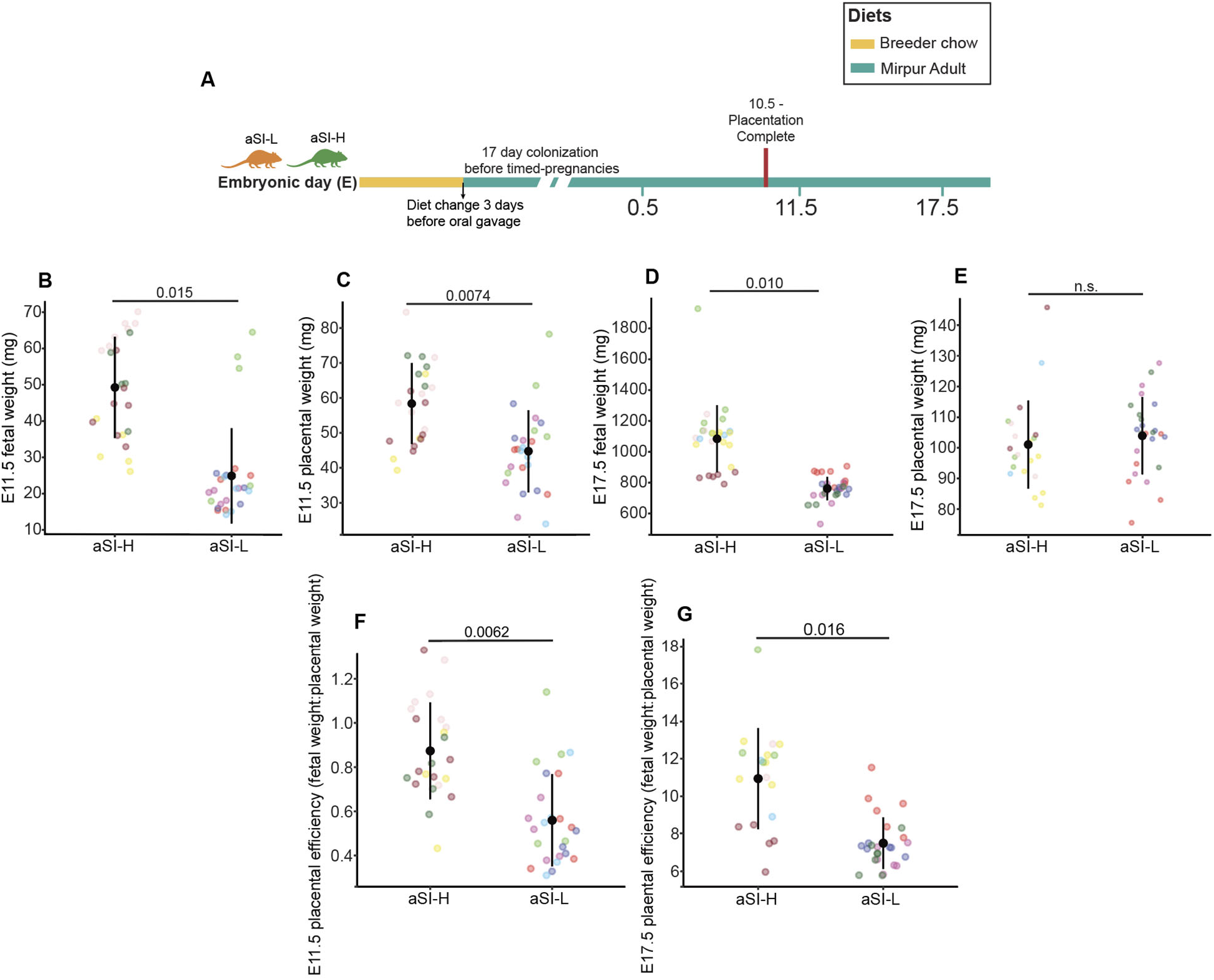
Intrauterine growth in aSI-L and aSI-H dams. **(A)** Experimental design. **(B)** E11.5 fetal weights [aSI-L: n= 5 dams, 26 fetuses, 24.9 ± 13.1 mg (mean ± SD); aSI-H: 4 dams, 24 fetuses, 49.3 ± 13.9 mg. **(C)** Corresponding E11.5 placental weights [aSI-L: 44.7 ± 11.7 mg (mean ± SD); aSI-H: 58.3 ± 11.6 mg]. **(D)** E17.5 fetal weights [aSI-L: 4 dams, 31 fetuses, 761.3 ± 75.5 mg; aSI-H: 5 dams, 27 fetuses, 1084.1 ± 216.9 mg]. **(E)** Corresponding E17.5 placental weights [aSI-L: 103.4 ± 12.6 mg; aSI-H: 100.6 ± 14.4 mg]. **(F,G**) Placental efficiency (weight of fetus produced per weight of placenta) at E11.5 (panel **F**) and E17.5 (panel **G**). Linear mixed model for pairwise comparisons (*Fetal or placental weight, or placental efficiency ∼ Microbiota + Litter Size + (1 | Litter ID)*). Each litter is represented by different colors.

Fetal weights were significantly reduced in aSI-L dams at embryonic day (E) 11.5, a time just after completion of placentation (n = 50 fetuses representing 4-5 litters/group, p = 0.015; linear mixed model) and at E17.5, a time near the end of gestation for C57BL/6J mice (n = 58 fetuses representing 4-5 litters/group, p = 0.01) (**Fig. 1B**,**D**) (*19*, *20*). aSI-L placental weights were significantly reduced at E11.5 (n = 48 fetuses representing 4-5 litters/group, p = 0.0074), but not at E17.5 (n = 47 fetuses representing 4-5 litters/group, p = 0.16) (**Fig. 1C**,**E**). However, there were significant decreases in aSI-L ‘placental efficiency’, defined as grams of fetus produced per gram of placenta (*21*), at both E11.5 (n = 47-48 fetuses representing 4-5 litters/group, p = 0.0062) and E17.5 (n = 47 fetuses representing 4-5 litters/group, p = 0.016) (**Fig. 1F**,**G**). There were no statistically significant differences in litter sizes at either timepoint (p = 0.23 and 0.15; Mann-Whitney-Wilcoxon test) (**fig. S1A**,**B**) nor the distribution of the number of matings until a successful pregnancy was achieved (D-statistic = 0.18, p = 0.99; Kolmogorov-Smirnov test) **(fig. S1C)**. In addition, fetal sex had no significant effect on fetal and placental weight differences within the aSI-L or aSI-H groups at either E11.5 or E17.5 (linear mixed models) (**fig. S1D-G**).

### Pregnancy ameliorates intestinal inflammation

Pregnancy has well-described immunomodulatory effects (*22*). In pregnant dams, there were no differences in villus length, crypt depth, or the villus-to-crypt ratio along the length of the SI (**fig. S2A**), or in intestinal levels of the EED biomarker lipocalin 2 (LCN2), between animals harboring the aSI-H versus aSI-L consortia (**fig. S2B**). To further assess the impact of pregnancy on maternal enteropathy, we performed single-nucleus RNA sequencing (snRNA-seq) of maternal duodenal and ileal segments harvested from dams at E11.5 and E17.5 (n = 4 mice/timepoint/group). We focused our analysis on the enterocytic lineage because it contained 85% and 98% of all differentially expressed genes [DEGs, **|**log_2_(fold-difference)|>0.5, FDR < 0.05, Wald test] in aSI-L versus aSI-H mice at E11.5 and E17.5, respectively (**table S1D-G**). Compared to their nonpregnant counterparts, the number of DEGs in the crypt, villus base, mid-villus, and villus tip enterocyte subclusters in aSI-L versus aSI-H mice was reduced, most markedly just before parturition at E17.5 (**fig. S3A**; data from non-pregnant controls are described in *18*). The majority of DEGs in the duodenum and ileum of non-pregnant aSI-L mice are involved in inflammatory responses; differences in the expression of these genes diminished between the two treatment groups as pregnancy progressed to E17.5 (**fig. S3B**; see **table S1D-G** for a complete list of DEGs across the treatment groups as a function of enterocyte subcluster/intestinal segment/time point). Moreover, at E17.5, there were no significant differences between aSI-L and aSI-H dams in duodenal and ileal levels of two determinants of mucosal barrier integrity – IL-22 and the tight-junction scaffold protein zonula occludens 1 (ZO-1) (**fig. S3C,D**). Both biomarkers were affected in the duodenums of non-pregnant aSI-L compared to aSI-H dams (*18*).

The absolute abundances of several bacterial species were significantly elevated in aSI-L pregnant dams and non-pregnant controls compared to aSI-H dams and non-pregnant controls (**fig. S4, table S1H**). However, the amelioration of intestinal inflammation with pregnancy could not be attributed to shifts in the gut microbiota, as there were no statistically significant changes in the absolute abundances of bacterial strains in the cecums of pregnant compared to non-pregnant mice across treatment groups (FDR > 0.05 for all species between E17.5 pregnant dams and non-pregnant controls, Benjamini Hochberg-corrected two-sided Wilcoxon Rank-Sum tests, **table S1J**).

### Characterization of fetal-derived compartments within the maternal-fetal interface (MFI)

We initially characterized the fetal-derived component of E11.5 and E17.5 MFIs after manually removing the decidual tissue. The fetal LZ contains a dense network of capillaries where nutrient and oxygen exchange occurs; its primary cell types are syncytiotrophoblasts (SynT) and sinusoidal trophoblastic giant cells (S-TGC) (*23*). The fetal JZ directly contacts both the LZ and the maternal-derived tissue. Its three main cell types – spongiotrophoblasts (SpT), glycogen cells (GC), and parietal TGC (P-TGC) – contribute to the production of hormones, growth factors, and cytokines essential for the progression of pregnancy (*24*, *25*).

Histological analyses disclosed that the dissected fetal placenta consisted principally of the LZ and JZ. However, the preparations also had residual maternal components as part of the normal murine MFI (see **Supplementary Results** for how maternal and fetal-derived cell types were identified and their proportions as determined from our snRNA-seq tables). Findings from bulk RNA-seq, snRNA-seq, proteomic, and histological analyses of E11.5 and E17.5 ‘fetal’ placental preparations are described in **Supplementary Results** (**figs. S4-S9, tables S2-S4**) and summarized in **table S4**; they revealed no major differences in trophoblast function or vascular remodeling. Rather, the effects of colonization status were largely confined to maternal-derived cell types.

### aSI-L colonization results in perturbations in uNK and decidual stromal cells populations and gene expression

Movat’s pentachrome staining with optical density (OD) normalization revealed stromal changes characterized by decreased collagen and reticular fibers in the E11.5 placental decidua and MLAp of aSI-L compared to aSI-H dams (**fig. S5E**). Additionally, snRNA-seq of the E11.5 placentas revealed that maternal-derived cells (endothelial, stromal) had the highest numbers of DEGs (**fig. S6G**). Therefore, we conducted snRNA-seq of E11.5 decidual/MLAp tissue preparations after the fetal-derived LZ and JZ were removed. We identified 13 cell clusters (**Fig. 2A**, **table S5A-C**, *27*). There were no differences in the proportions of cell clusters between aSI-L and aSI-H mice based on scCODA analysis (**fig. S10A**, **table S5D**, see *Methods*). The NK cell and two decidual cell clusters (*i.e.* proliferating and non-proliferating) had the largest number of DEGs (defined as **|**log_2_(fold-difference)|>0.5, FDR < 0.05, Wald test) (**Fig. 2B**, **table S5E**).

**Figure 2.**
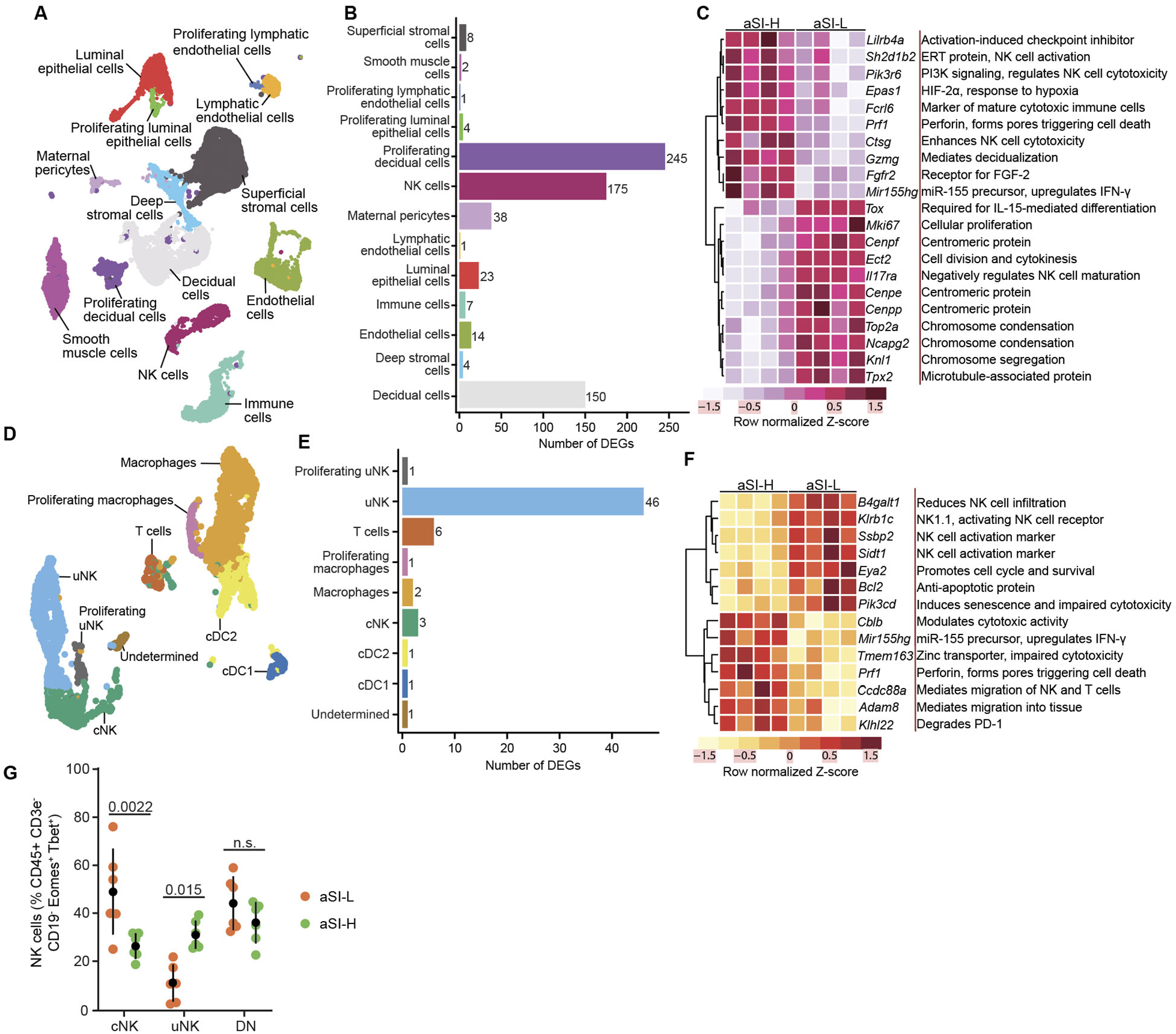
snRNA-seq and flow cytometry of E11.5 aSI-L deciduas/MLAps reveal alterations in uNK cell transcriptomic responses and their representation. **(A)** UMAP representation of 16,273 nuclei across eight E11.5 decidual/MLAp samples (4 aSI-L and 4 aSI-H; pooled across entire litter for each sample). Nuclei were assigned to 13 cell clusters. **(B)** Number of differentially expressed genes (DEGs) in each cell cluster identified by pseudobulk analysis of E11.5 aSI-L versus aSI-H deciduas/MLAps (DESeq2, |log_2_(fold-difference)| > 0.5, FDR p < 0.05, Wald test with BH correction). **(C)** DEGs identified when comparing all aSI-L to all aSI-H NK cells (|log_2_(fold-difference)| > 1). **(D)** UMAP representation of NK and immune cell clusters subset and re-clustered from the E11.5 pooled decidual/MLAp table (n = 2,901 nuclei). **(E)** Number of DEGs identified by pseudobulk analysis of the E11.5 aSI-L versus aSI-H cell clusters shown in panel **D** (DESeq2, |log_2_(fold-difference)| > 0.5, FDR < 0.05, Wald test with BH correction). **(F)** DEGs identified when comparing aSI-L to aSI-H uNK cells (|log_2_(fold-difference)| > 0.5). **(G)** Proportion of cNK, uNK, and DN NK cell populations in aSI-L compared to aSI-H deciduas/MLAps (n = 6 dams/treatment group, Mann-Whitney-Wilcoxon test).

### aSI-L uNK cells display transcriptional signatures of over-activation and anti-apoptotic responses

The E11.5 aSI-L NK cell cluster contained DEGs indicative of (i) increased proliferation (*Mki67, Fgfr2*) and mitosis [specifically those related to centromeric proteins (*Cenpe*, *Cenpf*, *Cenpp*), chromosome condensation (*Ncapg2*, *Top2a*), and microtubule, kinetochore, and kinesin formation (*Ect2*, *Knl1*, *Tpx2*)], (ii) decreased cell maturation (*Tox*, *Il17ra*, *Fcrl6*), (iii) decreased cytotoxic granule proteins (*Prf1*, *Ctsg*), (iv) decreased IFN-γ production (*Mir155hg*, *Epas1*), (v) decreased activation (*Lilrb4a*, *Sh2d1b2*, *Pik3r6*) and (vi) decreased decidualization (*Gzmg*) (|log_2_(fold-difference)| > 1, FDR < 0.05, Wald test) (**Fig. 2C**, **table S5E**).

We subsequently subclustered the immune cell clusters (NK and non-NK) and re-assigned the resulting nuclei based on known marker genes for decidual immune cells (see *Methods*, *28*). This yielded clusters with (i) specialized populations of uNK cells and conventional NK (cNK) cells, (ii) macrophages, (iii) conventional type 1 and 2 dendritic cells (cDC1 and cDC2), (iv) T cells, and (v) one that we could not classify based on reported marker genes (**Fig. 2D**, **table S5F-H**). uNK cells had the highest number of DEGs based on pseudobulk analysis (**|**log_2_(fold-difference)|>0.5, FDR < 0.05, Wald test) (**Fig. 2E**, **table S5I**). DEGs in the aSI-L uNK cell subset provided evidence of (i) decreased infiltration (*B4galt1*, *Adam8*, *Ccdc88a*), (ii) decreased cytotoxic granule and IFN-γ production (*Prf1*, *Tmem163*, *Cblb*, *Mir155hg*), (iii) increased activation (*Ssbp2*, *Sidt1*, *Klrb1c* [NK1.1]), and (iv) increased anti-apoptotic signals (*Bcl2*, *Eya2*, *Pik3cd*, *Klhl22*) (**Fig. 2F**, **table S5I**). Together, these results suggest that compared to the aSI-H group, aSI-L uNK cells exhibit transcriptional responses indicative of persistent activation.

### E11.5 aSI-L deciduas have lower proportions of uNK and higher proportions of cNK cells

We used flow cytometry to further characterize differences in NK cells at E11.5 between aSI-L and aSI-H dams (see **fig. S11A** for gating schematic). In mice, expression of the collagen I-binding integrin CD49b marks cNK cells that primarily circulate then enter tissues when recruited (*28*); tissue resident uNK cells instead express CD49a, an integrin that binds collagen IV and laminin in the extracellular matrix (ECM) (*29*, *30*). We identified significantly more cNK cells and fewer uNK cells in aSI-L versus aSI-H deciduas (n = 6 litters/group, p = 0.015 and 0.002, respectively; Mann-Whitney-Wilcoxon test) (**Fig. 2G**). An additional CD49a^−^CD49b^−^ double-negative (DN) population was present, athough the size of this population was not significantly different between the two treatment groups. DN NK cells have been reported as a phenotypically and functionally immature NK cell population with lower production of IFN-γ, granzyme B, and perforin after PMA-ionomycin stimulation (*31*). The CD49b^+^ cNK population had elevated median NK1.1 expression relative to uNK and DN NK cells (**fig. S10B**) and were more prominently marked by intravenously administered anti-CD45 antibodies (CD45-IV) prior to euthanasia (see *Methods*) (**fig. S10C**), supporting their circulating origin.

Additional investigation into NK cell maturation, as defined by the CD11b and CD27 surface markers (*i.e.* CD11b^−^ CD27^−^ → CD11b^−^ CD27^+^ → CD11b^+^ CD27^+^ → CD11b^+^ CD27^−^) (*32*), demonstrated no significant differences between the aSI-L and aSI-H treatment groups during all stages of maturation in the uNK, cNK, and DN populations (**fig. S10D-F**).

We also examined T-cell, B-cell, and neutrophil populations in the decidua/MLAp, as disturbances in their respective intestinal immune cell lineages have been implicated in EED (*18*, *33*, *34*). Although Th17 cells are elevated in the intestinal mucosa of non-pregnant aSI-L mice (*18*), we found no significant differences in the proportions of any decidual T cell population, including Treg, Th17, and γδ T cells, between the two groups (**fig. S10G**). We also found no significant difference in proportions of decidual B cells and neutrophils (**fig. S10G,H**). Among the myeloid populations in the decidua/MLAp, the only cell type that exhibited a significant difference in relative abundance was Ly6C^high^ monocytes, which were decreased in aSI-L mice (p = 0.015) (**fig. S10H**). Ly6C^high^ monocytes are pro-inflammatory, release cytokines (*e.g*. IL-1, IL-18, IL-15, and CCL2), and have a high antimicrobial capability during bacterial infections (*35*, *36*).

Together, the transcriptomic and flow cytometry results led us to conclude that, compared to dams harboring the aSI-H consortium, aSI-L dams have reduced uNK abundance associated with a transcriptional signature of persistent activation, alongside of increased recruitment of cNK cells to the MFI. Progesterone increases cNK cell infiltration into the decidua and helps upregulate expression of the tissue-resident uNK cell marker CD49a (*37*). We observed that estradiol and progesterone concentrations were significantly increased in homogenates prepared from E11.5 aSI-L decidual/MLAp tissue (n = 4 MFIs/dam, 5 litters/group; p < 0.05, linear mixed model) (**fig. S12A**,**B**). The increased decidual concentrations of progesterone could be one factor that contributes to increased entry of cNK cells into aSI-L deciduas.

### aSI-L decidual cells have altered expression of proliferation and ECM maintenance-related genes

The E11.5 decidual/MLAp snRNA-seq dataset disclosed that decidual cells also had numerous DEGs (**Fig. 2A**,**B**, **table S5E**). Proliferating decidual cells in aSI-L dams had DEGs that provided evidence for (i) increased cellular proliferation (*Prc1*, *Top2a*, *Rttn*, *Cdknc1*, *Usp29*, *K1fc3, Wnk2*), tissue growth (*Igf2*, *Peg10*, *Igf2bp2*), and inhibition of apoptosis (*Trim71*); (ii) perturbations in decidual stroma that mediate trophoblast invasion (*C1qb*, *Igf2bp3*, *Erv3*); (iii) effects on cellular junctions (*Dsg2*, desmoglein-2); and (iv) altered lipid metabolism (*Lipc*, *Hnf1aos1*, *Pparg*) and nutrient transport [*Slc16a9* (carnitine), *Trf* (iron), *Slc38a4* (cationic and neutral amino acids)] (**Fig. 3A**, **table S5E**). aSI-L non-proliferating decidual cells had DEGs associated with (i) perturbed decidualization (*Igfbp7*, *Tgfbi*, *Lum*, *Gzmg*, *Thbs1*, *Il15ra*), (ii) reduced implantation receptivity (*Adamtsl1*, *Tll1*, *Atp6v0d2*, *Col4a3*), (iii) increased tissue repair (*Ccn3*, *Il20ra*) and (iv) increased tumor suppressor signaling (*Meg3*, *Opcml*, *H19*) (**Fig. 3B**, **table S5E**).

**Figure 3.**
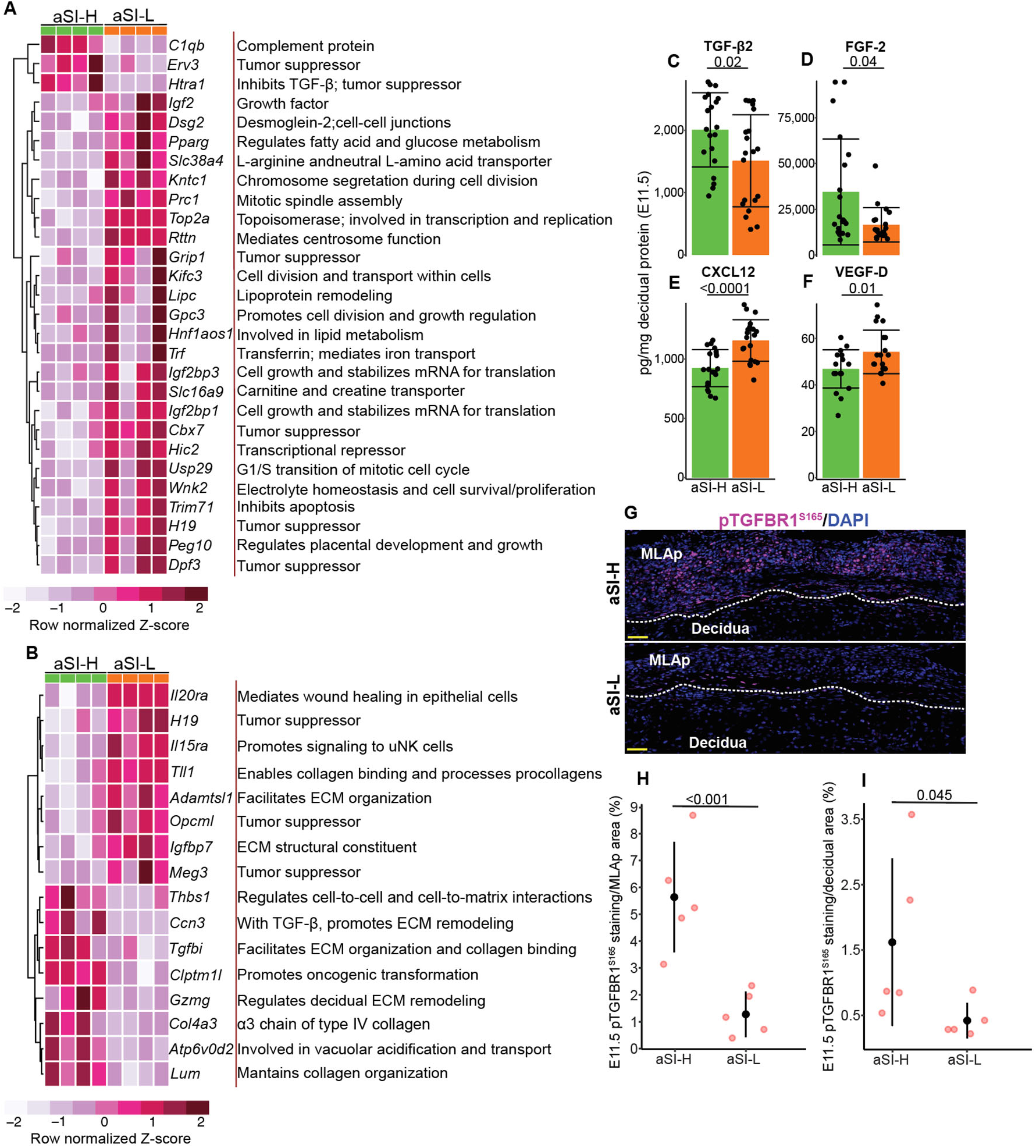
aSI-L deciduas and MLAps have a decrease in TGF-β signaling compared to aSI-H dams. DEGs identified when comparing all aSI-L to aSI-H **(A)** proliferating decidual cells (|log_2_(fold-difference)| > 3) and **(B)** non-proliferating decidual cells (|log_2_(fold-difference)| > 1). Quantification of **(C)** TGF-β2, **(D)** FGF-2, **(E)** CXCL12, and **(F)** VEGF-D proteins in decidual/MLAp homogenates at E11.5 (n = 5 dams/group, 3-4 placentas/dam). Mean ± SD shown. Adjusted p-values calculated with a linear mixed model (*Protein level ∼ Microbiota + (1 | Litter ID)*) for pairwise comparisons. **(G)** Representative examples of pTGFBR1^S165^ immunostaining in E11.5 aSI-L and aSI-H deciduas and MLAps. Scale bar = 50 µm. **(H, I)** Proportion of MLAp (panel **H**) and decidua (panel **I**) tissues stained with antibodies to pTGFBR1^S165^ (n = 1 tissue section/placenta, 1-2 placentas/dam, 4 dams per treatment group). Mean ± SD shown. Adjusted p-values defined using a linear mixed model (*%Staining ∼ Microbiota + (1 | Litter ID))* for pairwise comparisons.

Taken together, these transcriptional changes indicate that aSI-L colonization disrupts decidual cell proliferation in addition to ECM production and maintenance - changes that have the potential to alter cell-cell comunications in the MFI’s immune and stromal microenvironment. Quantifying the concentrations of proteins involved in decidual regulation of immune, ECM, and angiogenic signaling pathways in E11.5 decidual/MLAp homogenates (see *Methods*) revealed that levels of transforming growth factor beta 2 (TGF-β2) and fibroblast growth factor-2 (FGF-2) were significantly decreased, whereas levels of CXCL12 [also known as stromal cell-derived factor 1 (SDF-1)] and vascular endothelial growth factor-D (VEGF-D) were significantly increased in aSI-L compared to aSI-H dams (n = 4 MFIs/dam, 5 litters/group; p < 0.05, linear mixed model) (**Fig. 3C-F**). These results suggest that the EED donor-derived and healthy donor-derived bacterial consortia differentially affect the decidual immune-stromal signaling axis – a hypothesis that we explored further with spatial transcriptomics.

### Reduced TGF-β and FGF-2 signaling from NK to decidual cells in aSI-L dams

To visualize the cell clusters profiled in our snRNA-seq experiments and to further resolve their relative positions in the maternal and fetal components of the MFI, we utilized a mouse 5,000 gene panel and an *in situ*-based method to detect and quantify these transcripts. This approach allowed us to perform cell segmentation to determine which cells were expressing specified genes (see *Methods*). The marker genes used to classify the NK cells into two clusters (cNK and uNK) and the decidual cells into three clusters (types 1, 2, and 3) are described in **Supplementary Results** (**Fig. 4A-C**, **fig. S12C**).

**Figure 4.**
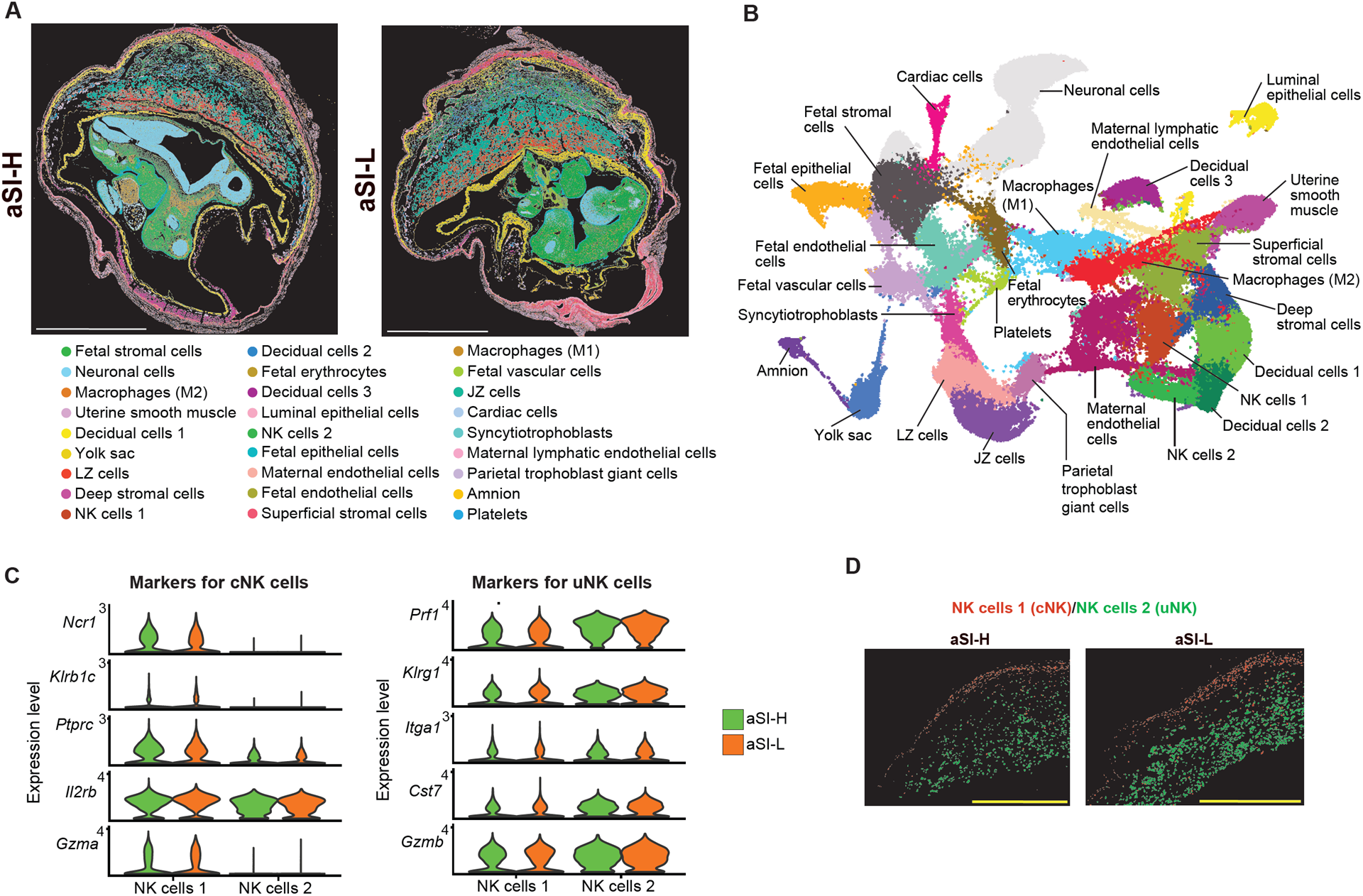
Spatial transcriptomics illustrates spatially segregated populations of NK cells with high and low expression of both *Ncr1* and *Klrb1c* in aSI-L and aSI-H deciduas and MLAp. (A) Representative Xenium cell segmentation of aSI-L and aSI-H E11.5 whole conceptuses with cluster identifications. Colors representing the clusters are listed underneath. Scale bar = 2000 µm. **(B)** UMAP representation of 805,053 cells across sections from six E11.5 whole conceptuses (n=1 section/conceptus; n= 3 aSI-L, 3 aSI-H conceptuses); cells were assigned to 27 clusters. **(C)** Expression of uNK-and cNK-associated genes in aSI-L and aSI-H NK cell type 1 and type 2 clusters. **(D)** Representative Xenium segmentation for type 1 and type 2 NK cell clusters in aSI-L and aSI-H samples. Scale bar = 1000 µm.

The proximity of the uNK and cNK cells to type 1 and 2 decidual cells (**Fig. 5A**) prompted us to apply the MultiNicheNet algorithm to our spatial transcriptomic dataset to investigate bidirectional NK cell-decidual cell communications in aSI-L versus aSI-H MFIs. MultiNicheNet infers differentially expressed ligand-receptor interactions: it predicts potential interactions by cell type and considers differential expression of genes known to operate downstream of a receptor’s activation by its known ligands (*38*).

**Figure 5.**
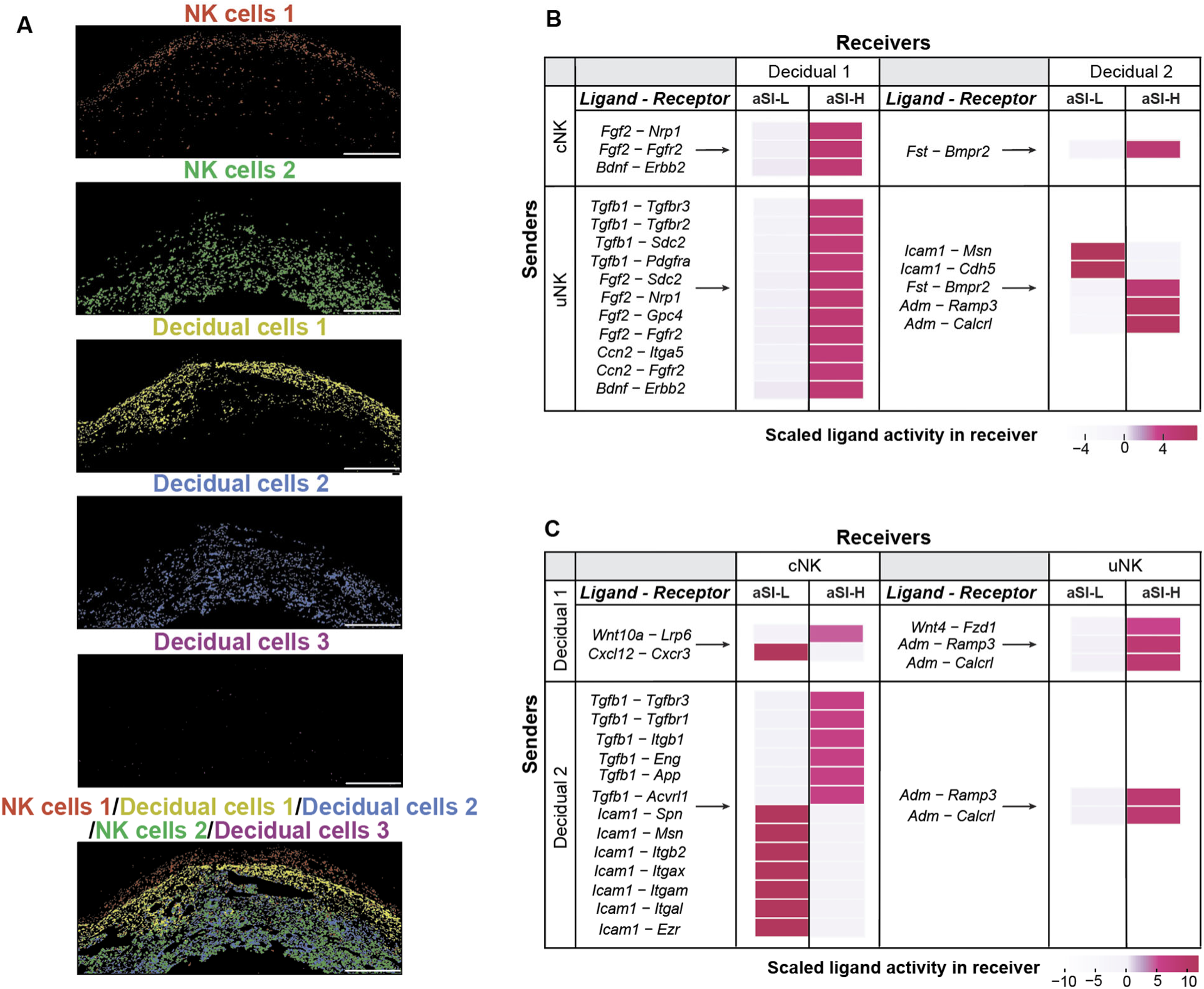
MultiNicheNet analysis of spatial transcriptomic table identifies perturbations in TGF-β, FGF-2, and Wnt signaling pathways mediating communication between E11.5 aSI-L NK cells and decidual stromal cells. **(A)** Representative Xenium cell segmentation of NK cell and decidual stromal cell clusters at the mesometrial side of an E11.5 aSI-L conceptus. Scale bar = 500 µm. (**B,C**) MultiNicheNet analysis of scaled (*i.e.* z-score normalization across all sender-receiver cell type combinations) activity for NK cell clusters as “senders” and decidual cell clusters as “receivers” (panel **B**) and decidual cell clusters as “senders” and NK cell clusters as “receivers” (panel **C**) (n = 1 conceptus from 3 dams/treatment group; aSI-H versus aSI-L contrast used; ligand-receptor pairs associated with the top 20 scaled ligand activities are shown).

We first designated the two NK cell clusters as ligand expressing “senders” and decidual cells as “receivers” (*i.e.* receptor expressing populations). Compared to dams harboring the aSI-H consortium, aSI-L cNK ‘senders’ exhibited significant decreases in *Fgf2* signaling through neuropilin 1 (*Nrp1*) and fibroblast growth factor receptor 2 (*Fgfr2*) in type 1 decidual cell ‘receivers’ (**Fig. 5B**, **table S6C**), consistent with reduced FGF2 levels in aSI-L versus aSI-H E11.5 decidual/MLAp homogenates (**Fig. 3D**). FGF-2 is normally expressed at high levels in the mesometrial decidua, where it promotes proliferation of stromal cells, in addition to the subsequent remodeling of the decidual ECM and vasculature (*39*, *40*). There was also a decrease in *Fgf2* ligand-mediated communication between uNK cell senders and type 1 decidual cell receivers, with decreased expression of type 1 decidual cell “receiver” genes related to ECM-and cell-cell adhesion-related receptors [syndecan-2 (*Sdc2*), glypican-4 (*Gpc4*), *Nrp1*, and *Fgfr2*] (**Fig. 5B**, **table S6C**). Immunohistochemistry (IHC) confirmed that there was less FGFR2 phosphorylated at its major activation site (pFGFR2^Y769^) (*41*) in E11.5 aSI-L compared to aSI-H deciduas/MLAps (**fig. S12D**). Hence, by E11.5, colonization with the aSI-L consortium reduces NK cell FGF-2 signaling to immune-associated type 1 decidual cells, potentially contributing to stromal cell dysfunction at the MFI of aSI-L mice.

TGF-β signaling is essential for decidual cell development and function; the level of this protein was significantly reduced in aSI-L versus aSI-H decidual/MLAp homogenates (**Fig. 3C**). MultiNicheNet analysis showed that when compared to cNK cells, uNK cells were the predominant source of TGF-β ligand activity (**Fig. 5B, table S6C**). Compared to aSI-H uNK cells, aSI-L uNK cells had decreased *Tgfb1* ligand activity with TGF-β family receptors expressed in type 1 decidual cell receivers, such as *Tgfbr2* and co-receptor *Tgfbr3* (**Fig. 5B**, **table S6C**). Moreover, the aSI-L uNK cells had reduced *Tgfb1* “sent” to ECM-related receptors, such as *Sdc2* and platelet-derived growth factor receptor alpha (*Pdgfra*), in type 1 decidual cell receivers (**Fig. 5B, table S6C**).

Serine 165 phosphorylation of TGF-β receptor 1 (pTGFBR1^S165^) modulates TGF-β– induced cellular responses and is associated with reduction in TGF-β–mediated growth inhibition and increase in TGF-β–induced apoptosis (*42*). We performed an IHC analysis of levels of pTGFBR1^S165^ in E11.5 aSI-L and aSI-H decidua/MLAp (**Fig. 3G**). Supporting our MultiNicheNet results, aSI-L decidual cells exhibited significantly decreased pTGFBR1^S165^ staining in the MLAp and decidua of aSI-L compared to aSI-H dams (n = 1 MFI from each of 5 litters/treatment group; p < 0.001 and p = 0.045 respectively, Wilcoxon rank-sum test) (**Fig. 3H,I).**

For decidual stromal cells, TGF-β facilitates decidualization (*43*), limits pathological fibrosis (*44*), and stimulates regulated apoptosis or ‘decidual regression’ (*45*). MultiNicheNet analysis indicated a reduction in signaling from uNK to type 1 decidual stromal cells involving connective tissue growth factor (CTGF, encoded by *Ccn2*) in aSI-L dams (**Fig. 5B, table S6C**). As TGF-β and CTGF stimulates extracellular matrix connective protein deposition (*46*, *47*), these results are consistent with the reduced connective tissue staining on Movat pentrochrome staining in E11.5 aSI-L versus aSI-H decidua (**fig. S5E**). In the case of type 2 decidual stromal cells, uNK sender cells in aSI-L mice had reduced ligand activity for *Adm* (inducer of decidual cell immune tolerance) and *Fst* (supports decidualization and decidual stromal cell migration (*48*, *49*)), while there was elevated ligand activity for the adhesion molecule *Icam1* (**Fig. 5B, table S6C**). Together, these results suggest that in aSI-L dams, uNK cells have reduced expression of ligands that promote decidualization.

### Decreased TGF-β and Wnt signaling from aSI-L decidual stromal to NK cells

We next conducted a ‘reciprocal analysis’ by designating members of the decidual cell clusters as ligand-expressing senders and those in NK cell clusters as receptor-expressing receivers (**Fig. 5C**, **table S6D**). At E11.5, aSI-L type 1 decidual senders exhibited decreased Wnt (*Wnt4*, *Wnt10a*) ligand signaling involving cNK cell receivers’ Wnt co-receptor (*Lrp6*) and uNK cell receivers’ Wnt receptor (*Fzd1*) (**Fig. 5C**, **table S6D**). Phosphorylation of the co-receptor LRP6 (pLRP6^T1479^) is important for initiating the canonical Wnt-β-catenin signaling pathway (*50*). IHC analyses demonstrated reduced levels of pLRP6^T1479^ in the MLAp of aSI-L dams (**fig. S12E**).

At E11.5, aSI-L type 2 decidual cell senders had decreased *Tgfb1* ligand signaling to cNK cell receivers; as TGF-β contributes to conversion of cNK to uNK cells (*51*, *52*), this is consistent with our flow cytometry findings of reduced uNK cell proportions in aSI-L versus aSI-H E11.5 decidua/MLAp (**Fig. 2G**). Additionally, aSI-L type 1 and 2 decidual senders had decreased adrenomedullin (*Adm*) signaling to uNK cells (**Fig. 5C**). Adrenomedullin is associated with multiple roles in pregnancy including the appropriate recruitment and activation of uNK cells (*53*).

aSI-L type 2 decidual cells showed increased *Icam1* ligand signaling involving cNK receivers’ CD43 (*Spn*), CD18 (*Itgb2*), CD11b (*Itgam*), CD11c (*Itgax*), lymphocyte function-associated antigen 1 (LFA-1 [CD11a]) (*Itgal*), moesin (*Msn*), and ezrin (*Ezr*) (**Fig. 5C**, **table S6D**). ICAM-1 (intracellular adhesion molecule 1) is a cell surface receptor expressed in human decidual stromal and endothelial cells that helps recruit cNK cells into the decidua (*54*, *55*). There was also an increase in aSI-L type 1 decidual cell *Cx3cl1* signaling to uNK integrin subunits (*Itgb1*, *Itgb3*, *Itgav*) (**table S6D**). Progesterone, also elevated in aSI-L decidua (**fig. S11A**), supports chemokine expression including CX3CL1 in decidual cells, which in turn may support NK cell migration and/or retention at the decidua (*56*).

Collectively, these reciprocal sender–receiver MultiNicheNet analyses demonstrate that compared to the consortium of cultured bacteria derived from the duodenal microbiota of healthy women, the cultured consortium from women with EED alters bidirectional communication between NK cell and decidual cell clusters. This reshaping is characterized by reductions in NK-derived FGF-2 and TGF-β signaling to decidual cells and, conversely, diminished decidual Wnt and TGF-β signaling to NK cells. These shifts in signaling are reflected in the differences in NK cell composition and transcriptomic evidence of decidual cell proliferation in aSI-L vs. aSII-H dams. Our findings raised the question of whether these perturbations in aSI-L dams can be ‘repaired’ by members of the aSI-H consortium.

### Co-housing experiments

Cohousing of coprophagic mice facilitates microbial mixing, in which bacterial strain transmission and associated changes in host phenotypes can be assessed. The experimental design, shown in **Fig. 6A**, relied on transmission of the aSI-L and aSI-H consortia from previously-colonized aSI-L and aSI-H females to GF recipients. This approach allowed us to establish lines of aSI-L and aSI-H mice that propagated their respective microbial communities. After 28 days of colonization, we co-housed aSI-L and aSI-H females for another 28 days before timed matings. The resulting pregnant mice yielded co-housed lines that had originally been colonized with either the aSI-L or the aSI-H bacterial consortium (labeled CoH-L and CoH-H in **Fig. 6A**). There were no significant differences in fetal (p = 0.94) or placental (p = 0.29) weights between the aSI-L and CoH-L groups at E11.5 (n = 46 fetuses representing 4 dams/treatment group, linear mixed model with Benjamini-Hochberg (BH) FDR correction, **fig. S13A**,**B**). However, bulk RNA-seq of placental tissue (fetal tissue with some residual decidua) revealed significant transcriptomic remodeling and “repair” in CoH-L compared to aSI-L dams. Genes that were significantly differentially expressed in CoH-L compared to non-cohoused aSI-L dams were indicative of (i) decreased ECM adhesion (*Dpt*, *Cuzd1*, *Fras1*) and collagen production (*Prr33*, *Col4a6*), (ii) decreased innate immunity (*C3*, *Pzp*, *Clec2h*, *C6*), (iii) decreased cellular proliferation (*Rarres1*, *Bnc1*, *Fgf1*, *Tubal3*, *Fgfr4*, *Rspo1*), (iv) decreased cellular stress (*Hpx*, *Fgf1*), and (v) decreased steroid hormone production (*Hsd3b1*) (**|**log_2_(fold-difference)| > 1.5, FDR < 0.05, Wald test) (**fig. S13C**, **table S7A**). Gene set enrichment analysis (GSEA) of Gene Ontology Biological Process terms (GOBP) showed that transcripts in E11.5 CoH-L fetal placentas were enriched for processes related to innate immune responses, ribosomal biogenesis, and translation, whereas expression of steroid metabolic processes was diminished compared to non-cohoused aSI-L animals (**table S7B**).

**Figure 6.**
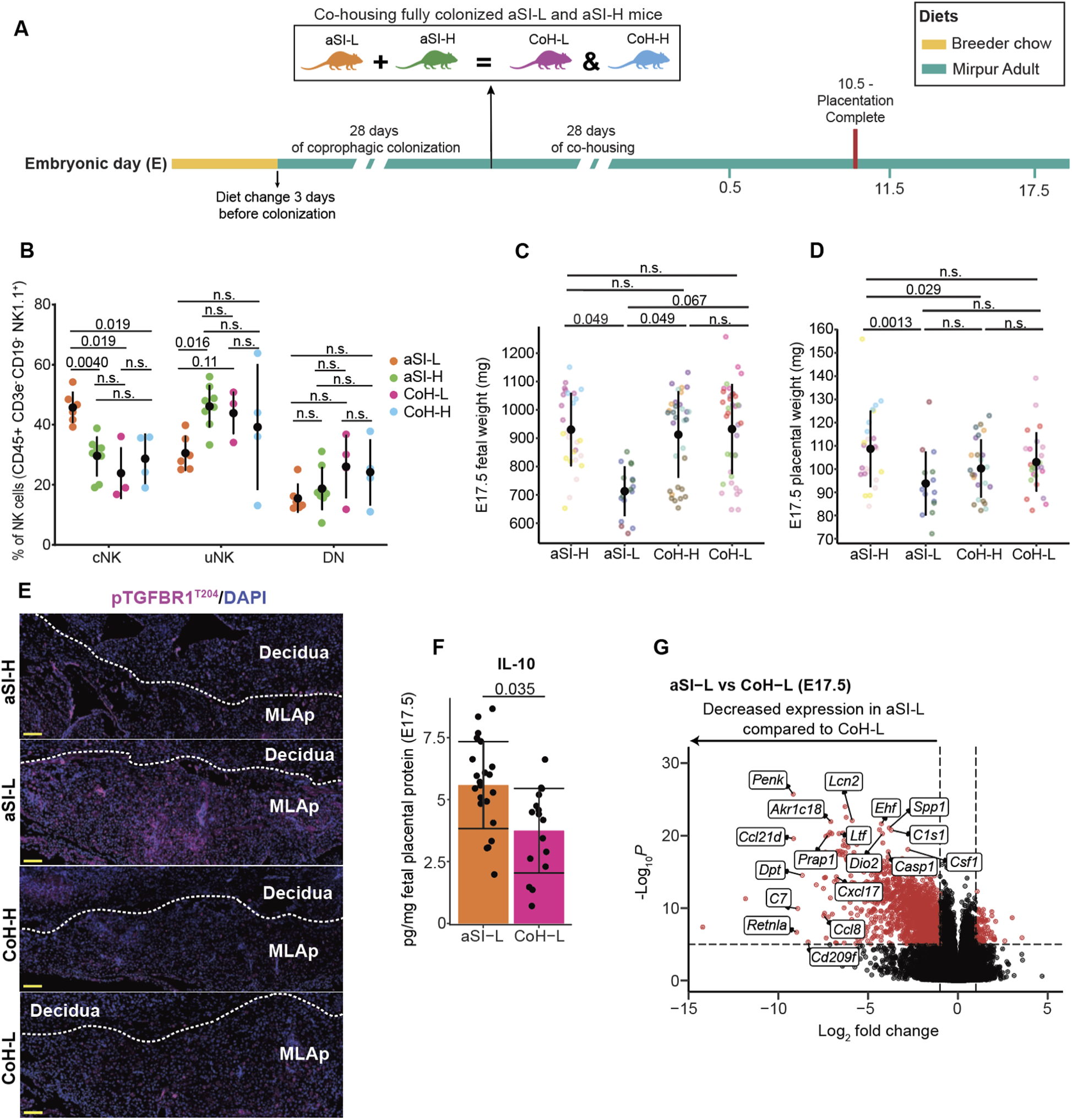
Effects of co-housing aSI-L and aSI-H mice on fetal weights and whole placental immune responses at E17.5. **(A)** Experimental design. (**B**) Quantification of cNK, uNK, and DN NK cell proportions in E11.5 deciduas (n = 4-8 dams/treatment group; Mann-Whitney-Wilcoxon test, BH correction). All deciduas are pooled across a litter. Mean ± SD are shown. **(C)** Fetal weights [aSI-L: 3 dams, 17 fetuses, 712.9 ± 88.0 mg (mean ± SD); aSI-H: 4 dams, 28 fetuses, 930 ± 129.6 mg; CoH-L: 5 dams, 37 fetuses, 932 ± 159.1 mg; CoH-H: 5 dams, 32 fetuses, 912.6 ± 152.7 mg]. **(D)** Corresponding placental weights [aSI-L: 93.8 ± 13.8 mg (mean ± SD); aSI-H: 108.7 ± 16.5 mg; CoH-L: 103.1 ± 12.7 mg; CoH-H: 100.2 ± 12.6 mg]. Each litter is represented by a different color. **(E)** Examples of pTGFBR1^T204^ staining in sections of E17.5 aSI-L and aSI-H deciduas and MLAps. Scale bar = 100 µm. **(F)** IL-10 protein quantification from E17.5 aSI-L and CoH-L placental homogenates containing fetal and residual decidual components. Mean values ± SD are shown (n = 4-6 dams/group, 2-4 placentas/dam). Adjusted p-values calculated using a linear mixed model (*IL-10 concentration ∼ Microbiota + (1 | Litter ID)*) for pairwise comparisons. **(G)** Volcano plot of bulk RNA-seq data showing statistically significant immune-related DEGs from E17.5 aSI-L vs CoH-L fetal placentas (n = 4-6 dams/group, 2-5 placentas/dam).

Although co-housing did not improve fetal or placental weights at E11.5 (**fig. S13A**,**B**), flow cytometry demonstrated that NK cell composition in the E11.5 decidua/MLAp was affected. Consistent with our earlier findings (**Fig. 2G**), compared to non-cohoused aSI-H mice, non-cohoused aSI-L dams had a significantly increased proportion of cNK cells (p = 0.0040). Compared to non-cohoused aSI-L controls, cNK cells were significantly decreased in CoH-L animals (p = 0.019) (**Fig. 6B**). These observations indicate that co-housing reduces the population of cNK cells in the MFI of aSI-L mice at this early post-placentation timepoint.

At E17.5, placental and fetal weights in dams belonging to the non-cohoused aSI-L group were significantly lower than in the non-cohoused aSI-H lines (p = 0.0013 and p = 0.049 respectively; n = 124 fetuses representing 4-5 litters/group, linear mixed model with BH correction; **Fig. 6C,D**). Co-housing aSI-L and aSI-H mice before timed matings led to increased fetal weights at E17.5 in co-housed animals compared to non-cohoused aSI-L controls (**Fig. 6C**). While placental weights between E17.5 aSI-L and CoH-L dams were not significantly different (**Fig. 6D**), proteomic and transcriptomic analyses provided evidence that cohousing decreases anti-inflammatory and increases immune responses, potentially repairing the dysregulated maternal immune responses produced by the EED-donor associated bacterial consortium. Specifically, the MLAp region of the MFI exhibited reduced levels of immunocytochemical staining for pTGFBR1^T204^ in CoH-L compared to non-cohoused aSI-L dams (**Fig. 6E**). This phosphorylated form is associated with TGF-β activation (*57*) which in turn promotes immune suppression (*58*) . Luminex assays also revealed a decrease in levels of IL-10, an anti-inflammatory protein (*59*), in E17.5 CoH-L compared to aSI-L animals (n = 4 placentas/dam; 4-5 litters/group; p = 0.035, linear mixed model) (**Fig. 6F**). In addition, bulk RNA-seq disclosed that compared to CoH-L placentas, E17.5 aSI-L placentas had significant decreases in expression of genes associated with (i) the complement cascade (*C7*, *C1s1*), (ii) immune cell recruitment, antimicrobial responses, and inflammation (*Ccl8*, *Ccl21d*, *Lcn2*, *Csf1*, *Cxcl17*, *Ltf*, *Retnla*, *Casp1*, *Cd209f*, *Prap1*), (iii) hormone metabolism (*Dio2*, *Akr1c18*, *Penk*), and (iv) ECM components and wound healing responses (*Ehf*, *Dot*) (**|**log_2_(fold-difference)| > 1, FDR < 0.05, Wald test) (**Fig. 6G**, **table S8A**). GSEA of GOBP terms showed that gene expression in E17.5 CoH-L fetal placentas was enriched for terms related to immune activation, signaling, and differentiation compared to aSI-L animals (**table S8B**).

### Bacterial taxa associated with fetal growth outcomes

There were no bacteria whose absolute abundances were significantly different between mice originally colonized with the aSI-L or aSI-H consortia and subsequently cohoused (p>0.05, Benjamini Hochberg-corrected Kruskal-Wallis with Tukey’s post-hoc tests) (**table S9A,B**). We concluded that cohousing efficiently ‘mixed’ the two consortia and, for the purpose of bacterial abundance analyses, we treated the co-housed animals as one group.

We compared the absolute abundances of bacteria in aSI-L compared to aSI-H and co-housed dams at both E11.5 (**fig. S14A**, **table S9C**) and E17.5 (**fig. S14B**, **table S9D**), first seeking bacteria whose abundances were significantly elevated in aSI-L compared to aSI-H and/or cohoused dams. A total of 11 unique ‘aSI-L–associated species’ satisfied these criteria. *Streptococcus gordonii, Klebsiella granulomatis, Escherichia ruysiae, Escherichia fergusonii, Clostridium isatidis* and *Granulicatella adiacens* were significantly higher in aSI-L dams compared to both aSI-H and co-housed dams; *Faecalibaculum rodentium, Akkermansia muciniphila, Weisella confusa, Cutibacterium acnes* and *Enterococcus lactis* were significantly higher in aSI-L dams compared to either aSI-H or cohoused dams, suggesting detrimental roles in fetal growth (**fig. S14**, **table S9C,D**). Conversely, the absolute abundances of four ‘aSI-H– associated species’, *Streptococcus parasanguinis, Streptococcus anginosus, Veillonella atypica* and *Bifidobacterium pseudolongum* subsp. *globosum* were significantly higher in aSI-H and/or co-housed dams compared to aSI-L dams suggesting beneficial or homeostatic roles in fetal growth (**fig. S14**; **tables S9C,D**),

We previously described the composition of the duodenal microbiota of adult Bangladeshi women with EED and their healthy counterparts from whom the aSI-L and aSI-H bacterial consortia were derived (*18*). Accordingly, we defined the composition of the cecal microbial communities in mice using full-length 16S rRNA amplicon sequencing to enable bacterial strain matching between our pre-clinical model and the duodenal aspirates of women (*18*). The absolute abundances of the ‘aSI-L–associated species’ described above were not significantly different between women with (n=60) and without (n=20) histopathologic evidence of EED (**table S10**). However, the absolute abundance of the ‘aSI-H–associated’ *V. atypica* was significantly higher in the duodenal aspirates of healthy women compared to those with EED (**table S10**), and *V. atypica, S. anginosus,* and *S. parasanguinis* were all predictive features that distinguished healthy controls from women with EED (*18*).

### Bacterial taxa from aSI-L mice correlate with inflammatory plasma proteins in Bangladeshi women

We collected plasma from undernourished women with histopathologic evidence of EED (n=22) compared to healthy controls (n=25) and performed plasma proteomics. Using the aptamer-based platform SomaScan, we quantified 9,664 proteins (10,800 unique aptamers) (*18*). We then performed pairwise alignments between the full-length 16S rRNA sequences of bacteria whose abundances were significantly different between aSI-L versus aSI-H and/or co-housed dams (described in the preceding section) and the full-length 16S rRNA sequences of bacteria present in the duodenal microbiota of these 47 women from whom the plasma proteomic data were acquired. Four aSI-L bacteria negatively implicated in fetal growth (*C. acnes*, *E. ruysiae, G. adiacens, S. gordonii*) and one aSI-H bacterium positively implicated in fetal growth (*V. atypica*) were prevalent enough in the duodenal aspirates of women to perform correlations between absolute bacterial abundance and levels of 6,973 plasma proteins (7,560 unique aptamers) that passed quality and variance thresholds in matched samples (**fig. S15A, table S11A,B**).

A total of 565 unique plasma proteins were significantly positively correlated (p < 0.05, rho > 0) with the absolute abundances of aSI-L bacteria in the women’s duodenal aspirates (966 significant bacterial-protein pairs, **table S11A**). These plasma proteins are involved in cytokine signaling (IL-1, IL-12, IL-18) and recognition and responses to bacteria (Toll Like Receptor signaling cascades, neutrophil degranulation, antigen processing, MAPK signaling cascades) (**fig. S15B-D, table S11C**). Conversely, the abundances of these aSI-L bacteria were significantly negatively correlated with 887 plasma proteins (p < 0.05, rho < 0, 1,299 significant bacterial-protein pairs, **table S11A**). These negatively correlated proteins are involved in regulation of hormone secretion, metabolism (prostaglandin metabolism, fatty acid biosynthesis, response to nutrient levels, insulin secretion), and vasculogenesis (vascular-associated smooth muscle cell proliferation) (**table S11C**).

We next sought to determine whether the plasma proteins that were significantly positively correlated with the absolute abundances of aSI-L bacteria in duodenal aspirates were also related to activities at the MFI in our pre-clinical mouse model. To do so, we queried our decidual tissue bulk RNA-Seq dataset from the co-housing experiment for genes that were significantly more highly expressed in non-cohoused aSI-L dams compared to co-housed dams originally colonized with the aSI-L consortium (CoH-L). The results revealed 108 plasma proteins that were (i) positively correlated with the abundances of duodenal taxa represented by the aSI-L bacteria and (ii) matched mouse homologues whose expression in the decidua was significantly higher in aSI-L compared to co-housed dams (**table S11D**). Conversely, 266 genes in the decidua were significantly more highly expressed in co-housed (CoH-L) compared to aSI-L dams and were homologs of plasma proteins that were significantly negatively correlated with the absolute abundances of aSI-L bacteria (**table S11E**).

Mouse homologues to positively-correlated plasma proteins included: (i) Arginosuccinate synthase 1 (Ass1), which provides the substrate for nitric oxide synthesis, ensuring proper uterine placental blood flow, (ii) Glutathione S-transferase alpha 5 (Gsta5), which protects against lipid perioxidation products that accumulate due to oxidative stress and may impair placental function, (iii) Tricellulin (Marveld2), a tight junction protein that maintains synctiotrophoblast barrier integrity, (iv) Transformation related protein 53 inducible protein 11 (Trp53i11), which mediates programmed cell death, (v) C-type lectin domain family 1 member B (Clec1b), required for the blood-lymph barrier at the placental interface, (vi) C-Maf-inducing protein (Cmip), which promotes Th2 immune polarization, and (vii) Numb, a primary antagonist of Notch signaling that mediates cell proliferation and migration in the placenta (**table S11D**).

The taxa most strongly enriched (largest magnitude of difference) in non-cohoused aSI-L compared to non-cohoused aSI-H and co-housed dams were *G. adiacens* and *S. gordonii* (**table S9C,D**). These two organisms are typically associated with the oral habitat in humans (*60*). Their absolute abundances in the duodenal microbiota of women were significantly correlated with plasma proteins involved in inflammatory activation, tissue repair, hormonal regulation and metabolic perturbation (**fig. S15, tables S11A,C**).

Together, these analyses highlight proteins that were altered at the MFI by manipulation of the maternal gut microbiota in our pre-clinical model and significantly correlated with the abundances of these bacteria in plasma of women from whom the aSI-L and aSI-H bacterial consortia derived. They also identify SI bacteria that are candidate therapeutic targets for treating or preventing IUGR and its associated developmental impairments.

## DISCUSSION

The present study illustrates one approach for determining whether and how the SI microbiota is related to impaired prenatal development in undernourished women with EED. The approach consists of (i) initial observational studies in a population where the burden of maternal undernutrition and EED is great and where both affected and healthy individuals can be studied, (ii) ‘reverse translation’ to a representative preclinical model, in which tests of a causal relationship between microbial community composition and prenatal development can be performed and putative mechanisms identified and (iii) contextualization of the results obtained in (ii) by referencing to the human study that provided the microbial communities incorporated into the preclinical model so that therapeutic targets (and biomarkers) can nominated and prioritized. Specifically, we have colonized pregnant gnotobiotic mice fed a diet representative of that consumed by the study population living in Dhaka, Bangladesh, with a consortium of bacteria cultured from the duodenal microbiota of undernourished Bangladeshi women with EED or cultured from healthy controls. Our comparative analysis indicated that the largest perturbation produced by the bacterial consortium from undernourished women with EED (aSI-L) is on the maternal-derived compartment of the maternal fetal interface, including the decidual uNK cells and stromal cells, midway through gestation (E11.5) rather than closer to parturition (E17.5). This finding contrasts with our previous study in GF, CONV-R, and CONV-D animals, in which differences were predominantly observed within the fetal placental compartment of the MFI (*14*). In that study, GF mice exhibited reduced angiogenesis localized to the fetal LZ and JZ, mediated by disruption of VEGF-A signaling through the VEGFR2 cascade. In the current study, colonization with an EED-associated microbiota did not markedly alter placental angiogenesis or vascular architecture. Instead, aSI-L colonization preferentially impacted the immune and stromal compartments of the decidua.

The most prominent immune phenotype at E11.5 involved maternal uNK cells. This tissue-resident NK cell type arises from peripheral cNK cells recruited into the uterus to aid in vascular and decidual remodeling (*61–63*). Flow cytometry of E11.5 decidua and MLAp tissue revealed a higher proportion of cNK cells in aSI-L mice. There was also an increase in E11.5 aSI-L maternal endothelial cells shown by fetal placental snRNA-seq and *Pecam1* FISH decidual staining. While uNK cells play a critical role in the remodeling of the maternal vascular supply, we did not find any significant changes in the decidual vascular morphometrics between the aSI-L and aSI-H mice at E11.5 or E17.5, leading us to conclude that the EED-associated microbial consortium affects the immune function of uNK cells more than their vascular remodeling capacity.

Our spatial transcriptomic results suggested a dynamic communication between NK cells and decidual stromal cells is affected by the aSI-L versus aSI-H microbiota. Comparative MultiNicheNet analysis revealed that aSI-L uNK cells had decreased TGF-β sent to decidual stromal cells; multiNicheNet also disclosed a decrease in TGF-β signaling from postmature decidual cells (type 2 decidual cells) to cNK cells in the aSI-L group. Deficiencies in TGF-β signaling has been found to disrupt multiple aspects of the maternal-fetal interface, including trophoblast organization and cNK-to-uNK conversion, leading to fetal growth restrition (*51*, *52*, *61*, *64*).

Our findings are consistent with the view that bidirectional TGF-β signaling is required between cNK cells and decidual stromal cells to dampen NK-activating receptor expression and facilitate cNK-to-uNK differentiation (*61*, *65*, *66*). In the absence of TGF-β, cNK cells convert inefficiently, retain elevated expression of activating receptors including NK1.1 and NKp46, and consequently exhibit heightened activation and eventual overactivation. The aSI-L uNK cellular phenotype points towards this hyporesponsive transcriptomic signature, with increased expression of genes involved in activation and anti-apoptotic pathways. Collectively, our findings suggest that EED-associated SI community contribute to reduced cNK-to-uNK conversion and heightened activation of unconverted cNK cells within the decidua and MLAp.

Co-housing aSI-L mice with aSI-H mice (*i.e.* CoH-L) before pregnancy resulted in improved fetal weights compared to non-cohoused aSI-L controls, as well as reduced levels of staining for activated pTGFBR1^T204^, IL-10, and expression of genes involved in immune cell recruitment, inflammation, and ECM wound healing responses. We have identified several bacterial strains elevated in co-housed animals and aSI-H non-cohoused controls at E11.5 and E17.5 compared to aSI-L dams. Strains associated with healthy fetal growth in our pre-clinical model included members of *Veillonella*, *Streptococcus*, and *Bifidobacterium*. Of these, *Veillonella* and *Streptococcus* species were identified as predictors that distinguished healthy women from women with EED.

*Limitations* - The path forward for further delineating the contributions of an EED microbiota to NK-decidual stromal cell cross-talk and IUGR will be challenging. There are combinatorial challenges to sorting out whether all or a subset of the candidate bacterial effectors nominated act directly or in concert with other members of the bacterial consortia, but whose abundances did not change with co-housing. There is the challenge of deciphering whether the actual effectors are directly produced by bacteria, originate in bacteria while requiring modification by the host, or are host-derived (*e.g*. inflammatory immune cell trafficking from the intestine to the decidua, or cytokines produced by intestinal cells that affect decidual immune cells). The complexity of the challenge is compounded by the fact that <10% of the ASVs detected in duodenal aspirates obtained from women in the study cohorts who underwent EGD were represented in our culture collection (*18*). Nonetheless, those strains from women with EED that did colonize our gnotobiotic mouse model were sufficient to produce enteropathy and IUGR, whereas those from healthy women were sufficient to ameliorate IUGR. As such, these findings provide a rationale for further advancing efforts to develop approaches for repairing the SI microbial community of undernourished women of childbearing age with EED prior to pregnancy, with the goal of improving birth outcomes and promoting healthy postnatal development of their offspring.

## Acknowledgments

We thank David O’Donnell, Maria Karlsson and the late Justin Serugo for their invaluable assistance with mouse husbandry, Martin Meier for generating libraries for bulk RNA-seq and 16S rRNA amplicon sequencing, Jessica Hoisington Lopez, MariaLynn Crosby plus members of the Genome Technology Access Core at Washington University School of Medicine for sequencing the libraries, Kymberli May and the Washington University Digestive Diseases Research Cores Center Tissue Analysis & Imaging Core (supported by NIH grant P30 DK052574) and Crystal Idleburg and Samantha Coleman Cathcart from the Washington University Musculoskeletal Research Center Histology and Morphometry Core (supported by NIH grant P30 AR074992) for tissue embedding and sectioning, and Krzysztof Hyrc from the Hope Center Alafi Neuroimaging Laboratory (supported by NIH grants S10 RR027552 and S10 OD032121) at Washington University School of Medicine for imaging and scanning services. Finally, we thank members of the Gordon and Diamond laboratories for reagents and their very valuable suggestions.

## Funding

National Institutes of Health grant DK131107 (JIG)

Gates Foundation INV-033564 (JIG)

National Institutes of Health grant F30HD115307 (RC)

National Institutes of Health grant T32DK007130 (ZC)

AGA Research Foundation’s AGA-Ironwood Fellowship-to-Faculty Transition Award AGA2024-32-02 (ZC)

Helen Hay Whitney Foundation fellowship (KMP)

## Author contributions

Conceptualization: RC, ZLC, KMP, JIG

Methodology: RC, ZLC, KMP, MSD, JIG

Investigation: RC, ZLC, KMP, HL, AMR, EML

Visualization: RC, ZLC, KMP

Funding acquisition: JIG, TA

Project administration: JIG

Supervision: JIG

Writing – original draft: RC, ZLC, KMP, JIG

Writing – review & editing: RC, ZLC, KMP, MJB, MSD, JIG

## Competing interests

MSD is a consultant or advisor for Inbios, Vir Biotechnology, IntegerBio, Moderna, Merck, and GlaxoSmithKline. The Diamond laboratory has received unrelated funding support in sponsored research agreements from Vir Biotechnology, Moderna, and IntegerBio.

## Data, code, and materials availability

Shotgun DNA sequencing, bulk RNA-seq, and snRNA-seq tables have been deposited in NCBI’s Sequence Read Archive (SRA) under project number PRJNA1406798. This study did not use any original code. Additional information required to reanalyze the data reported in this paper is available from the lead contact upon request.

## Supplementary Materials

### Materials and Methods

#### Supplementary Results

Figs. S1 to S15

Tables S1 to S14

References (*67–108*)

## Notes

### Summary of Updates

Reference numbering and formatting were corrected.

## References

1. World Health Organization||United Nations Children’s Fund (UNICEF)||International Bank for Reconstruction and Development/The World Bank. “Levels and trends in child malnutrition: UNICEF / WHO / World Bank Group joint child malnutrition estimates: key findings of the 2025 edition” (Collection 10665/26724, World Health Organization, 2025); https://iris.who.int/bitstream/handle/10665/381846/9789240112308-eng.pdf.

2. A. Kirolos, M. Goyheneix, M. Kalmus Eliasz, M. Chisala, S. Lissauer, M. Gladstone, M. Kerac, Neurodevelopmental, cognitive, behavioural and mental health impairments following childhood malnutrition: a systematic review. BMJ Glob Health 7, e009330 (2022).

3. K. G. Dewey, K. Begum, Long-term consequences of stunting in early life. Matern Child Nutr 7 Suppl 3, 5–18 (2011).

4. E. Ozaltin, K. Hill, S. V. Subramanian, Association of maternal stature with offspring mortality, underweight, and stunting in low-to middle-income countries. JAMA 303, 1507–1516 (2010).

5. R. Gul, S. Iqbal, Z. Anwar, S. G. Ahdi, S. H. Ali, S. Pirzada, Pre-pregnancy maternal BMI as predictor of neonatal birth weight. PLoS One 15, e0240748 (2020).

6. P. Christian, E. R. Smith, A. Zaidi, Addressing inequities in the global burden of maternal undernutrition: the role of targeting. BMJ Glob Health 5, e002186 (2020).

7. A. J. Prendergast, J. H. Humphrey, The stunting syndrome in developing countries. Paediatr Int Child Health 34, 250–265 (2014).

8. R. Martorell, Improved nutrition in the first 1000 days and adult human capital and health. Am J Hum Biol 29 (2017).

9. T. Ahmed, M. Hossain, K. I. Sanin, Global burden of maternal and child undernutrition and micronutrient deficiencies. Ann Nutr Metab 61 Suppl 1, 8–17 (2012).

10. V. Owino, T. Ahmed, M. Freemark, P. Kelly, A. Loy, M. Manary, C. Loechl, Environmental Enteric Dysfunction and Growth Failure/Stunting in Global Child Health. Pediatrics 138, e20160641 (2016).

11. M. S. Hossain, S. M. K. N. Begum, M. M. Rahman, M. Parvez, R. N. Mazumder, S. A. Sarker, M. M. Hasan, S. M. Fahim, M. A. Gazi, S. Das, M. Mahfuz, T. Ahmed, Environmental enteric dysfunction and small intestinal histomorphology of stunted children in Bangladesh. PLoS Negl Trop Dis 17, e0010472 (2023).

12. K. Watanabe, W. A. Petri, Environmental Enteropathy: Elusive but Significant Subclinical Abnormalities in Developing Countries. EBioMedicine 10, 25–32 (2016).

13. M. S. Hossain, S. M. K. N. Begum, M. M. Rahman, R. N. Mazumder, M. Parvez, M. A. Gazi, M. M. Hasan, S. M. Fahim, S. Das, M. Mahfuz, S. A. Sarker, T. Ahmed, Alterations in the histological features of the intestinal mucosa in malnourished adults of Bangladesh. Sci Rep 11, 2355 (2021).

14. R. Coskun, Z. L. Chang, A. Marcial Rodríguez, H. Liu, J. Cheng, Y. Alippe, M. S. Diamond, J. I. Gordon, Effects of the gut microbiota on placental angiogenesis and intrauterine growth in gnotobiotic mice. Proc Natl Acad Sci U S A 122, e2426341122 (2025).

15. M. M. Faas, Y. Liu, T. Borghuis, C. A. van Loo-Bouwman, H. Harmsen, P. de Vos, Microbiota Induced Changes in the Immune Response in Pregnant Mice. Front Immunol 10, 2976 (2019).

16. J. Lopez-Tello, Z. Schofield, R. Kiu, M. J. Dalby, D. van Sinderen, G. Le Gall, A. N. Sferruzzi-Perri, L. J. Hall, Maternal gut microbiota Bifidobacterium promotes placental morphogenesis, nutrient transport and fetal growth in mice. Cell Mol Life Sci 79, 386 (2022).

17. G. N. Pronovost, K. B. Yu, E. J. L. Coley-O’Rourke, S. S. Telang, A. S. Chen, H. E. Vuong, D. W. Williams, A. Chandra, T. K. Rendon, J. Paramo, R. H. Kim, E. Y. Hsiao, The maternal microbiome promotes placental development in mice. Sci Adv 9, eadk1887 (2023).

18. K. M. Pruss, Z. L. Chang, Md. S. Hossain, M. M. Rahman, M. Mahfuz, R. Coskun, R. Sharmin, A. Rezwan, S. A. Sarker, S. Das, S. M. Fahim, Md. A. Gazi, K. A. Hudson, A. M. Rodriguez, H. Liu, R. Kitchen, A. E. Byrne, C. Kao, B. Brodrick, A. Rose, B. Bhattarai, D. Khantakova, J. Fachi, M. Colonna, T. Ahmed, M. J. Barratt, J. I. Gordon, Functional characterization of duodenal microbiota and associated enteropathy in undernourished Bangladeshi women and gnotobiotic mice. medRxiv [Preprint] (2026). 10.64898/2026.08.14.26360472.

19. B. A. Croy, A. Yamada, F. DeMayo, S. L. Adamson, The Guide to Investigation of Mouse Pregnancy (Academic Press, 2014).

20. C. F. Holinka, Y. C. Tseng, C. E. Finch, Prolonged gestation, elevated preparturitional plasma progesterone and reproductive aging in C57BL/6J mice. Biol Reprod 19, 807–816 (1978).

21. M. E. Wilson, S. P. Ford, Comparative aspects of placental efficiency. Reprod Suppl 58, 223–232 (2001).

22. A. Schumacher, S.-D. Costa, A. C. Zenclussen, Endocrine factors modulating immune responses in pregnancy. Front Immunol 5, 196 (2014).

23. M. Hemberger, C. W. Hanna, W. Dean, Mechanisms of early placental development in mouse and humans. Nat Rev Genet 21, 27–43 (2020).

24. L. Woods, V. Perez-Garcia, M. Hemberger, Regulation of Placental Development and Its Impact on Fetal Growth-New Insights From Mouse Models. Front Endocrinol (Lausanne) 9, 570 (2018).

25. M. J. Soares, The prolactin and growth hormone families: pregnancy-specific hormones/cytokines at the maternal-fetal interface. Reprod Biol Endocrinol 2, 51 (2004).

26. J.-P. He, Q. Tian, Q.-Y. Zhu, J.-L. Liu, Identification of Intercellular Crosstalk between Decidual Cells and Niche Cells in Mice. Int J Mol Sci 22, 7696 (2021).

27. G. Kaur, C. B. M. Porter, O. Ashenberg, J. Lee, S. J. Riesenfeld, M. Hofree, M. Aggelakopoulou, A. Subramanian, S. B. Kuttikkatte, K. E. Attfield, C. A. E. Desel, J. L. Davies, H. G. Evans, I. Avraham-Davidi, L. T. Nguyen, D. A. Dionne, A. E. Neumann, L. T. Jensen, T. R. Barber, E. Soilleux, M. Carrington, G. McVean, O. Rozenblatt-Rosen, A. Regev, L. Fugger, Mouse fetal growth restriction through parental and fetal immune gene variation and intercellular communications cascade. Nat Commun 13, 4398 (2022).

28. G. H. Ran, Y. Q. Lin, L. Tian, T. Zhang, D. M. Yan, J. H. Yu, Y. C. Deng, Natural killer cell homing and trafficking in tissues and tumors: from biology to application. Signal Transduct Target Ther 7, 205 (2022).

29. I. Bank, M. Book, R. Ware, Functional role of VLA-1 (CD49A) in adhesion, cation-dependent spreading, and activation of cultured human T lymphocytes. Cell Immunol 156, 424–437 (1994).

30. F. Wang, A. E. Qualls, L. Marques-Fernandez, F. Colucci, Biology and pathology of the uterine microenvironment and its natural killer cells. Cell Mol Immunol 18, 2101–2113 (2021).

31. B. Wang, J. Zhou, Y. Chen, H. Wei, R. Sun, Z. Tian, H. Peng, A novel spleen-resident immature NK cell subset and its maturation in a T-bet-dependent manner. J Autoimmun 105, 102307 (2019).

32. L. Chiossone, J. Chaix, N. Fuseri, C. Roth, E. Vivier, T. Walzer, Maturation of mouse NK cells is a 4-stage developmental program. Blood 113, 5488–5496 (2009).

33. K. M. Pruss, C. Kao, A. E. Byrne, R. Y. Chen, B. Di Luccia, L. Karvelyte, R. Coskun, M. Lemieux, K. Nepal, D. M. Webber, M. C. Hibberd, Y. Wang, H. Liu, D. A. Rodionov, A. L. Osterman, M. Colonna, C. Maueroder, K. Ravichandran, M. J. Barratt, T. Ahmed, J. I. Gordon, Enteropathy produced in mice by intergenerational transmission of small intestinal microbiota from undernourished children. Nat Microbiol, doi: 10.1038/s41564-026-02394-4 (2026).

34. D. I. Campbell, S. H. Murch, M. Elia, P. B. Sullivan, M. S. Sanyang, B. Jobarteh, P. G. Lunn, Chronic T cell-mediated enteropathy in rural west African children: relationship with nutritional status and small bowel function. Pediatr Res 54, 306–311 (2003).

35. P. Yang, L. Liu, L. Sun, P. Fang, N. Snyder, J. Saredy, Y. Ji, W. Shen, X. Qin, Q. Wu, X. Yang, H. Wang, Immunological Feature and Transcriptional Signaling of Ly6C Monocyte Subsets From Transcriptome Analysis in Control and Hyperhomocysteinemic Mice. Front Immunol 12, 632333 (2021).

36. N. V. Serbina, T. Jia, T. M. Hohl, E. G. Pamer, Monocyte-mediated defense against microbial pathogens. Annu Rev Immunol 26, 421–452 (2008).

37. B. K. Tatematsu, D. K. Sojka, Tissue-resident natural killer cells derived from conventional natural killer cells are regulated by progesterone in the uterus. Mucosal Immunol 18, 390– 401 (2025).

38. R. Browaeys, J. Gilis, C. Sang-Aram, P. D. Bleser, L. Hoste, S. Tavernier, D. Lambrechts, R. Seurinck, Y. Saeys, MultiNicheNet: a flexible framework for differential cell-cell communication analysis from multi-sample multi-condition single-cell transcriptomics data. bioRxiv [Preprint] (2023). 10.1101/2023.06.13.544751.

39. R. K. Srivastava, Y. Gu, S. Ayloo, M. Zilberstein, G. Gibori, Developmental expression and regulation of basic fibroblast growth factor and vascular endothelial growth factor in rat decidua and in a decidual cell line. J Mol Endocrinol 21, 355–362 (1998).

40. B. C. Paria, W. Ma, J. Tan, S. Raja, S. K. Das, S. K. Dey, B. L. Hogan, Cellular and molecular responses of the uterus to embryo implantation can be elicited by locally applied growth factors. Proc Natl Acad Sci U S A 98, 1047–1052 (2001).

41. M. G. Gartside, H. Chen, O. A. Ibrahimi, S. A. Byron, A. V. Curtis, C. L. Wellens, A. Bengston, L. M. Yudt, A. V. Eliseenkova, J. Ma, J. A. Curtin, P. Hyder, U. L. Harper, E. Riedesel, G. J. Mann, J. M. Trent, B. C. Bastian, P. S. Meltzer, M. Mohammadi, P. M. Pollock, Loss-of-function fibroblast growth factor receptor-2 mutations in melanoma. Mol Cancer Res 7, 41–54 (2009).

42. S. Souchelnytskyi, P. ten Dijke, K. Miyazono, C. H. Heldin, Phosphorylation of Ser165 in TGF-beta type I receptor modulates TGF-beta1-induced cellular responses. EMBO J 15, 6231–6240 (1996).

43. N. Ni, Q. Li, TGFβ superfamily signaling and uterine decidualization. Reprod Biol Endocrinol 15, 84 (2017).

44. I. Osokine, J. Siewiera, D. Rideaux, S. Ma, T. Tsukui, A. Erlebacher, Gene silencing by EZH2 suppresses TGF-β activity within the decidua to avert pregnancy-adverse wound healing at the maternal-fetal interface. Cell Rep 38, 110329 (2022).

45. B. C. Moulton, Transforming growth factor-beta stimulates endometrial stromal apoptosis in vitro. Endocrinology 134, 1055–1060 (1994).

46. C. Bonnans, J. Chou, Z. Werb, Remodelling the extracellular matrix in development and disease. Nat Rev Mol Cell Biol 15, 786–801 (2014).

47. K. E. Lipson, C. Wong, Y. Teng, S. Spong, CTGF is a central mediator of tissue remodeling and fibrosis and its inhibition can reverse the process of fibrosis. Fibrogenesis Tissue Repair 5, S24 (2012).

48. P. T. Fullerton, D. Monsivais, R. Kommagani, M. M. Matzuk, Follistatin is critical for mouse uterine receptivity and decidualization. Proc Natl Acad Sci U S A 114, E4772– E4781 (2017).

49. G. Liu, Y. Qi, J. Wu, F. Lin, Z. Liu, X. Cui, Follistatin is a crucial chemoattractant for mouse decidualized endometrial stromal cell migration by JNK signalling. J Cell Mol Med 27, 127–140 (2023).

50. W. Pan, S.-C. Choi, H. Wang, Y. Qin, L. Volpicelli-Daley, L. Swan, L. Lucast, C. Khoo, X. Zhang, L. Li, C. S. Abrams, S. Y. Sokol, D. Wu, Wnt3a-mediated formation of phosphatidylinositol 4,5-bisphosphate regulates LRP6 phosphorylation. Science 321, 1350–1353 (2008).

51. J. Peng, D. Monsivais, R. You, H. Zhong, S. A. Pangas, M. M. Matzuk, Uterine activin receptor-like kinase 5 is crucial for blastocyst implantation and placental development. Proc Natl Acad Sci U S A 112, E5098–5107 (2015).

52. M. Eriksson, S. K. Meadows, C. R. Wira, C. L. Sentman, Unique phenotype of human uterine NK cells and their regulation by endogenous TGF-beta. J Leukoc Biol 76, 667–675 (2004).

53. M. Li, N. M. J. Schwerbrock, P. M. Lenhart, K. L. Fritz-Six, M. Kadmiel, K. S. Christine, D. M. Kraus, S. T. Espenschied, H. H. Willcockson, C. P. Mack, K. M. Caron, Fetal-derived adrenomedullin mediates the innate immune milieu of the placenta. J Clin Invest 123, 2408–2420 (2013).

54. K. Marzusch, P. Ruck, A. Geiselhart, R. Handgretinger, J. A. Dietl, E. Kaiserling, H. P. Horny, G. Vince, C. W. Redman, Distribution of cell adhesion molecules on CD56++, CD3-, CD16-large granular lymphocytes and endothelial cells in first-trimester human decidua. Hum Reprod 8, 1203–1208 (1993).

55. P. Ruck, K. Marzusch, E. Kaiserling, H. P. Horny, J. Dietl, A. Geiselhart, R. Handgretinger, C. W. Redman, Distribution of cell adhesion molecules in decidua of early human pregnancy. An immunohistochemical study. Lab Invest 71, 94–101 (1994).

56. C. Carlino, H. Stabile, S. Morrone, R. Bulla, A. Soriani, C. Agostinis, F. Bossi, C. Mocci, F. Sarazani, F. Tedesco, A. Santoni, A. Gismondi, Recruitment of circulating NK cells through decidual tissues: a possible mechanism controlling NK cell accumulation in the uterus during early pregnancy. Blood 111, 3108–3115 (2008).

57. R. Wieser, J. L. Wrana, J. Massagué, GS domain mutations that constitutively activate T beta R-I, the downstream signaling component in the TGF-beta receptor complex. EMBO J 14, 2199–2208 (1995).

58. M. Zhou, J. Wang, J. Pan, H. Wang, L. Huang, B. Hou, Y. Lai, F. Wang, Q. Guan, F. Wang, Z. Xu, H. Yu, Nanovesicles loaded with a TGF-β receptor 1 inhibitor overcome immune resistance to potentiate cancer immunotherapy. Nat Commun 14, 3593 (2023).

59. M. Saraiva, P. Vieira, A. O’Garra, Biology and therapeutic potential of interleukin-10. J Exp Med 217, e20190418 (2020).

60. J. A. Aas, B. J. Paster, L. N. Stokes, I. Olsen, F. E. Dewhirst, Defining the normal bacterial flora of the oral cavity. J Clin Microbiol 43, 5721–5732 (2005).

61. D. B. Keskin, D. S. J. Allan, B. Rybalov, M. M. Andzelm, J. N. H. Stern, H. D. Kopcow, L.

62. A. Koopman, J. L. Strominger, TGFbeta promotes conversion of CD16+ peripheral blood NK cells into CD16-NK cells with similarities to decidual NK cells. Proc Natl Acad Sci U S A 104, 3378–3383 (2007).

62. D. K. Sojka, L. Yang, W. M. Yokoyama, Uterine Natural Killer Cells. Front Immunol 10, 960 (2019).

63. D. K. Sojka, L. Yang, B. Plougastel-Douglas, D. A. Higuchi, B. A. Croy, W. M. Yokoyama, Cutting Edge: Local Proliferation of Uterine Tissue-Resident NK Cells during Decidualization in Mice. J Immunol 201, 2551–2556 (2018).

64. X. Fang, N. Ni, Y. Gao, J. P. Lydon, I. Ivanov, M. Rijnkels, K. J. Bayless, Q. Li, Transforming growth factor beta signaling and decidual integrity in mice†. Biol Reprod 103, 1186–1198 (2020).

65. S. Viel, A. Marçais, F. S.-F. Guimaraes, R. Loftus, J. Rabilloud, M. Grau, S. Degouve, S. Djebali, A. Sanlaville, E. Charrier, J. Bienvenu, J. C. Marie, C. Caux, J. Marvel, L. Town, N. D. Huntington, L. Bartholin, D. Finlay, M. J. Smyth, T. Walzer, TGF-β inhibits the activation and functions of NK cells by repressing the mTOR pathway. Sci Signal 9, ra19 (2016).

66. J. D. Barahona, L. Yang, D. M. Nelson, W. M. Yokoyama, TGF-β drives the conversion of conventional NK cells into uterine tissue-resident NK cells to support murine pregnancy. eLife 15, RP109878 (2026).

67. S. M. Fahim, S. Das, M. A. Gazi, M. A. Alam, M. Mahfuz, T. Ahmed, Evidence of gut enteropathy and factors associated with undernutrition among slum-dwelling adults in Bangladesh. Am J Clin Nutr 111, 657–666 (2020).

68. P. Bankhead, M. B. Loughrey, J. A. Fernández, Y. Dombrowski, D. G. McArt, P. D. Dunne, S. McQuaid, R. T. Gray, L. J. Murray, H. G. Coleman, J. A. James, M. Salto-Tellez, P. W. Hamilton, QuPath: Open source software for digital pathology image analysis. Sci Rep 7, 16878 (2017).

69. J. Kieckbusch, L. M. Gaynor, A. Moffett, F. Colucci, MHC-dependent inhibition of uterine NK cells impedes fetal growth and decidual vascular remodelling. Nat Commun 5, 3359 (2014).

70. M. Arenas-Hernandez, E. N. Sanchez-Rodriguez, T. N. Mial, S. A. Robertson, N. Gomez-Lopez, Isolation of Leukocytes from the Murine Tissues at the Maternal-Fetal Interface. J Vis Exp, e52866 (2015).

71. M. I. Love, W. Huber, S. Anders, Moderated estimation of fold change and dispersion for RNA-seq data with DESeq2. Genome Biol 15, 550 (2014).

72. G. Korotkevich, V. Sukhov, N. Budin, B. Shpak, M. N. Artyomov, A. Sergushichev, Fast gene set enrichment analysis. bioRxiv [Preprint] (2016). 10.1101/060012.

73. A. S. Castanza, J. M. Recla, D. Eby, H. Thorvaldsdóttir, C. J. Bult, J. P. Mesirov, Extending support for mouse data in the Molecular Signatures Database (MSigDB). Nat Methods 20, 1619–1620 (2023).

74. H.-W. Chang, E. M. Lee, Y. Wang, C. Zhou, K. M. Pruss, S. Henrissat, R. Y. Chen, C. Kao, M. C. Hibberd, H. M. Lynn, D. M. Webber, M. Crane, J. Cheng, D. A. Rodionov, A. A. Arzamasov, J. J. Castillo, G. Couture, Y. Chen, N. P. Balcazo, C. B. Lebrilla, N. Terrapon, B. Henrissat, O. Ilkayeva, M. J. Muehlbauer, C. B. Newgard, I. Mostafa, S. Das, M. Mahfuz, A. L. Osterman, M. J. Barratt, T. Ahmed, J. I. Gordon, Prevotella copri and microbiota members mediate the beneficial effects of a therapeutic food for malnutrition. Nat Microbiol 9, 922–937 (2024).

75. S. J. Fleming, M. D. Chaffin, A. Arduini, A.-D. Akkad, E. Banks, J. C. Marioni, A. A. Philippakis, P. T. Ellinor, M. Babadi, Unsupervised removal of systematic background noise from droplet-based single-cell experiments using CellBender. Nat Methods 20, 1323– 1335 (2023).

76. Y. Hao, T. Stuart, M. H. Kowalski, S. Choudhary, P. Hoffman, A. Hartman, A. Srivastava, G. Molla, S. Madad, C. Fernandez-Granda, R. Satija, Dictionary learning for integrative, multimodal and scalable single-cell analysis. Nat Biotechnol 42, 293–304 (2024).

77. C. S. McGinnis, L. M. Murrow, Z. J. Gartner, DoubletFinder: Doublet Detection in Single-Cell RNA Sequencing Data Using Artificial Nearest Neighbors. Cell Syst 8, 329–337.e4 (2019).

78. C. Hafemeister, R. Satija, Normalization and variance stabilization of single-cell RNA-seq data using regularized negative binomial regression. Genome Biol 20, 296 (2019).

79. S. Choudhary, R. Satija, Comparison and evaluation of statistical error models for scRNA-seq. Genome Biol 23, 27 (2022).

80. M. Büttner, J. Ostner, C. L. Müller, F. J. Theis, B. Schubert, scCODA is a Bayesian model for compositional single-cell data analysis. Nat Commun 12, 6876 (2021).

81. I. Korsunsky, N. Millard, J. Fan, K. Slowikowski, F. Zhang, K. Wei, Y. Baglaenko, M. Brenner, P.-R. Loh, S. Raychaudhuri, Fast, sensitive and accurate integration of single-cell data with Harmony. Nat Methods 16, 1289–1296 (2019).

82. A. Janesick, R. Shelansky, A. D. Gottscho, F. Wagner, S. R. Williams, M. Rouault, G. Beliakoff, C. A. Morrison, M. F. Oliveira, J. T. Sicherman, A. Kohlway, J. Abousoud, T. Y. Drennon, S. H. Mohabbat, 10x Development Teams, S. E. B. Taylor, High resolution mapping of the tumor microenvironment using integrated single-cell, spatial and in situ analysis. Nat Commun 14, 8353 (2023).

83. L. Fei, H. Chen, L. Ma, W. E R. Wang, X. Fang, Z. Zhou, H. Sun, J. Wang, M. Jiang, X. Wang, C. Yu, Y. Mei, D. Jia, T. Zhang, X. Han, G. Guo, Systematic identification of cell-fate regulatory programs using a single-cell atlas of mouse development. Nat Genet 54, 1051–1061 (2022).

84. R. Wang, P. Zhang, J. Wang, L. Ma, W. E S. Suo, M. Jiang, J. Li, H. Chen, H. Sun, L. Fei, Z. Zhou, Y. Zhou, Y. Chen, W. Zhang, X. Wang, Y. Mei, Z. Sun, C. Yu, J. Shao, Y. Fu, Y. Xiao, F. Ye, X. Fang, H. Wu, Q. Guo, X. Fang, X. Li, X. Gao, D. Wang, P.-F. Xu, R. Zeng, G. Xu, L. Zhu, L. Wang, J. Qu, D. Zhang, H. Ouyang, H. Huang, M. Chen, S.-C. Ng, G.-H. Liu, G.-C. Yuan, G. Guo, X. Han, Construction of a cross-species cell landscape at single-cell level. Nucleic Acids Res 51, 501–516 (2023).

85. R. A. Amezquita, A. T. L. Lun, E. Becht, V. J. Carey, L. N. Carpp, L. Geistlinger, F. Marini, K. Rue-Albrecht, D. Risso, C. Soneson, L. Waldron, H. Pagès, M. L. Smith, W. Huber, M. Morgan, R. Gottardo, S. C. Hicks, Orchestrating single-cell analysis with Bioconductor. Nat Methods 17, 137–145 (2020).

86. T. Moore, J. M. Williams, M. A. Becerra-Rodriguez, M. Dunne, R. Kammerer, G. Dveksler, Pregnancy-specific glycoproteins: evolution, expression, functions and disease associations. Reproduction 163, R11–R23 (2022).

87. G. N. Sulkowski, J. Warren, C. T. Ha, G. S. Dveksler, Characterization of receptors for murine pregnancy specific glycoproteins 17 and 23. Placenta 32, 603–610 (2011).

88. S. M. Blois, I. Tirado-González, J. Wu, G. Barrientos, B. Johnson, J. Warren, N. Freitag, B. F. Klapp, S. Irmak, S. Ergun, G. S. Dveskler, Early expression of pregnancy-specific glycoprotein 22 (PSG22) by trophoblast cells modulates angiogenesis in mice. Biol Reprod 86, 191 (2012).

89. J. A. Wu, B. L. Johnson, Y. Chen, C. T. Ha, G. S. Dveksler, Murine pregnancy-specific glycoprotein 23 induces the proangiogenic factors transforming-growth factor beta 1 and vascular endothelial growth factor a in cell types involved in vascular remodeling in pregnancy. Biol Reprod 79, 1054–1061 (2008).

90. E. Wennerberg, V. Kremer, R. Childs, A. Lundqvist, CXCL10-induced migration of adoptively transferred human natural killer cells toward solid tumors causes regression of tumor growth in vivo. Cancer Immunol Immunother 64, 225–235 (2015).

91. A. L. Gregory, G. Xu, V. Sotov, M. Letarte, Review: the enigmatic role of endoglin in the placenta. Placenta 35 Suppl, S93-99 (2014).

92. Q.-X. Xu, W.-Q. Zhang, L. Lu, K.-Z. Wang, R.-W. Su, Distinguish Characters of Luminal and Glandular Epithelium from Mouse Uterus Using a Novel Enzyme-Based Separation Method. Reprod Sci 30, 1867–1877 (2023).

93. S. H. Robbins, K. B. Nguyen, N. Takahashi, T. Mikayama, C. A. Biron, L. Brossay, Cutting edge: inhibitory functions of the killer cell lectin-like receptor G1 molecule during the activation of mouse NK cells. J Immunol 168, 2585–2589 (2002).

94. N. D. Huntington, H. Tabarias, K. Fairfax, J. Brady, Y. Hayakawa, M. A. Degli-Esposti, M. J. Smyth, D. M. Tarlinton, S. L. Nutt, NK cell maturation and peripheral homeostasis is associated with KLRG1 up-regulation. J Immunol 178, 4764–4770 (2007).

95. Š. Magister, H.-C. Tseng, V. T. Bui, J. Kos, A. Jewett, Regulation of split anergy in natural killer cells by inhibition of cathepsins C and H and cystatin F. Oncotarget 6, 22310–22327 (2015).

96. Z. Chen, J. Zhang, K. Hatta, P. D. A. Lima, H. Yadi, F. Colucci, A. T. Yamada, B. A. Croy, DBA-lectin reactivity defines mouse uterine natural killer cell subsets with biased gene expression. Biol Reprod 87, 81 (2012).

97. S. Sivori, M. Vitale, L. Morelli, L. Sanseverino, R. Augugliaro, C. Bottino, L. Moretta, A. Moretta, p46, a novel natural killer cell-specific surface molecule that mediates cell activation. J Exp Med 186, 1129–1136 (1997).

98. A. Pessino, S. Sivori, C. Bottino, A. Malaspina, L. Morelli, L. Moretta, R. Biassoni, A. Moretta, Molecular cloning of NKp46: a novel member of the immunoglobulin superfamily involved in triggering of natural cytotoxicity. J Exp Med 188, 953–960 (1998).

99. T. Walzer, M. Bléry, J. Chaix, N. Fuseri, L. Chasson, S. H. Robbins, S. Jaeger, P. André, L. Gauthier, L. Daniel, K. Chemin, Y. Morel, M. Dalod, J. Imbert, M. Pierres, A. Moretta, F. Romagné, E. Vivier, Identification, activation, and selective in vivo ablation of mouse NK cells via NKp46. Proc Natl Acad Sci U S A 104, 3384–3389 (2007).

100. E. Giuliani, K. L. Parkin, B. A. Lessey, S. L. Young, A. T. Fazleabas, Characterization of uterine NK cells in women with infertility or recurrent pregnancy loss and associated endometriosis. Am J Reprod Immunol 72, 262–269 (2014).

101. Z. Jia, Y. Wei, Y. Zhang, K. Song, J. Yuan, Metabolic reprogramming and heterogeneity during the decidualization process of endometrial stromal cells. Cell Commun Signal 22, 385 (2024).

102. M. Dalamaga, G. S. Christodoulatos, C. S. Mantzoros, The role of extracellular and intracellular Nicotinamide phosphoribosyl-transferase in cancer: Diagnostic and therapeutic perspectives and challenges. Metabolism 82, 72–87 (2018).

103. M. Yang, J. Ong, F. Meng, F. Zhang, H. Shen, K. Kitt, T. Liu, W. Tao, P. Du, Spatiotemporal insight into early pregnancy governed by immune-featured stromal cells. Cell 186, 4271–4288.e24 (2023).

104. D. J. Stadtmauer, S. Basanta, J. D. Maziarz, A. G. Cole, G. Dagdas, G. R. Smith, F. van Breukelen, M. Pavličev, G. P. Wagner, Cell type and cell signalling innovations underlying mammalian pregnancy. Nat Ecol Evol 9, 1469–1486 (2025).

105. N. Salleh, N. Giribabu, Leukemia inhibitory factor: roles in embryo implantation and in nonhormonal contraception. ScientificWorldJournal 2014, 201514 (2014).

106. L. L. Shuya, E. M. Menkhorst, J. Yap, P. Li, N. Lane, E. Dimitriadis, Leukemia inhibitory factor enhances endometrial stromal cell decidualization in humans and mice. PLoS One 6, e25288 (2011).

107. H. Lv, J. Tong, J. Yang, S. Lv, W.-P. Li, C. Zhang, Z.-J. Chen, Dysregulated Pseudogene HK2P1 May Contribute to Preeclampsia as a Competing Endogenous RNA for Hexokinase 2 by Impairing Decidualization. Hypertension 71, 648–658 (2018).

108. I. Tamura, H. Takagi, Y. Doi-Tanaka, Y. Shirafuta, Y. Mihara, M. Shinagawa, R. Maekawa, T. Taketani, S. Sato, H. Tamura, N. Sugino, Wilms tumor 1 regulates lipid accumulation in human endometrial stromal cells during decidualization. J Biol Chem 295, 4673–4683 (2020).

