## Supplementary Materials for "Small intestinal microbiota of undernourished women perturbs placental development in mice"

### MATERIALS AND METHODS

#### Diets

The Adult Mirpur diet was formulated based on results of a 24-hour dietary recall survey and food frequency questionnaire taken from adults enrolled in the BEED study (*67*). A pelleted diet was produced by Dyets, Inc. (Bethlehem, PA) and sterilized by irradiation. The ingredients and formulation of the Adult Mirpur diet are described in a previous publication (*33*).

#### Mouse experiments

All mouse experiments were conducted using protocols approved by the Washington University animal Studies Committee (assurance # D16-00245 (A3381-01) and protocol # 23-0271). All animals were housed in plastic flexible film gnotobiotic isolators (Class Biologically Clean Ltd., Madison, WI) and maintained at 23°C under a strict 12-hour light cycle (lights on at 0700h). Autoclaved paper 'shepherd shacks' were kept in each cage to facilitate the natural nesting behaviors and for environmental enrichment.

#### Husbandry

The method used for generating the aSI-L and aSI-H culture collections, and the preparation of strains for gavage into germ-free mice are previously described (*18*). Before oral gavage, GF female C57Bl/6J mice were given autoclaved standard chow (Lab Diet 5021) and sterile water *ad libitum*. Three days before gavage, animals were switched to the pelleted Adult Mirpur diet, which was provided *ad libitum*. 200 µL of the aSI-L or aSI-H consortium was administered via an oral gavage needle (Cadence Science; catalog no. 7901). Gavaged mice were colonized for 17 days before starting timed matings.

In preparation for co-housing experiments, adult female C57Bl/6J mice were switched from an autoclaved standard chow to the Mirpur Adult diet three days before having one colonized aSI-L or aSI-H dam (17 days after oral gavage) placed into the same cage. After 28 days of coprophagic colonization, mice were either maintained in their cage and isolator (non-cohoused aSI-L and aSI-H control groups) or were transferred to a new cage in a new isolator to be co-housed with an equal number of mice transferred from an isolator with the other treatment group. Twenty-eight days later, aSI-L and aSI-H controls and co-housed mice (CoH-L and CoH-H) started timed matings.

Primiparous female mice were subjected to timed matings with 1-2 females placed in a cage with 1-2 males. After 12-16 hours (E0.5), females were checked for a vaginal plug, then separated from the males and placed in their original cages for the duration of pregnancy. Pregnancy was determined by a weight gain > 2 g from E1.5-E10.5. All mice were euthanized without prior fasting on E11.5 or E17.5. Cervical dislocation was used instead of $CO_2$ for euthanasia to prevent perturbations to maternal physiology. Fetal sex was determined using real-time PCR for the Y chromosome and DNA isolated from either fetal heads (E11.5) or tails (E17.5) (Transnetyx; Cordova, TN). All other biospecimens for transcriptomic or proteomic analysis were flash-frozen in liquid nitrogen and stored at −80 °C before use.

#### MFI and intestinal histology and histomorphometric measurements

Intestinal sections of aSI-L and aSI-H dams at E17.5 were fixed for 24 h at 4°C in 10% neutral buffered formalin, rehydrated in 70% ethanol at 4°C, then positioned in 2% agar followed by paraffin embedding and cutting 5 µm sections onto slides. The MFIs of aSI-L, aSI-H, CoH-L, and CoH-H dams were collected at E11.5 and E17.5, rinsed in PBS and fixed for 48 h at 4°C in 10% neutral buffered formalin. Samples were rehydrated in 70% ethanol and refrigerated until processing for paraffin embedding. Serial 10-µm sections were generated. Sections were stained with hematoxylin (Vector, #H-3401) and eosin (Sigma-Aldrich, #HT110132) following standard procedures, then scanned with a Hamamatsu NanoZoomer at 20X resolution. Intestinal villus height and crypt depth measurements and placental LZ/JZ areas and vascular measurements were performed using QuPath v0.5.1 (*68*). Placental zones were manually drawn and annotated. The method used for vascular measurements was adapted from Kieckbusch *et al* (*69*). Vasculature in both the LZ and JZ were assumed to be oval-like in shape and the major/minor axes of the lumen were measured to calculate luminal area. Histomorphometric data were evaluated in a blinded fashion.

**Movat-Russell Pentachrome staining**

E11.5 aSI-L and aSI-H MFI sections were stained with the Movat-Russell Modified Pentachrome Stain Kit (Newcomer Supply, #9150A) following the manufacturer's protocol. Briefly, 10 µm-thick formalin-fixed paraffin-embedded (FFPE) sections were deparaffinized (100% xylene, 100% ethanol, 95% ethanol, $dH_2O$), stained with a 1% Alcian blue stain for 20 min at room temperature (RT), followed by immersion for 30 min in an alkaline alcohol solution (combine 5 mL of 28-30% ammonium hydroxide and 45 mL 95% ethanol). The slides were then stained with a hematoxylin working stain solution, made by combining 10 mL each of 10% hematoxylin, 100% ethanol, 10% ferric chloride, and Verhoeff's iodine, for 15 min at room temperature (RT). Slides were then stained with 2% ferric chloride (5-10 dips) followed by 5% sodium thiosulfate for 1 min. Sections were subsequently stained for 1 min at RT with a Crocein scarlet-acid fuchsin solution, followed by 0.5% acetic acid for 30 sec and 5% phosphotungstic acid for 10 min (both at RT), before being rinsed in 0.5% acetic acid and placed in 1% Orange G stain for 15 min. Slides are mounted using an organic-solvent-based mounting media (Epredia, #22-050-102), and full-tissue sections were scanned with a Hamamatsu NanoZoomer at 20X resolution and analyzed with QuPath v0.5.1 (*68*). Visual assessments were performed blinded, and 8-13 MFI sections from 4-5 dams for each treatment group were analyzed.

**Immunohistochemistry (IHC)**

Both 5 µm-thick FFPE intestinal and 10 µm-thick FFPE MFI sections were deparaffined with xylene, ethanol, and aqueous solutions (100% xylene, 1:1 xylene: ethanol, 100% ethanol, 95% ethanol, 70% ethanol, 50% ethanol, and $dH_20$). Antigen retrieval was performed by boiling in Tris-EDTA Buffer pH 9.0 buffer (Abcam, ab93684) for 20 min. Slides were then washed in Tris-buffered saline (TBS) with 0.025% Triton X-100, and blocked for 2 h at RT with 10% normal goat serum (Abcam, ab7481) and 1% FBS in TBS. Primary antibodies were diluted in antibody diluent (Abcam, ab64211) according to **table S12** and incubated overnight at 4 °C. After washing with TBS-Triton-X 100, sections were incubated with secondary antibodies at a concentration of 2 mg/mL in TBS supplemented with 1% FBS. Sections were mounted with Fluoroshield Mounting Media containing DAPI (Abcam, #ab104139). Full-tissue IHC sections were scanned with a 3DHistech P250. Histological analysis was performed in a blinded fashion, using QuPath for fluorescence quantification (*68*).

**Fluorescent *In situ* hybridization (FISH) of the MFI**

Detection of *Pecam1* mRNA transcript in FFPE E11.5 MFI sections was performed using RNAscope Multiplex Fluorescent Reagent Kit v2 (Advanced Cell Diagnostics, # 323100) following the manufacturer's protocol. Briefly, the 10 μm-thick FFPE sections were incubated for 60 min at 60°C and deparaffinized with xylene followed by 100% ethanol. Endogenous peroxidases were quenched with $H_2O_2$ for 10 min at RT. Slides were then boiled for 15 min in RNAscope Target Retrieval Reagents and incubated for 30 min in RNAscope Protease Plus reagent prior to *Pecam1* RNA probe hybridization, signal amplification and detection with the fluorophore Opal 690 (**table S12**). Slides were subsequently counterstained with 4′,6-diamidino-2-phenylindole (DAPI), and mounted using ProLong Gold Antifade Mountant (Thermo Fisher Scientific, P36930). Full-tissue FISH sections were scanned with a Hamamatsu NanoZoomer at 20X resolution. Histological analysis was performed in a blinded fashion, using QuPath for fluorescence quantification (*68*).

**Flow cytometry of decidual immune populations**

Whole deciduas were dissected apart from fetal placentas and pooled from a dam's entire litter as a single sample. Using procedures adapted from Arenas-Hernandez *et al* (*70*), decidual tissue was minced for 3 min in Accumax solution (Innovative Cell Technologies, #AM105) on ice, and incubated for 30 min at 37 °C with shaking at 80 rpm. The sample was centrifuged at 1,500 x g for 5 min at 4 °C. Decidual cell pellets were then passed through sterile filters with 70 μm and 40 μm pore diameters (Fisher Scientific, #08-771-2 and #08-771-1, respectively). Between each filtering step, cells were washed with sterile Dulbecco's PBS (DPBS) and recovered by centrifugation.

Decidual cells were stained with anti-CD16/32 Fcɣ block (Biolegend, #101320). Dead cells were excluded using the Zombie Red fixable viability kit (Biolegend, #423109). Super Bright Complete Staining Buffer (eBiosciences, #SB-4401-75) was used during antibody-fluorochrome conjugate staining to reduce nonspecific polymer dye-dye interactions. Samples were analyzed using the Cytek Northern Lights spectral cytometer. The fluorophore-labeled monoclonal antibodies used for flow cytometry are listed in **tables S13** and **S14**. Representative gating strategies are shown in **fig. S9A**. Data were analyzed using FlowJo v11.0.2.

**Protein assays of intestinal, fetal placental and decidual/MLAp homogenates**

Fetal placental and decidual/MLAp homogenates were prepared by first taking two-thirds by weight of either an individual E11.5 or E17.5 fetal placenta or an individual E11.5 decidua/MLAp and adding 1 mm diameter Zirconia/Silica beads (BioSpec, #11079110z). Intestinal homogenates were prepared by first adding a 20-30 mg fragment from the first (duodenal) or final (ileal) third of the small intestine to a vial with 1.6 mm aluminum oxide and silicon carbide particles (MP Biomedicals, #1169150). Cold T-PER buffer (Thermo Fisher Scientific, #78510) with Complete Ultra protease inhibitor (Roche) was added to each sample (600 μL for fetal placentas and intestines, 300 μL for deciduas/MLAp) followed by bead beating at 21 ºC (1 min for fetal placenta and intestines; 2 min for deciduas/MLAp). All samples were then centrifuged at 13,000 x g for 5 min at 4 °C. Protein concentrations in the resulting supernatants were measured (Pierce BCA Protein Assay Kit; Thermo Fisher Scientific, #23225) and adjusted to a concentration of 1 mg protein/mL using sterile PBS with protease inhibitor cocktail (Roche, #11697498001). Proteins were analyzed by Eve Technologies Corporation

(Calgary, Canada). For intestinal tissue, the Mouse Cytokine Th17 12-Plex Discovery Assay® Array (MDTH17-12) or a LCN2 ELISA kit (R&D Systems, #DY1857) was applied. For E11.5 and E17.5 fetal placentas, the Mouse Angiogenesis and Growth Factor 16-Plex Discovery Assay® Array (MDAG16) and the Mouse Cytokine/Chemokine 32-Plex Discovery Assay® Array (MD32) were used. For E11.5 deciduas/MLAp, the MDAG16, the Steroid/Thyroid 5-Plex Discovery Assay® Multi Species Array for non-blood-based samples (STTHD-Cell/Tissue), and the TGFβ 3-Plex Discovery Assay® Multi Species Array (TGFβ1-3) were employed.

#### Transcriptional analyses

##### *Bulk RNA-Seq of placenta*

RNA was extracted from one-third (by weight) of flash-frozen fetal placenta samples (RNeasy 96 Kit; Qiagen) (the remaining two-thirds of the fetal placenta was used for protein quantification). Total RNA quantification and quality assessment was performed using a TapeStation (Agilent). cDNA libraries were generated with the Illumina TruSeq Stranded Total RNA Prep with Ribo-Zero kit (Illumina). Barcoded libraries were sequenced on an Illumina NovaSeq 6000 instrument [150 nt pair-end reads to a depth $2.68 \times 10^7 \pm 1.86 \times 10^6$ reads/sample (mean ± SD) for the E11.5 and E17.5 aSI-L/H fetal placenta tables, $6.08 \times 10^7 \pm 5.33 \times 10^6$ reads/sample for the E11.5 co-housed fetal placental table, and $8.02 \times 10^7 \pm 1.25 \times 10^7$ reads/sample for the E17.5 co-housed fetal placental table]. DESeq2-based differential gene expression analysis was performed with statistical significance defined as an adjusted $p < 0.05$ (Wald's test with BH correction) (*71*) (**table S2C**,**D**). Genes were ranked by -$\log_{10}$(adjusted p-value)*$\log_2$(fold-difference) and then GSEA was performed using the fgsea R package (*72*) (**table S2C**,**D**). Mouse gene sets were selected from the orthology-mapped Ontology gene set from MSigDB (*73*) (**table S2C**,**D**). Pathways were selected and ranked based on the normalized enrichment score (NES), adjusted p-value, and the number of significant leading edges (**table S2C**,**D**).

##### *snRNA-seq of placental and intestinal tissue*

*Isolation of nuclei* - At both E11.5 and E17.5, fetal-derived (LZ and JZ) placental tissue was dissected from the underlying maternal-derived tissue (decidua/MLAp combined). Two-thirds of the E11.5 and E17.5 fetal-derived placenta by weight was used for nuclear extraction (n = 1 fetal-placental tissue from a given litter; n = 4 litters/treatment group). For the intestines, a 1 cm fragment positioned 3 cm from the pylorus (duodenum), and a 1.5 cm fragment located 1.5 cm from the ileocecal valve (ileum) were cleared of luminal contents and flash frozen for nuclear extraction. Using methods adapted from a protocol for jejunum (*74*), samples were thawed and minced in lysis buffer [25 mM citric acid, 250 mM sucrose, 0.1% NP-40, and 1X protease inhibitor (Roche)]. Nuclei were extracted via a Dounce homogenizer (Wheaton). E11.5 fetal placentas were dounced 5 times with a loose pestle and 10 times with the tight pestle. E17.5 fetal placentas were dounced 10 times with the loose pestle and 5 times with the tight pestle. Combined E11.5 decidual/MLAp tissue samples were divided in half (by weight), and each half portion was pooled across an entire litter for nuclear extraction (n = 4 litters/treatment group for each assay). E11.5 pooled decidual/MLAp samples were dounced 10 times with the loose pestle and 10 times with the tight pestle in the lysis buffer described for the E11.5 and E17.5 fetal placental tissue. Intestinal tissue was dounced 10 times with the loose pestle and 6 times with the tight pestle.

*Processing of nuclei* **-** All samples were washed 3 times with buffer [25 mM citric acid, 0.25 M sucrose, 1X protease inhibitor (Roche)] and filtered successively through 100 µm, 70 µm, and 40 µm pore diameter strainers (pluriSelect) to obtain nuclei in resuspension buffer [25 mM KCl, 3 mM MgCl2, 50 mM Tris, 1 mM DTT, 0.4 U/µL RNase inhibitor (Sigma), and 0.4 U/µL Superase inhibitor (ThermoFisher)]. Approximately 10,000 nuclei per sample were subjected to gel bead-in-emulsion (GEM) generation, reverse transcription, and construction of libraries for sequencing according to the manufacturer's instructions in the 3' gene expression v3.1 kit (10X Genomics). The concentration of the final cDNA libraries was determined by qPCR using the KAPA library Quantification Kit (KAPA Biosystems/Roche) to produce cluster counts appropriate for the Illumina NovaSeq6000 instrument. Normalized libraries were sequenced (NovaSeq6000 S4 Flow Cell using the XP workflow and a 50x10x16x150 sequencing recipe according to the manufacturer's protocol). A sequencing depth of 600 million read pairs was targeted for both the E11.5 and E17.5 fetal-placental samples, 400 million read pairs for the E11.5 decidual/MLAp samples, and 300 million read pairs for each intestinal sample.

*Pre-processing and quality control* -The 10x CellRanger 8.0 pipeline was used to perform read alignment, generate feature-barcode matrices, and conduct quality controls, including introns (GRCm38/mm10). The *remove-backgroup function* in CellBender (v0.2.2) rescued RNA that was identified as empty droplets, and removed empty droplets identified as RNA (ambient RNA) (*75*). Using the R package Seurat v4.0 or 5.0, filtered feature-barcode matrices were imported as Seurat objects and sample integration, count normalization, cell clustering, and marker gene identification was performed (*76*). Samples were filtered to remove low-quality nuclei (defined as nuclei with < 200 or > 5000 genes or < 400 UMIs). Nuclei with >5% reads from mitochondrial genes and/or >5% reads from ribosomal protein genes were excluded from further analyses. Each sample had predicted doublets removed via DoubletFinder (*77*) and was normalized via *SCTransform* (*78*, *79*). Samples from the same tissue type (fetal placenta, decidua/MLAp, duodenum or ileum) and same timepoint (E11.5 or E17.5) were integrated using *SelectIntegrationFeatures*, *PrepSCTIntegration*, *FindIntegrationAnchors* and *IntegrateData* from the Seurat package. The integrated tables were subjected to unsupervised clustering using *FindNeighbors* (dimensions = 1:30) and *FindClusters* (resolution = 1.5 for E11.5 decidual/MLAp tissue, 1.0 for sub-clustered immune cells from E11.5 decidual/MLAp tissue, 1.0 for E11.5 fetal placental tissue, 1.5 for E17.5 fetal placental tissue, and 1.4 for E11.5 and E17.5 duodenum and ileum).

*Cell cluster annotation and scCODA* - Annotation was performed using the *FindMarkers* function in Seurat. Manual cell type assignments were conducted based on expression of reported markers (see *Results*, **tables S1A-C**; **S2B**,**D**; **S5B**,**C**,**F**,**G**). scCODA is a Bayesian probabilistic model designed for snRNA-seq tables. It detects 'statistically credible differences' in the proportional representation of clusters between treatment groups. Its Bayesian generalized linear multivariate regression model reduces false positives by accounting for low sample numbers and the compositionality of the table (*80*). Hamiltonian Monte Carlo sampling calculates the posterior inclusion probability of including the effect of treatment in the model. The type I error (false discovery) is derived from the posterior inclusion probability for each effect. The set of 'statistically credible effects' is the largest set of effects that can be chosen without exceeding a user-defined false discovery threshold $\alpha$ ($\alpha = 0.1$).

The scCODA model was applied to the E11.5 and E17.5 fetal placental and E11.5 maternal placental tables using default parameters. Based on scCODA's recommendations, we used cell clusters from each table that had consistent proportional representation across samples and treatment groups. The reference clusters used for each snRNA-seq table's scCODA analysis were as follows: E11.5 fetal placenta – fetal endothelial cells; E17.5 fetal placenta – endothelial cells; and E11.5 deciduas – smooth muscle cells. (For scCODA results, see **tables S3D,H** and **S5D**.)

*Pseudobulk analysis of differential gene expression* - Genes that had low levels of expression (read count < 10) were filtered out before count aggregation across nuclei for a given cell cluster in a given biological sample. Reads from each sample were input into DESeq2 for differential gene expression (DEG) analysis (*71*).

**Spatial Transcriptomics**

All Xenium slides, probes, and other reagents are provided in the Xenium Prime 5K Mouse Pan Tissue and Pathways Assay Kit (10X Genomics, 1000672). The mean DV200 score of samples was 66.07% (**table S6A**), and the DV200 score for each sample is reported in **table S6A**.

*De-paraffinization and de-crosslinking sections* – FFPE E11.5 aSI-L and aSI-H whole conceptus samples (includes uterus, MLAp, decidua, fetal placenta, yolk sac, amnion, and fetus) were cut into 5 µm sections onto Xenium slides and subsequently deparaffinized and de-crosslinked following the manufacturer's protocol (Protocol CG000578 and CG0000580, 10X Genomics). Briefly, the slides were incubated for 3 h at 42°C in an oven and placed in a desiccator overnight at RT. The slides were deparaffinized (100% xylene, 100% ethanol, 96% ethanol, 70% ethanol, and nuclease-free deionized water). A de-crosslinking buffer was added to the slide. The slide was subsequently incubated for 30 min at 80 °C and 10 min at 22 °C in a thermal cycler followed by rinsing in 0.05% PBS with Tween detergent (PBS-T).

*Probe hybridization, ligation, amplification, and autofluorescence quenching* – The de-paraffinized and de-crosslinked samples were hybridized with the pre-designed Xenium Prime 5K Mouse Pan Tissue panel mix following the manufacturer's protocol (Protocol CG000058, 10X Genomics). Ligation of the probe ends generated a circular DNA probe that sealed the junction between the probe region hybridized to the RNA, with subsequent amplification of the ligation products. The final step involved quenching of autofluorescence and staining of nuclei with DAPI. Slides were imaged with the 10X Genomics Xenium analyzer.

*Cell segmentation* – Cell segmentation was performed by using a combination of boundary stains (ATP1A1/CD45/E-Cadherin), interior RNA stain (18S rRNA), and by DAPI-stained nucleus expansion (5.0 µm). For the E11.5 conceptuses, an average of 92.6% of cells were segmented by 18S rRNA staining across all samples.

*Pre-processing and quality control* – Using the R package Seurat v5.0, each sample's cell feature matrix was imported as Seurat objects (*76*). Each sample's Seurat object was merged via the *CreateSeuratObject* function for downstream processing before sample integration. Each sample's cells were filtered based on transcript count; the upper limit cutoff was the 98$^{th}$ percentile and the lower limit cutoff was cells with < 20 transcripts. Library size normalization was performed via *NormalizeData* based on median transcript count. After identifying the top 2,000 variable genes via *FindVariableFeatures*, *SketchData* was applied to a subset of 25,000 cells from each sample via the "*LeverageScore*" method. Samples were integrated with Harmony (*81*) using *IntegrateLayers* for batch correction. The integrated table was then subjected to unsupervised clustering using *FindNeighbors* (dimensions = 1:30), *FindClusters* (resolution = 1.0), and *RunUMAP* (dimensions = 1:30). The batch-corrected clustering and UMAP results from 25,000 cells was projected to all cells in the integrated table via *ProjectData* (dimensions = 1:30).

*Cell cluster identification and annotation* – Integrated clustering results with spatial coordinates were generated for each sample via *FetchData*. The resulting clustering coordinates were applied to each sample through 10X Genomics' Xenium Explorer v4.0 software to visualize the locations of cell clusters (*82*). Annotation was performed using the *FindMarkers* function in Seurat. The marker genes from both the E11.5 fetal-derived placentas and decidua/MLAps snRNA-seq tables were used for cluster identification. For the fetus-exclusive clusters (e.g., cardiac and neuronal cells), we used the Mouse Cell Atlas (MCA) 2.0 (*83*) and 3.0 (*84*). The number of cells detected per cluster is listed in **table S6B**.

*Intercellular signaling via MultiNicheNet* – Signaling between the NK cell clusters (type 1 and 2) and the decidual cell clusters (type 1 and 2) within and between the aSI-L and aSI-H treatment groups was inferred with MultiNicheNet (*38*) using the 'aSI-H versus aSI-L' contrast (**table S6C**,**D**). Briefly, the Seurat object with the transcript count data was converted to a SingleCellExperiment (SCE) object (*85*). This was used as an input for the wrapper function *multinichenet_output*, which yielded pseudobulk expression for ligands or receptors per cell-type-condition-replicate combination via the R package edgeR. To rank ligand-receptor interactions by condition-cell type specificity, prioritization scores were generated by aggregating the average normalized pseudobulk expression across each ligand-receptor interaction, across all sender-receiver cell type combinations. Default values were used for the following parameters of the *multinichenet_output* function: 'default ligands', 'receptors', 'pseudobulk results', and 'prioritization scores'. Interactions with a prioritization score > 0.7 were evaluated.

**Bacterial quantitation**

Cecal contents from mice were collected at the time of euthanasia, flash-frozen in liquid nitrogen, and stored at -80 °C. Before DNA extraction, frozen cecal contents were weighed, and 2.22 x $10^8$ cells/mL genome equivalents of the ZymoBIOMICS High Microbial Load Spike-in Control I (D6320) were added. DNA was extracted using 0.1 mm silica beads and one 3.97 mm steel ball in 500 µL of phenol:chloroform:isoamyl alcohol and 710 µL of a 500:210 (v/v) mixture of 2X buffer A:20%SDS followed by bead-beating for 4 min. Crude DNA was purified (Qiagen, QIAquick PCR purification kit). The resulting purified DNA was quantified with Qubit and normalized to a concentration of 0.4 ng/µL.

Full-length bacterial 16S rRNA amplicon libraries were prepared for sequencing according to protocols described by the manufacturer (PacBio, Kinnex); libraries were sequenced on a Sequel II instrument (3.48 x $10^5$ ± 2.06 x $10^5$ reads/sample). De-concatenation was performed using Read Segmentation, and the resulting demultiplexed reads were trimmed (*TrimGalore* v0.6.6 *–quality 20*). Adaptors were removed from trimmed reads with *Cutadapt* (-g

AGRGTTYGATYMTGGCTCAG... AAGTCGTAACAAGGTARCY *--error-rate 0.1 –revcomp*). Subsequent 16S rRNA read processing was performed with the *dada2* pipeline with minor modifications to the *filterAndTrim* step *(minLen=1000, maxLen=1600, maxN=0, rm.phix=TRUE)*. Error rate (*learnErrors*) learning was performed for each fastq file. Dereplication (*deRepFastq*), and *dada* steps were performed with standard parameters. Taxonomic predictions were generated with *assignTaxonomy* using the GreenGenes (2024) database.

In *phyloseq*, bacterial reads were normalized to spike-in counts to approximate absolute abundance in cecal contents. ASV abundances were collapsed into species-level abundances using *tax_glom*(*taxrank = "Species"*). Bacteria with a prevalence of less than 3 mice were removed from subsequent analyses. Differential abundance analyses were performed using FDR-corrected Wilcoxon rank-sum and Kruskal-Wallis followed by Tukey's post-hoc tests (FDR < 0.05).

16S rRNA sequences of bacteria that colonized mice were aligned to the 16S rRNA sequences of bacteria detectable in human duodenal aspirates using the R package *Biostrings* (*pairwiseAlignment(type = 'global')*) after conversion to a *DNAStringSet* object. Percent identity was calculated (*pid(type = 'PID1')*); a 98.7% threshold was used to consider a species match. Plasma SomaScan data were filtered to remove the 30% of aptamers with the lowest variance and subsequently correlated with $\log_{10}$-transformed bacterial species abundance using *cor.test(method = 'spearman')*. Enrichment analyses were performed for each set of proteins significantly (adjusted p-value<0.05) positively (rho>0) or negatively (rho<0) correlated with the absolute abundance of each bacterium in duodenal aspirates using the R package *gprofiler2(query, 'hsapiens', sources = c("GO:BP", "GO:MF", "KEGG", "REAC"), correction_method = 'fdr', significant = T, evcodes = F, mutli_query = F)*. Mouse homologues of human plasma proteins were found using the mapping tool SynGO (https://www.syngoportal.org/convert, version 25-03-2025).

**Statistical analysis**

Statistical significance was assigned to p-values < 0.05 using either non-parametric Wilcoxon rank-sum test for maternal measurements or generalized linear mixed models for measurements involving multiple litter members or multiple intestinal histomorphometric measurements per dam (*lmerTest* package in R). The Benjamini-Hochberg (BH) False Discovery method was used when accounting for multiple comparisons, unless otherwise indicated. Statistical tests, the number of animals analyzed, and mean values and standard deviations, and groups used for statistical comparisons are described in the Figures and associated legends.

#### SUPPLEMENTARY RESULTS

##### Maternal-derived compartments are more affected by aSI-L colonization compared to fetal-derived compartments within the MFI

***Comparison of E11.5 aSI-L versus aSI-H fetal placentas***

Based on nuclei counts from snRNA-seq, dissected 'fetal' concentrated placentas had 51% and 18% maternal-derived cell types in the E11.5 and E17.5 tables, respectively (**table S3A**,**D**). Proteomic analyses of homogenates generated from these fetal preparations revealed significant increases in levels of pro-inflammatory proteins, such as eotaxin, interleukin (IL)-2, leukemia inhibitory factor (LIF), CXCL9, and monokine induced by gamma interferon (MIG) in aSI-L compared to aSI-H mice (**fig. S5A**). Based on bulk RNA-seq, these E11.5 aSI-L fetal-placental preparations also had decreased expression of genes encoding pregnancy-specific glycoproteins (PSG) and carcinoembryonic antigen-related cell adhesion molecule (CEACAM) family members ($\log_2$(fold-difference) < -1.5, FDR < 0.05, Wald test) (**fig S5B**, **table S2A**).

We previously observed a decrease in PSG and CECAM gene expression in fetal placentas from GF mice compared to placentas harvested from their CONV-R or conventionalized (CONV-D) counterparts (the latter are GF mice that had been colonized with cecal microbial communities harvested from CONV-R animals) (*14*). PSGs are produced by trophoblasts and have roles in autocrine and paracrine immunomodulatory, angiogenic, and cell adhesion functions (*86*). *Psg16*, *Psg21*, and *Psg23*, which have significantly reduced expression in aSI-L compared to aSI-H placentas, are produced by junctional zone (JZ)-specific SpT cells (*86*, *87*). In mice, PSG17 and PSG22 bind heparin and heparan sulfate proteoglycans, and PSG23 induces TGF-$\beta$1 and vascular endothelial growth factor (VEGF) in yolk sac-derived endothelial cells and labyrinth zone (LZ) trophoblasts, indicative of their pro-angiogenic function (*87–89*).

The E11.5 aSI-L fetal placentas also had significant increases in expression of genes that are (i) involved in inhibition of cellular proliferation (*Plac9*, *Mt3*, *Gml*, *Rprm*), (ii) components of the extracellular matrix (ECM) (*Adamtsl3*, *Eln*, *S100a4*), and (iii) participate in NK cell activation (*Klra18*, *Ncr1*, *Gzma*, *Klrb1c, Ptgds*) (**fig. S5C**, **table S2A**). GSEA identified significantly higher normalized enrichment scores among aSI-L compared with aSI-H fetal placental transcripts for GOBP terms related to immune activation, and vasculature and tissue remodeling (**fig. S5D**, **table S2B**). In addition to transcriptomic evidence for effects at the ECM, Movat's pentachrome staining with optical density normalization revealed decreased collagen and reticular fibers in the deciduas and MLAp of aSI-L dams (**fig. S5E**). Overall, both the proteomic and transcriptomic data point towards disrupted ECM and immune response in the E11.5 aSI-L placentas.

To identify the cell types responsible for these transcriptomic perturbations seen in E11.5 aSI-L placentas, we conducted snRNA-seq of the E11.5 aSI-L and aSI-H 'fetal' placental preparations (**fig. S6A**, **table S3A-C**). Nuclei clusters were identified using previously described marker genes (*14*). Fetal placenta-derived cell types included (i) trophoblast-precursors [labyrinth trophoblast progenitors (LaTP), junctional zone precursors (JZP), and spongiotrophoblast (SpT) precursors], (ii) fully differentiated trophoblast subtypes [SpTs, glycogen cells (GCs), S-TGCs, and four transcriptionally-distinct syncytiotrophoblast type I and II populations (SynTI 1, SynTI 2, SynTII 1, SynTII 2)], (iii) yolk sac cells, (iv) fetal mesenchymal cells, and (v) fetal endothelial cells. In addition, maternal clusters were identified that included stromal cells, endothelial cells, immune cells, plus a separate NK cell cluster (**fig. S6A**, **table S3B**). The pericyte cluster represented a mixture of maternal and fetal-derived cells (**fig. S6A**, **table S3B**).

We utilized scCODA (*80*), a Bayesian model for evaluating the compositional differences between different treatment groups, to assess whether there were differences in the proportion of each cluster between the E11.5 aSI-L and aSI-H groups (**fig. S6B**, **table S3D**). The results indicated that there was an increased proportion of maternal endothelial and stromal clusters in aSI-L compared to aSI-H placentas (**fig. S6B**, **table S3D**). Consistent with the scCODA results for maternal endothelial cells, E11.5 fetal placental homogenates revealed significant increases in angiopoietin-2, follistatin, and VEGF-A and a decrease in amphiregulin in aSI-L compared to aSI-H mice (n = 4 placentas/dam, 4-5 litters/group; $p < 0.05$, linear mixed model) (**fig. S5F**). Fluorescent *in situ* hybridization (FISH) for *Pecam1*, which encodes the endothelial cell marker CD31, demonstrated no differences in staining in the LZ ($p = 0.98$) (**fig. S6C**,**D**), but a significant increase in the deciduas of aSI-L dams (**fig. S6E**,**F)**. Histomorphometric analysis of hematoxylin and eosin (H&E) stained E11.5 placental sections showed no significant difference in LZ vessel wall thickness ($p = 0.41$) or luminal area ($p = 0.36$) in E11.5 decidual vessel thickness ($p = 0.55$) or luminal area ($p = 0.99$) between aSI-L and aSI-H placentas (n = 2-3 sections/placenta; 1-2 placentas from 6-8 litters/group; linear mixed model) (**fig. S7B**,**C**,**F**,**G**). Collectively, these results revealed an increase in maternal endothelial cells in aSI-L colonized mice. However, since there were no differences in decidual vessel thickness or luminal area, we surmised that the function of uNK cells in remodeling maternal vessels (*62*) was not different in aSI-L compared to aSI-H colonized dams.

We next employed pseudobulk analysis of the snRNA-seq table to identify genes with statistically significant differences in their expression between all aSI-L and aSI-H placental clusters ($|\log_2(\text{fold-difference})|>0.5$, FDR < 0.05, Wald test) (**fig. S6G**, **table S3C**). The maternal endothelial cell cluster had the most DEGs, with the top 27 indicative of increased inhibition of cellular proliferation (*Cdkn1c*, *Fhit*, *Epb41l3*, *Usp29*, *Rad51b*, *Ctdspl*, *Zbtb7c*, *Glis1, Arhgap10*), increased endothelial cell stress (*Slco2b1*, *H19*, *Rbbp7*, *Colec12*, *Lgals3*, *Morf4l2*, *Zbtb7c*, *Sgpp2*), increased angiogenesis (*Arhgap6*, *Serpine2*, *Gata3*, *Mitf*, *Basp1*, *Csf1r*), and increased markers of cell polarity and orientation (*Pdzrn3*, *Patj*, *Fn1*, *Pard3*) (**fig. S6G**,**H**, **table S3C**). The maternal stromal cell cluster had the second highest number of DEGs, with the top 9 DEGs indicative of increases in cellular proliferation (*Kif1a*, *Nxn*), plus decreases in decidual ECM remodeling (*Col3a*, *1Dusp1, Tfpi, Itga8*), vitamin A metabolism (*Bco1*), carnitine transport (*Slc22a5*), and calcium release (*Ryr2*) (**fig. S6G**,**I**, **table S3C**).

***Comparison of E17.5 aSI-L versus aSI-H fetal placentas***

We also performed bulk RNA-seq, snRNA-seq, and proteomic analysis on E17.5 fetal placental preparations. The bulk RNA-seq results indicated that in aSI-L mice there was increased expression of genes associated with immune responses (*Mmp7*, *Cd5l*, *S100a9*, *Acod1*, *Pfpl*), granzymes (*Gzmb*, *Gzmg*, *Gzmd*, *Gzmf*, *Gzmc*, *Gzme*) and NK cell-directed cytotoxicity (*Fcrl6*, *Sh2d1b2*, *Prf1, Cst7*) ($|\log_2(\text{fold-difference})|>1$, FDR < 0.05, Wald test) (**fig. S8A**, **table S2C**). There was also decreased expression of genes associated with steroid hormone signaling and metabolism (*Gper1*, *Prok1*, *Hsd3b1*), vascular tone (*Nppb*, *Adgra1*, *Tph1*), and the ECM (*Elfn1*) (**fig. S8A**, **table S2C**). GSEA disclosed that E17.5 aSI-L placentas had increases in GOBP terms related to organic acid transport, nucleic acid-related processes, G protein-coupled receptor regulation, energy production, and cellular organization (**fig. S8B**, **table S2D**). There was a corresponding decrease in GOBP terms related to immune responses to pathogens and ECM structure (**fig. S8B**, **table S2D**).

Homogenates prepared from E17.5 aSI-L fetal placental preparations had increased concentrations of eotaxin, but decreased concentrations of CXCL9 and CXCL10 (n = 2-4 placentas/dam, 4-5 litters/group; $p < 0.05$, linear mixed model) (**fig. S8C**). The latter is notable given that $CXCR3^+$ NK cells migrate towards high CXCL10 gradients (*90*). Pro-angiogenesis protein angiopoietin-2 was increased, but endoglin, a co-receptor for TGF-β, was decreased in E17.5 aSI-L fetal placental homogenates (n = 2-4 placentas/dam, 4-5 litters/group; $p < 0.05$, linear mixed model) (**fig. S8D**) (*91*).

Similar to the E11.5 fetal placental preparations, E17.5 'fetal' placental preparations contained both maternal and fetal cell clusters (**fig. S9A**, **table S3E-G**). Unlike the E11.5 pseudobulk results, there were fewer DEGs across all E17.5 fetal placental clusters (54 total DEGs) compared to the number of DEGs across all E11.5 clusters (99 total DEGs) ($|\log_2(\text{fold-difference})|>0.5$, FDR $< 0.05$, Wald test) (**fig. S9B**, **table S3G)**. The two E17.5 clusters with the highest number of DEGs were the yolk sac 1 and endothelial cell precursor clusters (**fig. S9B)**. scCODA did not reveal any difference in the abundances of E17.5 fetal-derived clusters across treatment groups (**fig. S9C**, **table S3H**).

H&E staining of E17.5 placental sections demonstrated no significant difference in the cross-sectional area of the LZ in aSI-L versus aSI-H placentas (n = 2 sections/placenta; 1-2 placentas from each of 6-7 litters/group; p = 0.88; linear mixed model) (**fig. S7A**). In the case of E11.5 placental preparations, histomorphometry of E17.5 LZ vessel thickness (p = 0.64) and luminal area (p = 0.36), as well as E17.5 decidual vessel thickness (p = 0.60) and luminal area showed no significant differences between the aSI-L and aSI-H groups (p = 0.39) (n = 2-3 sections/placenta; 1-2 placentas from 6 litters/group; linear mixed model with BH correction) (**fig. S8D**,**E**,**H**,**I**).

In summary, despite our efforts to remove the majority of maternal tissue from our 'fetal' placental preparations, these results disclosed that compared to aSI-H placentas, aSI-L placentas exhibit the most pronounced perturbations in maternal tissue. The alterations in the aSI-L maternal-derived placental tissue were more evident at the E11.5 rather than the E17.5 timepoint. Both timepoints demonstrated that aSI-L maternal placental cells had pro-inflammatory and disrupted ECM responses. However, the vascular remodeling function of uNK cells appears to be intact based on vascular measurements at both gestational stages. These findings spurred us to focus exclusively on the maternal placental components (*i.e.* the decidua and MLAp) and on the earlier gestational timepoint of E11.5. These analyses are described in the main *Results* section.

**Marker genes for NK and decidual cell clusters from spatial transcriptomics**

For our spatial transcriptomics analysis, we examined 805,053 cells across six samples (n = 3 E11.5 aSI-L and 3 E11.5 aSI-H). This yielded 27 clusters, with maternal-derived cell types consisting of (i) immune cells including two NK cell clusters (which we designated type 1 and 2) and two macrophage clusters (M1 and M2), (ii) three decidual stromal cell clusters (designated types 1, 2 and 3), along with the superficial and deep stromal cells described in our decidual/MLAp snRNA-seq clustering, (iii) maternal endothelial cells, including a lymphatic endothelial cell cluster, (iv) uterine smooth muscle cells, and (v) luminal epithelial cells (*i.e.* those that line the central cavity of the uterus, the blastocyst attachment point) (*92*) (**Fig. 4A,B**, **table S6B**). Fetal-derived placental cells included LZ cells, JZ cells, SynT, and P-TGC clusters (**Fig. 4A,B**, **table S6B**).

***NK cell clusters***

Of the two NK cell clusters, 'type 2' was assigned to the uNK subtype given its higher expression of uNK cell-associated genes, such as *Prf1* (perforin 1), *Klrg1* (KLRG1), *Itga1* (CD49a), *Cst7* (cystatin 7) and *Gzmb* (granzyme B) (**Fig. 4C**). KLRG1 is a receptor that inhibits both cytokine and NK-cell mediated cytotoxicity (*93*, *94*). Cystatin 7 decreases cytotoxicity by inhibiting major granzyme convertases – cathepsin C and H (*95*). Granzyme B levels are elevated in uNK (type 2) cells due to their larger cytotoxic granules relative to cNK (type 1) cells, reflecting granule retention rather than increased cytotoxic function (*96*).

In turn, 'type 1' NK cells were considered cNK cells, with higher expression of NK cell activation receptors and NK cell-directed cytotoxic genes, including *Ncr1* (NKp46), *Ptprc* (CD45), *Il2rb* (IL-2 receptor subunit beta), *Gzma* (granzyme A), as well as *Klrb1c* (NK1.1) (**Fig. 4C**). NKp46 triggers NK cell cytotoxicity against infected, transformed, and/or stressed cells (*97–99*). Expression of the NKp46 surface protein on NK cells has been linked to reproductive outcomes; the endometrial stroma of women with endometriosis and unexplained recurrent pregnancy loss (RPL) or infertility have a higher ratio of $NKp46^{+}$:$CD56^{+}$ NK cells (*100*). Cell segmentation revealed cNK (type 1) cells to be concentrated toward the MLAp of the E11.5 MFI compared to uNK (type 2) cells in both treatment groups (**Fig. 4D**).

***Decidual cell clusters***

Three populations of decidual cells were distinguished based on their patterns of gene expression and spatial location. One population (type 1) was identified as immune-associated based on its high expression of genes involved in uNK cell chemotaxis and activation (*Cxcl14*) (*101*), inflammation and trophoblast invasion (*Nampt*) (*102*) and immunomodulatory pre-decidual cell markers (*Smoc2*, *Wt1*) (*103*, *104*). A second population (type 2) was identified as a subset of postmature decidual stromal cells (*103*) (**fig. S12C**); their transcriptomes are enriched in genes involved in lysosomal activity and tissue remodeling (*Ctsk*) (*104*), cytokine and prostaglandin production (*Lifr*) (*105*, *106*), glycolysis (*Hk2*) (*107*), and lipid homeostasis (*Vldlr*) (*108*) (**fig. S12C**). The third population (type 3) is largely confined to the anti-mesometrial pole of the conceptus, where they exhibit high expression of the decidual marker *Prl8a2* and stress-related marker *Cryab*, marking them as an additional subpopulation of postmature decidual cells (*103*, *104*) (**Figs. 4A**, **5A**, **and fig. S12C**).

Spatially, cells belonging to the type 1 NK cluster overlie type 1 decidual cells, while type 2 NK and type 2 decidual cells are dispersed throughout the decidua (**Figs. 4A**, **5A**). Some type 2 NK cells directly contact type 1 decidual cells (**Figs. 4A**, **5A**). However, the two NK cell types have little contact with type 3 decidual cells, which are positioned on the antimesometrial side of the conceptus (**Figs. 4A**, **5A**).

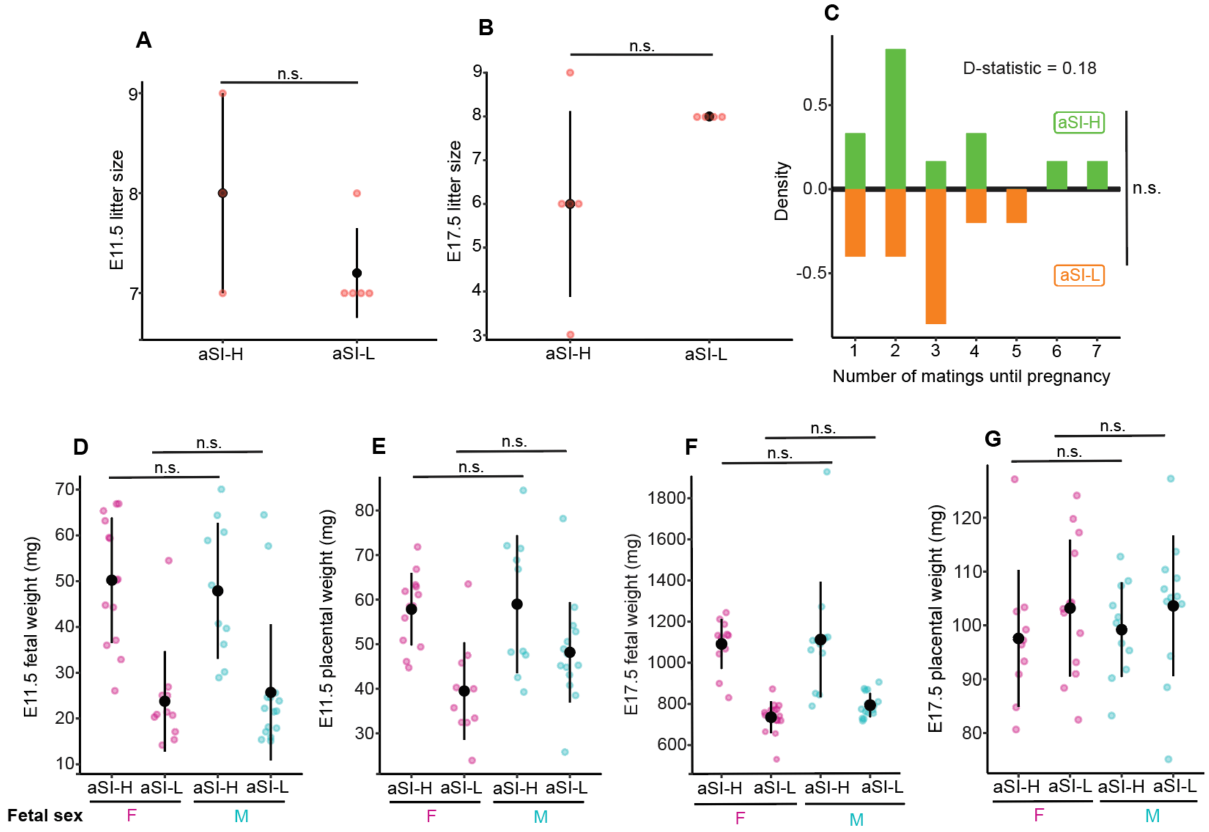


**Figure S1. Litter size, fertility, fetal sex, fetal and placental weights in aSI-L and aSI-H mice.** (**A**,**B**) Litter sizes at E11.5 (aSI-L: 5 litters; aSI-H: 3 litters) and E17.5 (aSI-L: 5 litters; aSI-H: 4 litters; Wilcoxon rank-sum test). **(C)** Density histograms depicting the relative frequency of matings needed for each aSI-L and aSI-H mouse to achieve a successful pregnancy. (**D**,**E**) E11.5 values for fetal weights (aSI-L: 26 fetuses; aSI-H: 24 fetuses) and corresponding placental weights. (**F**,**G**) E17.5 values for fetal weights (aSI-L: 31 fetuses; aSI-H: 27 fetuses) and corresponding placental weights by fetal sex. Linear mixed model used for pairwise comparisons (*Fetal or placental weight ~ Fetal Sex + (1|Litter_ID)*). Mean values ± SD are shown for all but panel **C**.

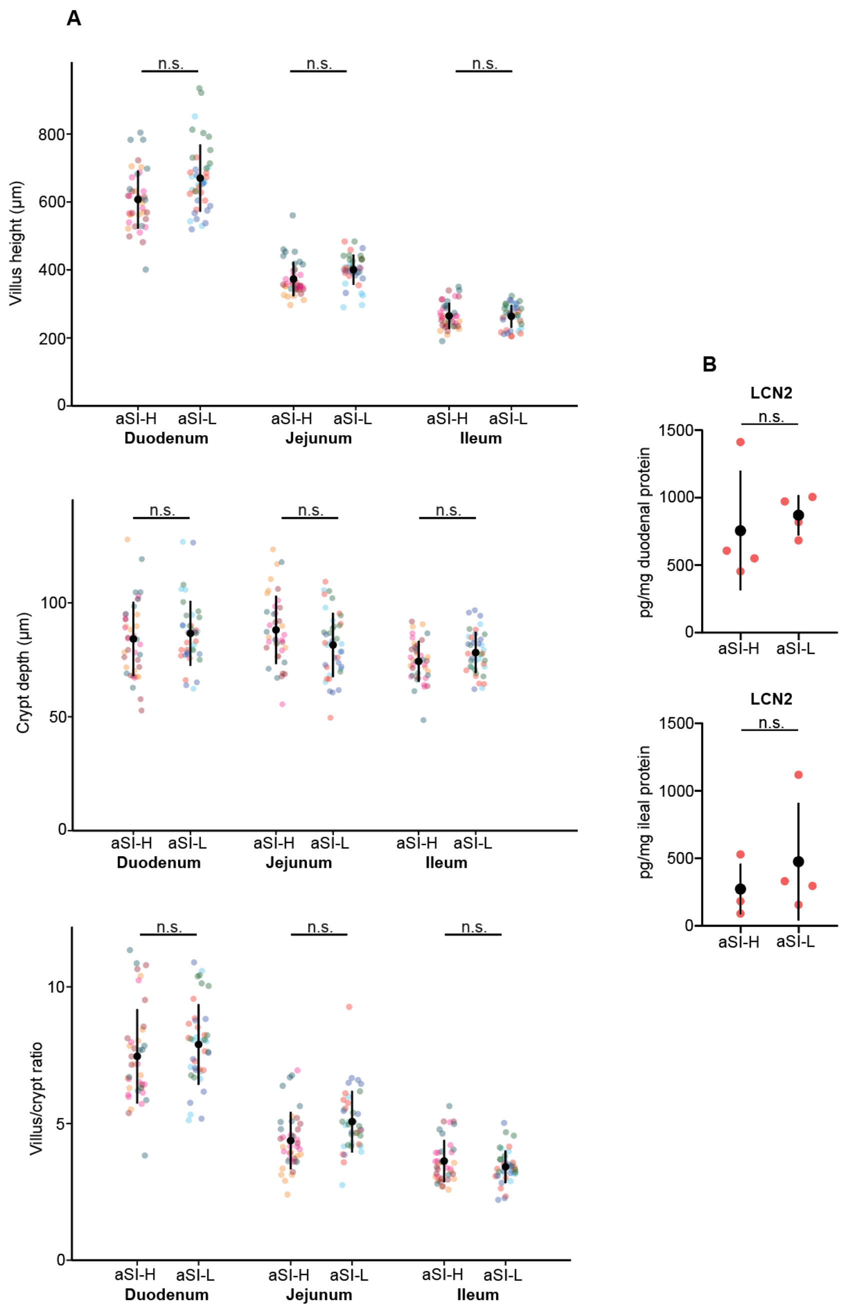


**Figure S2. Intestinal histomorphometrics and LCN2 levels at E17.5. (A)** Villus height, crypt depth, and villus-to-crypt ratios from duodenum, jejunum, and ileum from E17.5 dams from the aSI-L and aSI-H treatment groups (n = 10 measures/intestinal segment/dam for n = 4-6 dams/treatment group). A linear mixed-effects model *(Measure ~ Microbiota + (1 | Mouse ID)*) was used to assess statistical significance. Each color reflects measurements from one mouse. **(B)** LCN2 levels were measured in homogenized duodenum and ileum from E17.5 dams (n = 4/treatment group). The Wilcoxon rank-sum test was used to assess statistical significance.

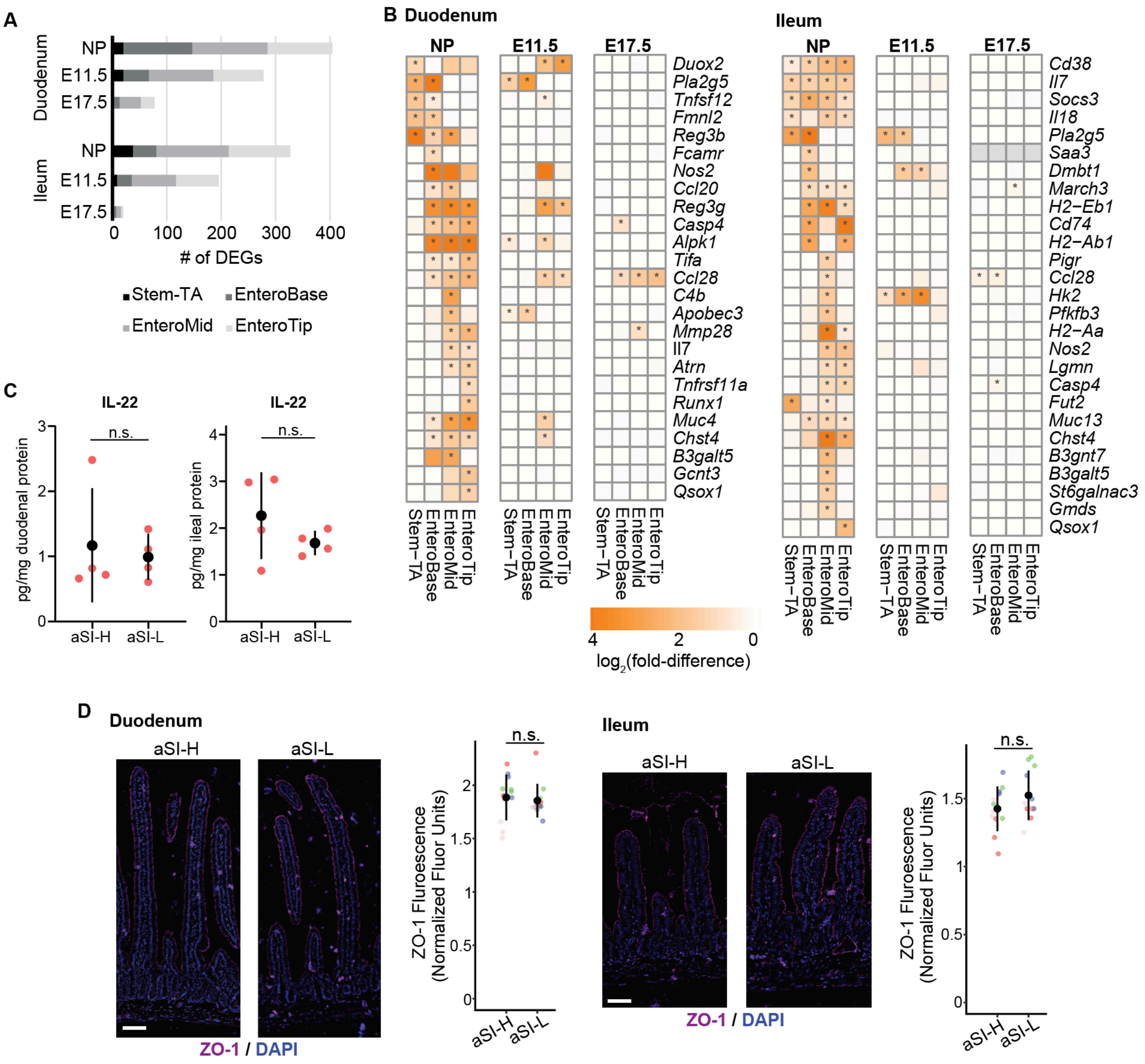


**Figure S3. Dampening of intestinal inflammation in aSI-L dams in pregnancy. (A)** snRNA-seq was performed on the duodenum and ileum at E11.5 and E17.5. Clusters representing intestinal stem and transit amplifying cells (Stem-TA), plus enterocytes from the base, middle, or tip of the villus (EnteroBase, EnteroMid, and EnteroTip) were identified (n = 4/intestinal segment/pregnancy time point/treatment group). The nonpregnant (NP) case is duplicated here from (*18*). DEGs in aSI-L versus aSI-H counterparts were identified in each cell cluster with DESeq2 ($|\log_2(\text{fold-difference})| > 0.5$, FDR $p < 0.05$, Wald test with BH correction) and are enumerated. **(B)** The relative expression levels of transcripts involved in immunoinflammatory pathways that were previously identified to be significantly elevated in non-pregnant aSI-L in comparison to aSI-H dams were assessed at E11.5 and E17.5. * $p < 0.05$. **(C)** Levels of IL-22 in intestinal tissue from aSI-H and aSI-L dams at E17.5 (n = 4/intestinal segment/treatment group). The Wilcoxon rank-sum test was used to assess statistical significance. **(D)** ZO-1 immunostaining of duodenal and ileal tissue segments from aSI-H and aSI-L dams at E17.5. Fluorescence intensity of staining at the villus periphery was quantified and normalized to background staining (n = 3 villi/intestinal segment were measured for n = 4 intestinal segments/treatment group). Mean ± SD are shown. A linear mixed-effects model *(Fluor ~ Microbiota + (1 | Mouse ID))* was used to assess statistical significance (scale bar = 50 μm).

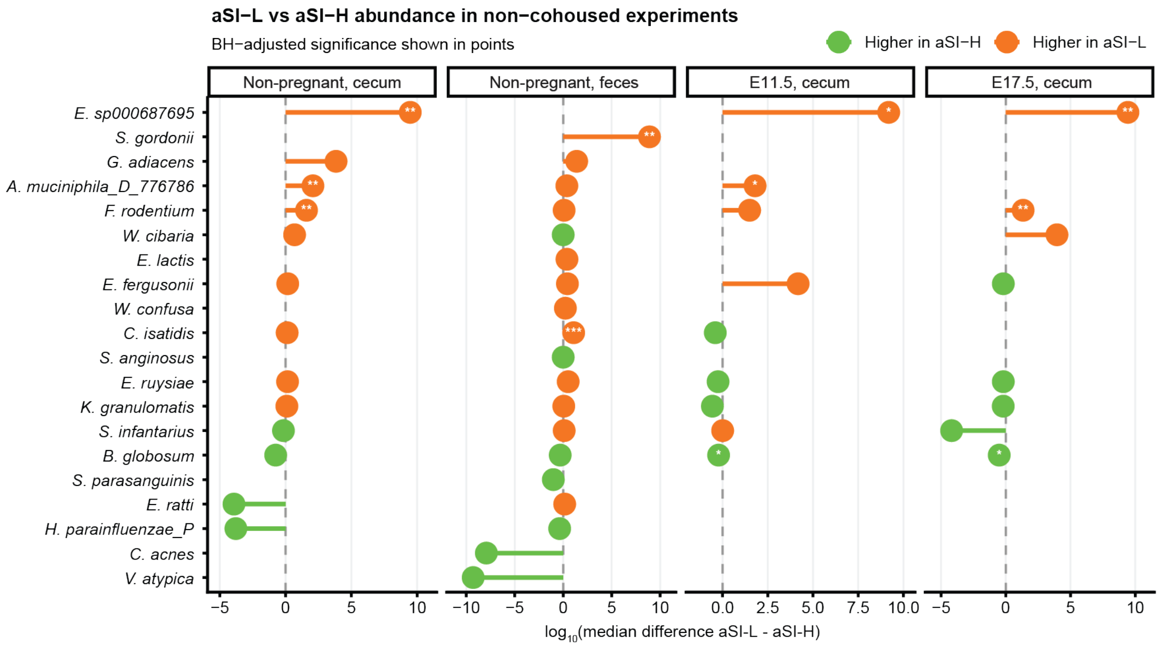


**Figure S4. Differential bacterial abundance in pregnant compared to non-pregnant controls.** The difference in log$_{10}$-transformed absolute bacterial abundance in the cecal contents and/or feces of non-pregnant controls (left) or pregnant dams (right). All bacterial taxa shown were detected in the duodenal aspirates or fecal samples of women. Median differences (n=4-6 mice per group) are shown, *, adjusted p < 0.1; **, adjusted p < 0.05; *** adjusted p < 0.01 (BH-corrected Wilcoxon rank-sum tests).

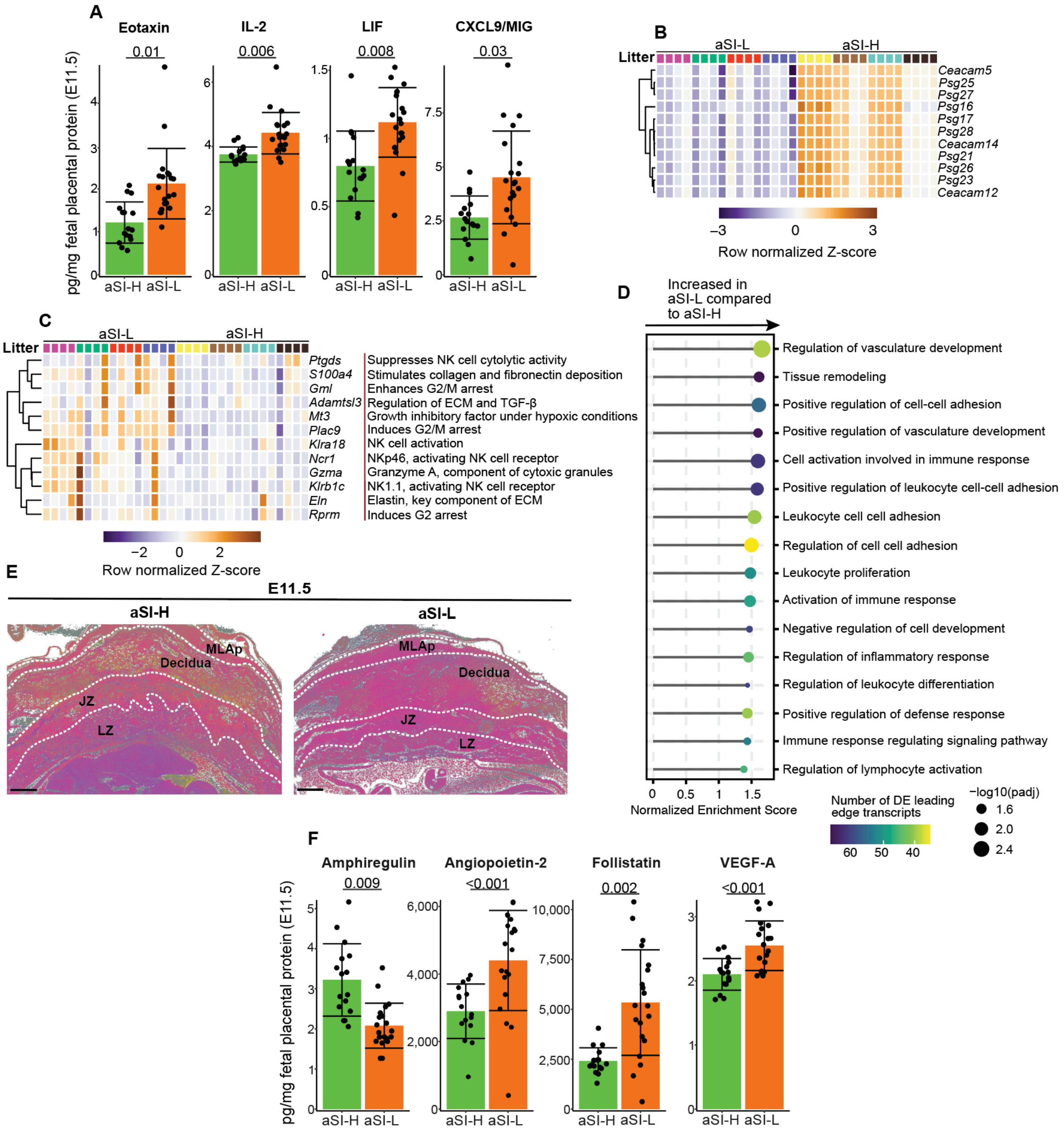


**Figure S5. Immune and ECM perturbations at E11.5 in aSI-L fetal-derived placentas. (A)** Proteomic analysis: levels of eotaxin, IL-2, LIF, and CXCL9 in placental homogenates (n = 4-5 dams/group, 4 placentas/dam). Adjusted P-values are shown as calculated using a linear mixed model (*Protein level ~ Microbiota + (1 | Litter ID)*) for pairwise comparisons. Mean values ± SD are shown. **(B)** Bulk RNA-seq-based differential expression of PSGs and CECAMs ($\log_2$(fold-difference) < -1.5; n = 4 litters/treatment group, 4 placentas/litter; DESeq2, FDR p < 0.05, Wald test with BH correction). **(C)** DEGs from bulk RNA-seq (|$\log_2$(fold-difference)| > 1.5; n = 4 litters/treatment group, 4 placentas/litter; DESeq2, FDR p < 0.05, Wald test with BH correction). **(D)** Selected GO Biological Process terms enriched in leading-edge transcripts identified from GSEA analysis DEGs in bulk RNA-Seq tables (FDR p < 0.05, Wilcoxon rank-sum test with BH correction). **(E)** Representative Movat-Russell pentachrome staining of sections from aSI-L and aSI-H placentas, with OD normalized colors to highlight collagen proteins (yellow). Scale bar = 400 µm. **(F)** Angiogenesis-associated proteins amphiregulin, angiopoietin-2, follistatin, and VEGF-A in the same placental homogenates used for the analysis described in panel **B**. Mean values ± SD are shown.

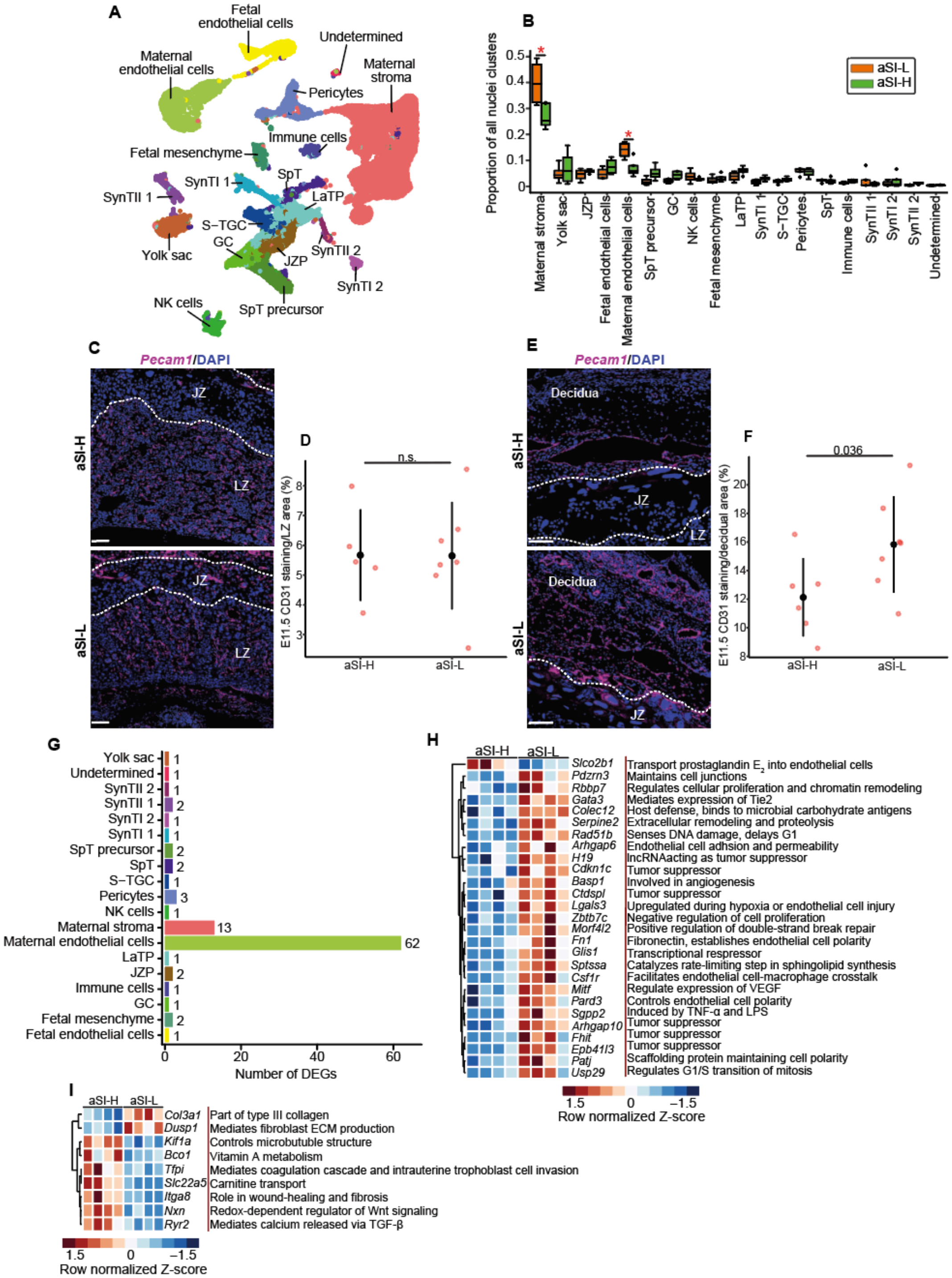


**Figure S6. snRNA-Seq-based analysis of differences in gene expression in maternal endothelial and stromal cells in E11.5 aSI-L versus aSI-H placentas. (A)** UMAP representation of 28,371 nuclei [across eight (4 aSI-L and 4 aSI-H) E11.5 placentas] assigned to 18 cell clusters. **(B)** Proportional representation of cell clusters identified by snRNA-seq from the E11.5 fetal placental preparations containing residual decidual tissue. Asterisks denote 'statistically credible differences' and diamonds denote outliers as defined by scCODA (**table**

**S4A** and *Methods*). The box plots represent upper and lower quartiles; the middle line is the median and whiskers indicate the range. **(C-F)** Representative images from FISH of *Pecam1* (CD31) staining and quantification in the LZ (**C**,**D**) and (**E**,**F**) decidua of placental cross-sections (n = 1-2 sections/placenta, 1-2 placentas/litter, 4 litters/treatment group). Means ± SD are shown. Adjusted p-values defined using a linear mixed model (*% Staining ~ Microbiota + (1 | Litter ID))* for pairwise comparisons. **(G)** Number of DEGs in each cell cluster identified by pseudobulk analysis of aSI-L versus aSI-H placentas (DESeq2, $|\log_2(\text{fold-difference})| > 0.5$, FDR $p < 0.05$, Wald test with BH correction). DEGs identified when comparing aSI-L to aSI-H in **(H)** maternal endothelial cells ($|\log_2(\text{fold-difference})| > 1$) and **(I)** maternal stroma ($|\log_2(\text{fold-difference})| > 0.5$).

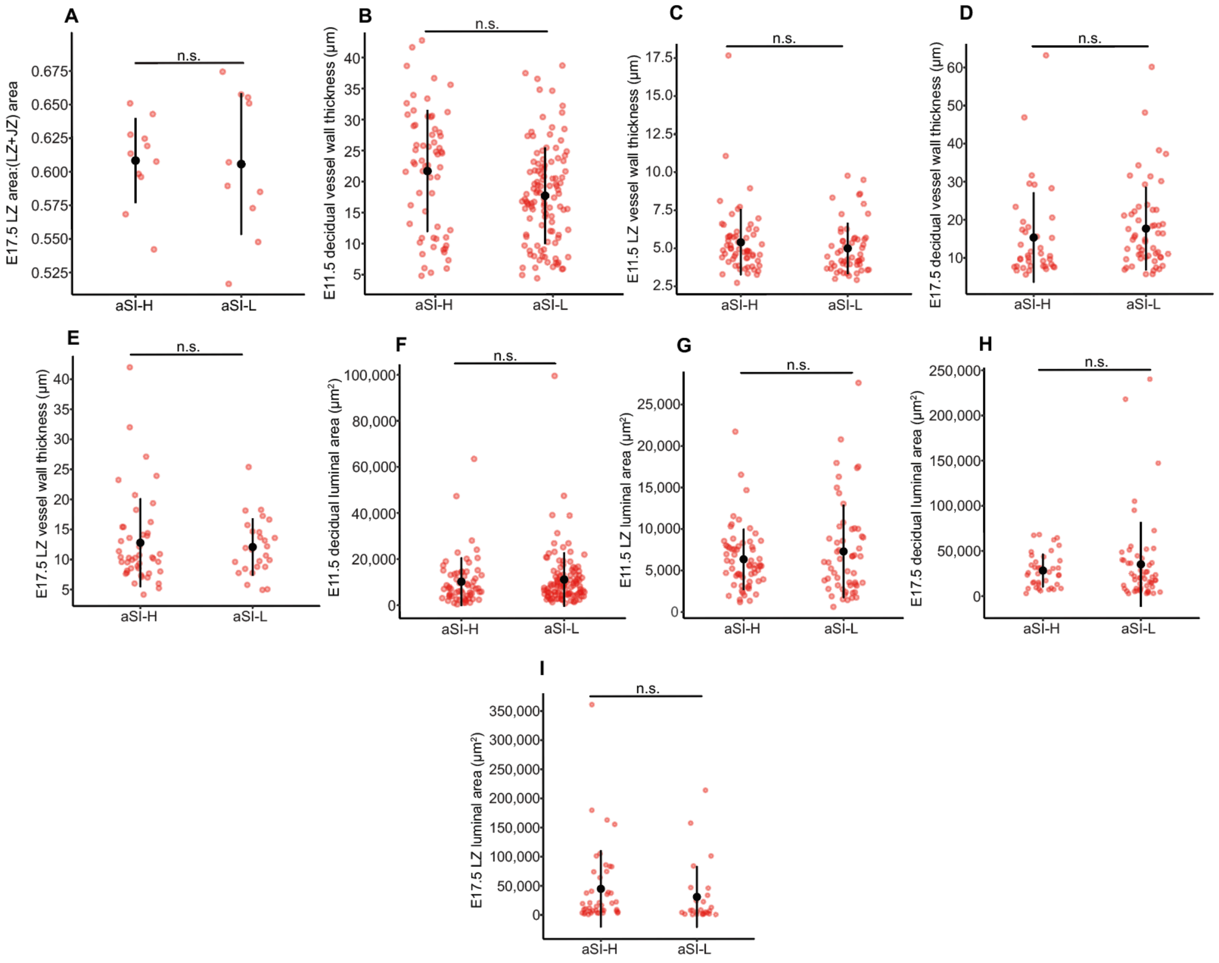


**Figure S7. Blood vessel histomorphometrics in the LZ and decidua in E11.5 and E17.5 aSI-L and aSI-H placentas.** (**A**) The ratio of E17.5 LZ area to LZ + JZ area (n = 1-2 placentas/litter, 6-7 litters/treatment group). Statistical significance was defined using a linear mixed model (*Ratio ~ Microbiota + (1 | Litter ID))*. Mean values ± SD are plotted. (**B-E**) Histomorphometric measurements. E11.5 decidual and LZ arterial wall thickness (panels **B** and **C**). E17.5 decidual and LZ arterial wall thickness (panels **D** and **E**). **(F-I)** Histomorphometric measurements of E11.5 **(F)** decidual and **(G)** LZ luminal areas (panel **F** and **G**), and E17.5 decidual and LZ luminal areas (panels **H** and **I**) (n = 1-2 placentas/litter, 4-5 litters/treatment group/timepoint). Statistical significance was defined using the linear mixed model (*Feature ~ Microbiota + (1 | Litter ID))*. Mean values ± SD are shown.

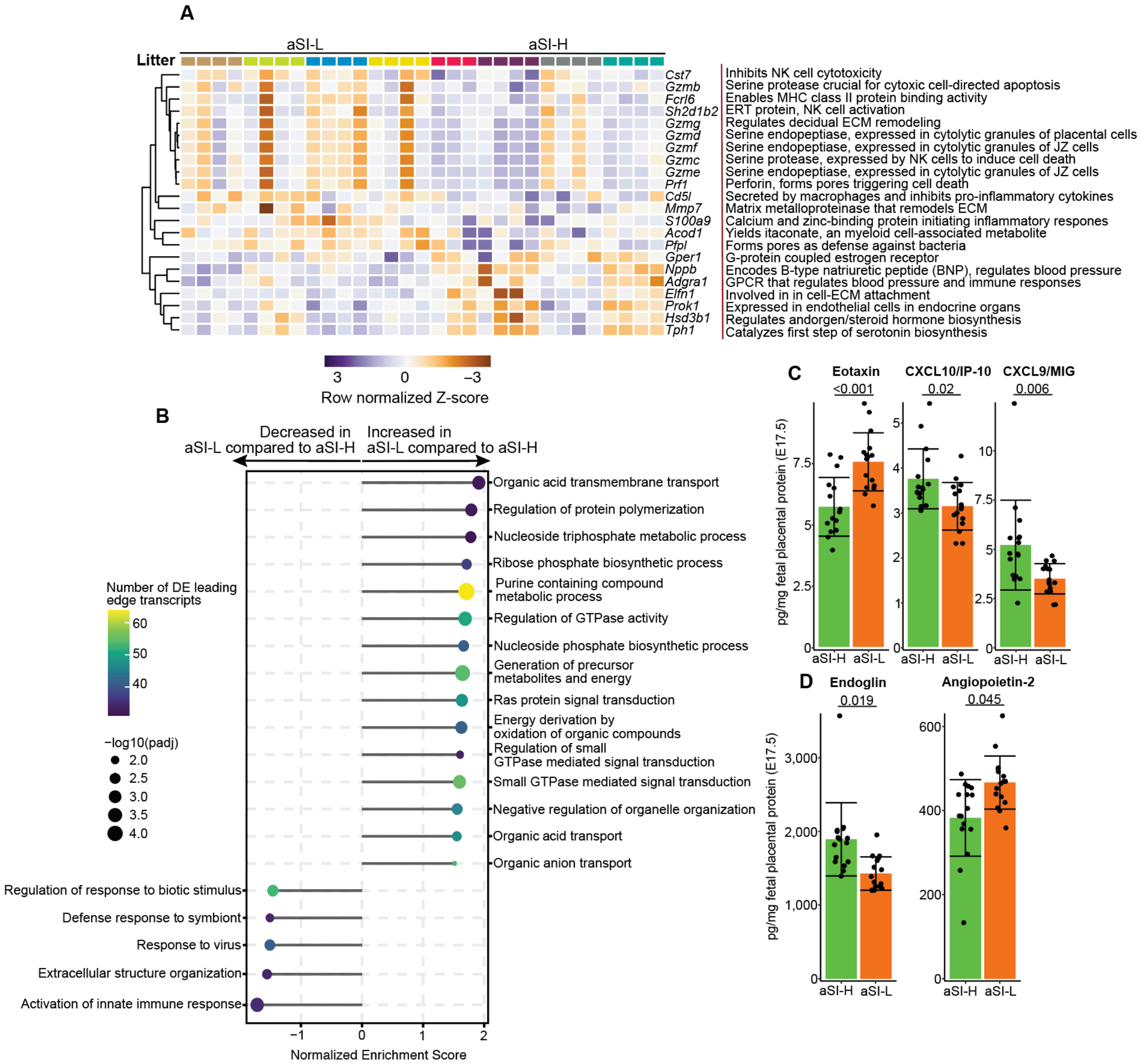


**Figure S8. Bulk RNA-seq and proteomic analyses of E17.5 aSI-L and aSI-H placentas. (A)** DEGs from bulk RNA-seq analysis of E17.5 aSI-L and aSI-H 'fetal' placental preparations ($|\log_2(\text{fold-difference})| > 1$; n = 4 litters/treatment group, 4 placentas/litter; DESeq2, FDR $p < 0.05$ (Wald test with BH correction). **(B)** Selected GOBP terms enriched in leading-edge transcripts (FDR p-value < 0.05, Wilcoxon rank-sum test with BH correction). **(C)** Immune-associated proteins CXCL10/IP-10, and CXCL9 quantified in placental homogenates (n = 4-5 dams/group, 2-4 placentas/dam). Adjusted p-values for pairwise comparisons, calculated using a linear mixed model (*Protein level ~ Microbiota + (1 | Litter ID)*) **(D)** Angiogenesis-associated proteins endoglin and angiopoietin-2 quantified as in panel **C**. Mean values ± SD are shown in panels **C** and **D**.

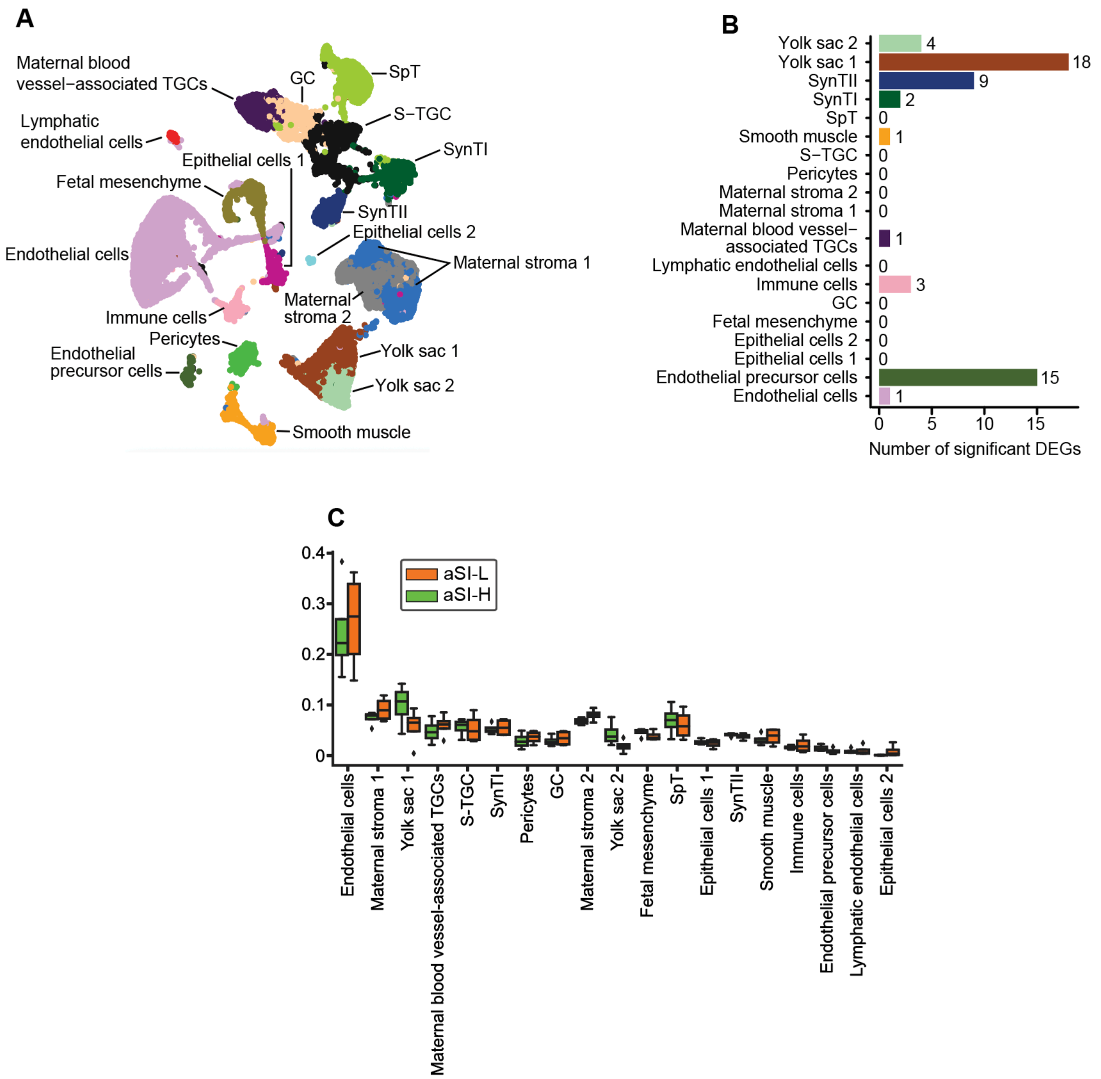


**Figure S9. snRNA-Seq analysis of gene expression in E17.5 aSI-L versus aSI-H fetal placentas. (A)** UMAP representation of 35,834 nuclei across eight (4 aSI-L and 4 aSI-H) E17.5 fetal placentas and assigned to 19 cell clusters. **(B)** Number of DEGs in each cell cluster identified by pseudobulk analysis of E17.5 aSI-L versus aSI-H fetal placentas (DESeq2, $|\log_2(\text{fold-difference})| > 0.5$, FDR $p < 0.05$ (Wald test with BH correction). **(C)** Proportional representation of cell clusters identified by snRNA-seq from E17.5 fetal placentas. Asterisks denote ‘statistically credible differences’ and diamonds denote outliers as defined by scCODA (see **table S4B** and *Methods*). The box plots represent upper and lower quartiles, the middle line is the median and whiskers indicate the range.

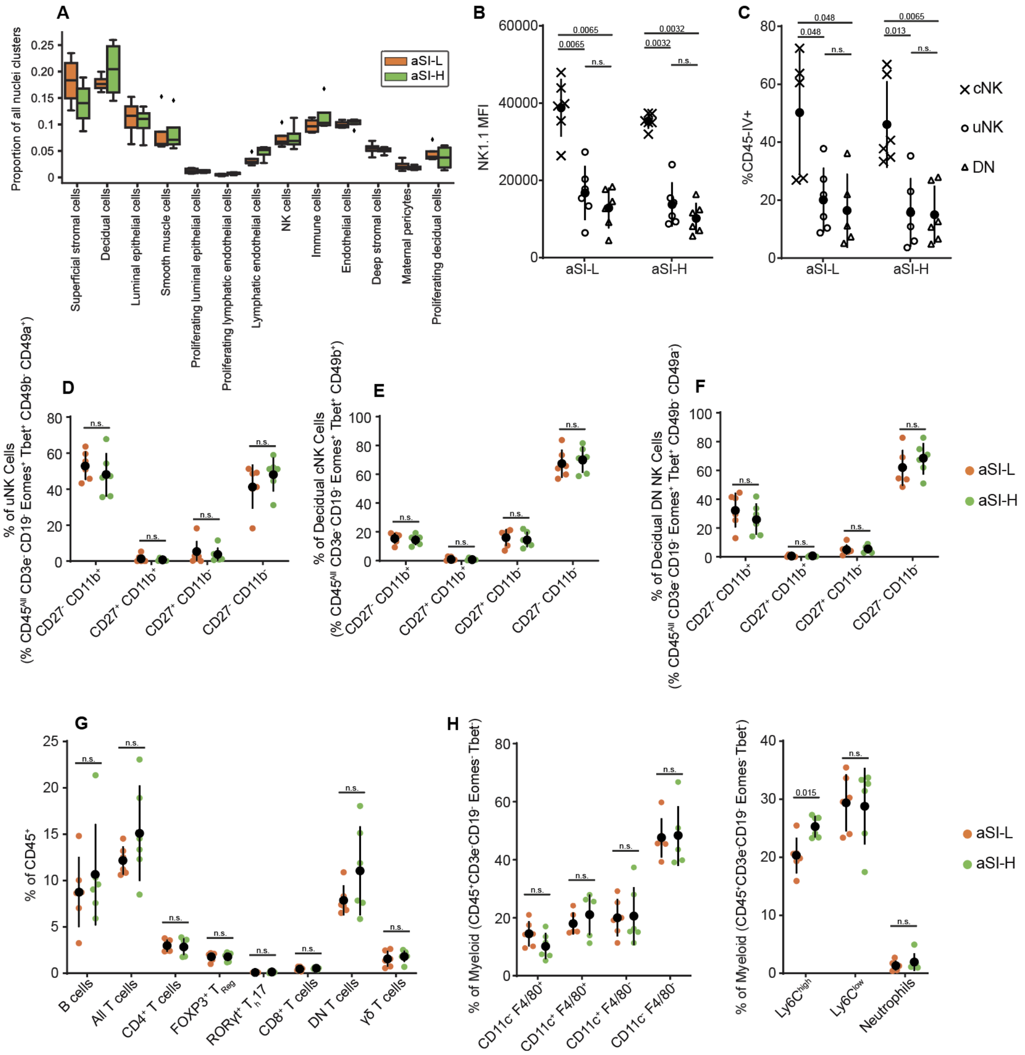


**Figure S10. Immune cell flow cytometry and cluster proportion defined by snRNA-Seq in E11.5 deciduas and spleens. (A)** Proportional representation of cell clusters identified by snRNA-seq from E11.5 deciduas/MLAps. Asterisks denote 'statistically credible differences' and diamonds denote outliers as defined by scCODA (**table S7A** and *Methods*). The box plots represent upper and lower quartiles; the middle line is the median; whiskers indicate the range. **(B)** Median fluorescence intensity of NK1.1 immunostaining and (**C**) proportion of CD45-IV staining in cNK, uNK, and DN NK cells in the E11.5 decidua. (**D-F**) Proportion of CD27 and CD11b expression in **(D)** uNK, **(E)** cNK, and **(F)** DN NK cell populations in E11.5 decidua. (**G,H**) Proportion of decidual **(G)** B or T cells and **(H)** myeloid cells. All deciduas in each litter were pooled (n = 6 dams/treatment group, Wilcoxon rank-sum test with BH correction for multiple comparisons). Mean values ± SD are shown. Gating of immune cell populations was performed as depicted in **fig. S11A**.

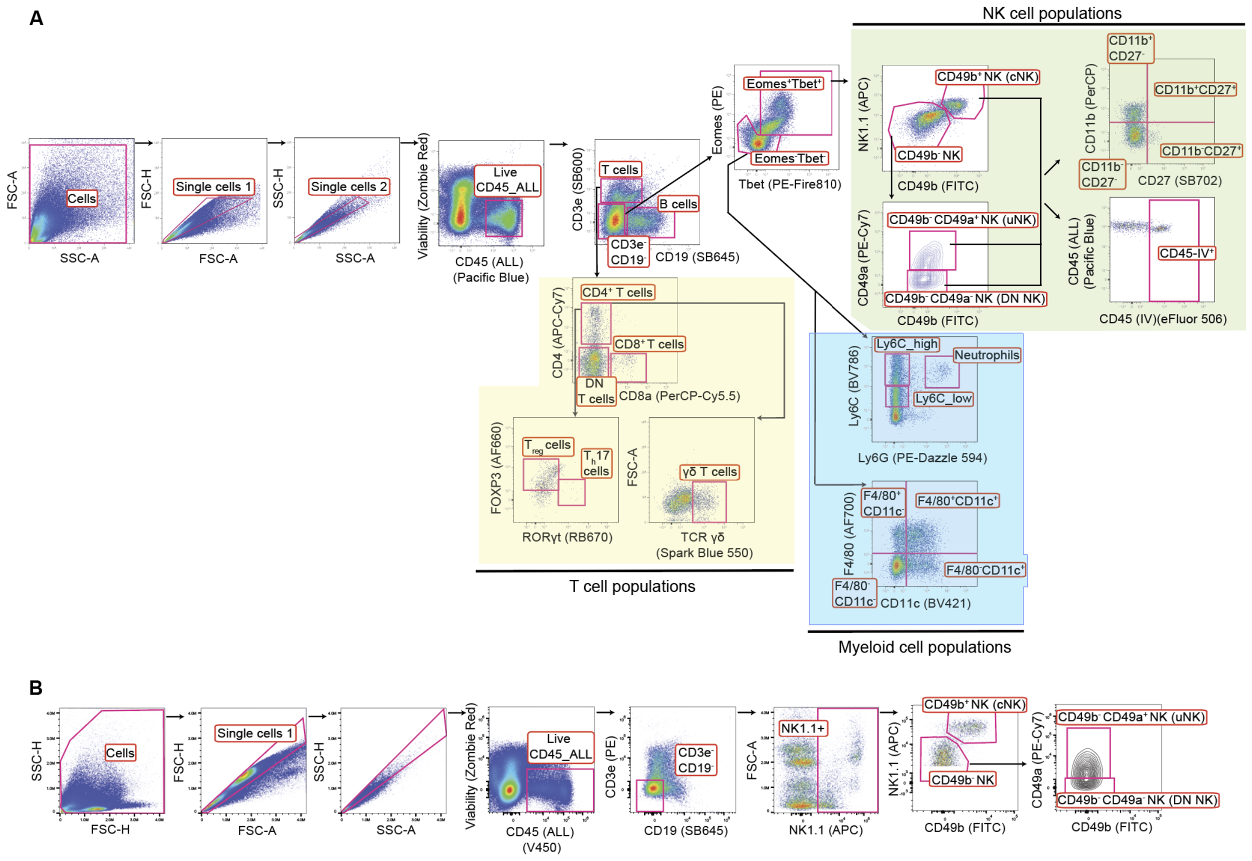


**Figure S11. Gating scheme for flow cytometry analysis of immune cell populations in E11.5 decidual tissue. (A)** Gating scheme for identifying NK, lymphocyte, and myeloid cell populations by flow cytometry for **Fig. 2G** and **fig. S10B-H**. All deciduas in a litter were pooled into a single sample. **(B)** Gating scheme for identifying NK cell populations by flow cytometry for **Fig. 6B**. All deciduas in a litter were pooled into a single sample.

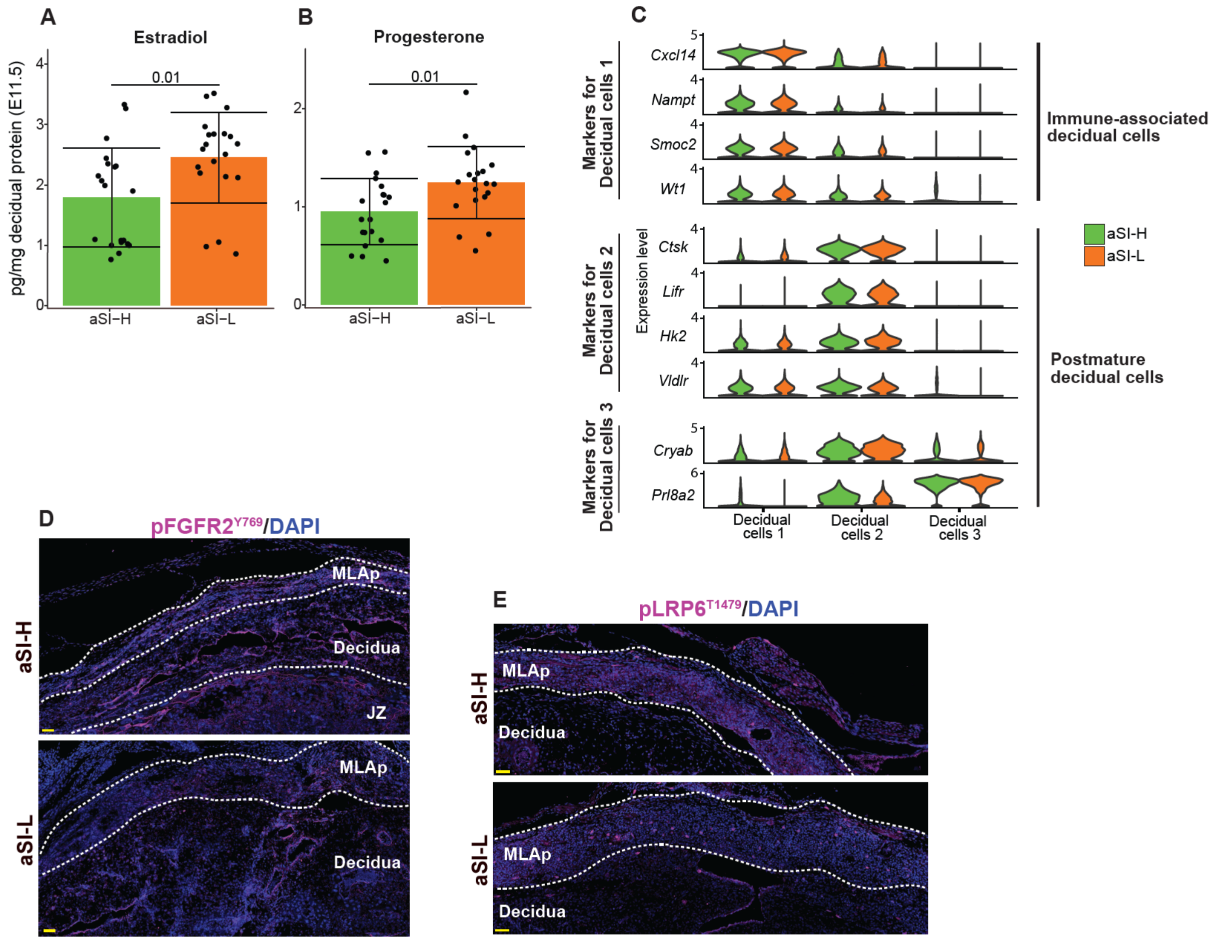


**Figure S12. Characterization of E11.5 decidual stromal cells. (A,B)** Estradiol and progesterone quantification in decidual/MLAp homogenates (n = 5 dams/group, 3-4 placentas/dam). Adjusted p-values for pairwise comparisons calculated using a linear mixed model (*Protein level ~ Microbiota + (1 | Litter ID)*). Mean values ± SD are shown. **(C)** Expression of immune-associated decidual cell and postmature decidual cell-associated genes in type 1, 2, and 3 clusters defined from the spatial transcriptomics table. **(D,E)** Representative example of immunocytochemical staining for pFGFR2(Y769) (panel **D**) and pLRP6(T1479) (panel **E**) in E11.5 aSI-L and aSI-H deciduas and MLAps. Scale bar = 50 µm.

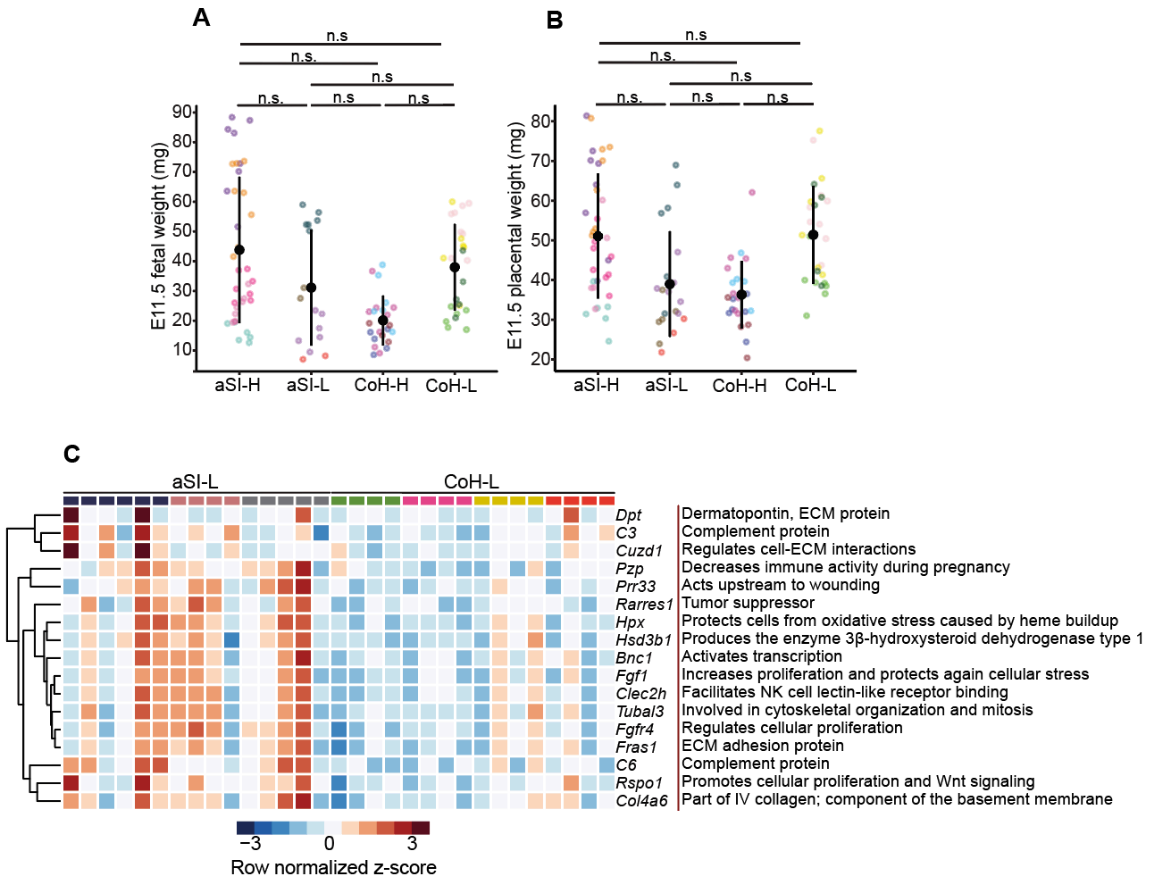


**Figure S13. Co-housing effects on immune and ECM transcriptional responses in E11.5 placentas.** **(A)** E11.5 fetal weights [aSI-L: 4 dams, 16 fetuses, 31.2 ± 19.5 mg (mean ± SD); aSI-H: 5 dams, 36 fetuses, 43.8 ± 24.6 mg; CoH-L: 4 dams, 25 fetuses, 37.9 ± 14.6 mg; CoH-H: 4 dams, 23 fetuses, 20.1 ± 8.4 mg] and **(B)** Corresponding E11.5 placental weights [aSI-L: 38.6 ± 13.3 mg; aSI-H: 50.7 ± 15.7 mg; CoH-L: 51.1 ± 12.3 mg; CoH-H: 35.9 ± 8.5 mg]. Each litter is represented by a different color. **(C)** DEGs from bulk RNA-seq of E11.5 aSI-L and CoH-L fetal placental preparations ($|\log_2(\text{fold-difference})| > 1.5$; n = 3-4 litters/treatment group, 4-6 placentas/litter; DESeq2, FDR p-value < 0.05, Wald test with BH correction).

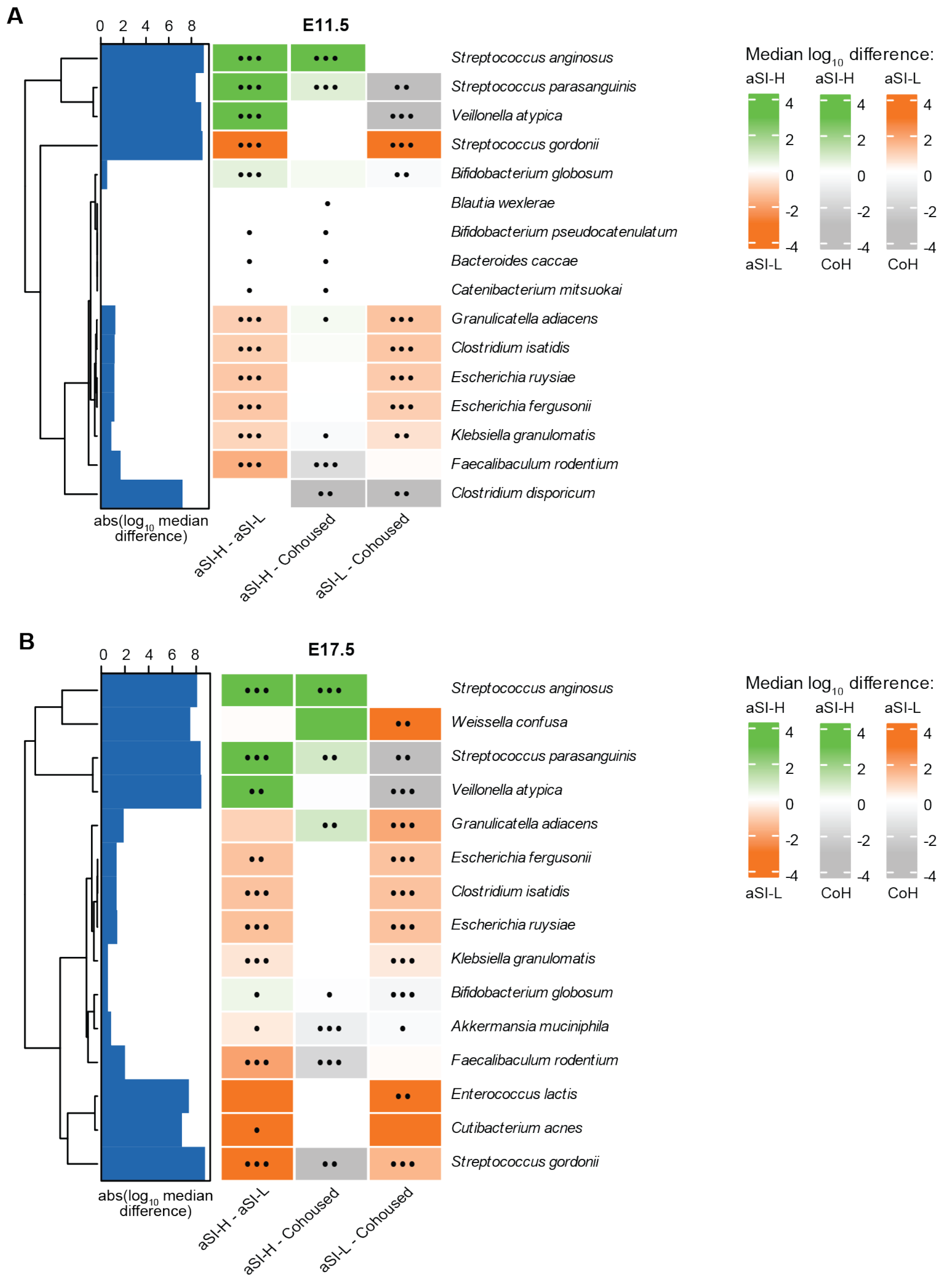


**Figure S14. Differential bacterial abundances in the cecal contents of E11.5 and E17.5 co-housed and control (aSI-L, aSI-H) pregnant dams.** Heatmap shows bacterial species whose absolute abundances were significantly different in at least one pairwise comparison shown. •, adjusted $p < 0.1$; ••, adjusted $p < 0.05$; •••, adjusted $p < 0.01$ (Kruskal-Wallis test with Dunn's multiple comparisons, BH correction). Heatmap color denotes the median difference in log$_{10}$-transformed absolute abundance between aSI-L (orange), aSI-H (green), and co-housed (CoH, gray) (**A**) E11.5 and (**B**) E17.5 dams. The bar chart to the left of the heatmap shows the absolute value of the log$_{10}$ difference. For the full list of differentially abundant taxa, see **Table S9**.

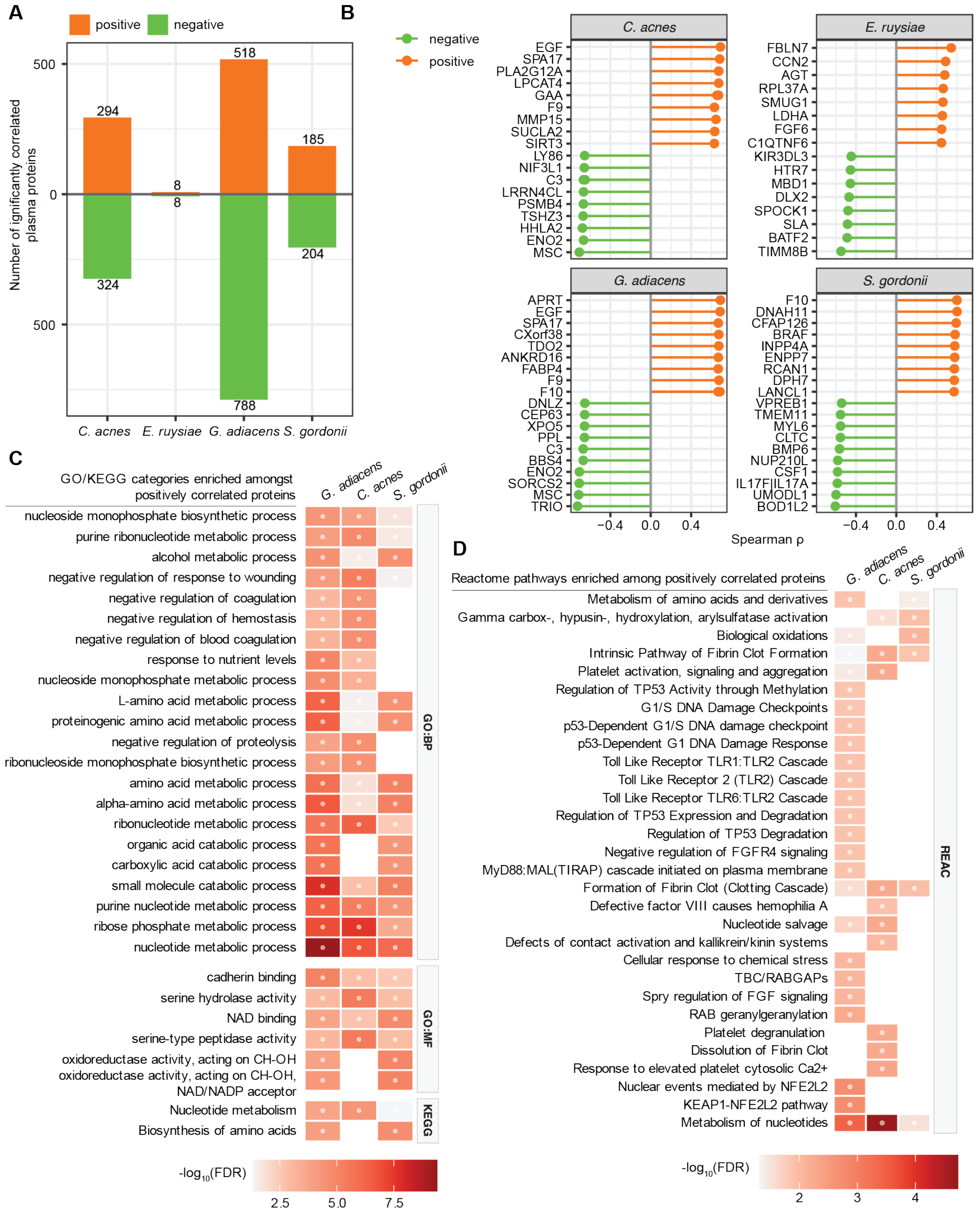


**Figure S15. Correlations between the abundances of aSI-L and aSI-H bacteria in duodenal aspirates and plasma proteins of women in the clinical trials.** Bacteria that were implicated in fetal growth based on co-housing experiments performed in gnotobiotic mice were identified in the duodenal aspirates of women. Their absolute abundances in duodenal aspirates were correlated with levels of plasma proteins in the same participants. (**A**) The total number of significantly correlated plasma proteins with the absolute abundances of aSI-L bacteria (bacteria were included if prevalence in duodenal aspirates of women was >12.5%. Spearman correlations, FDR-corrected p-values). (**B**) The ten plasma proteins most significantly positively and negatively correlated with the abundance of bacteria in duodenal aspirates. (**C,D**) GSEA was performed on plasma proteins that were significantly positively correlated with the absolute abundances of aSI-L species in duodenal aspirates. The 30 Gene Ontology (GO) or KEGG categories (**C**) and Reactome pathways (**D**) that were most strongly enriched are included (•, FDR-corrected $p < 0.01$ for **C**, $p < 0.05$ for **D**; see **table S11** for the full list of correlations and enrichment results).

| Timepoint | Results |
| --- | --- |
| **E11.5** | ↑ Maternal immune transcriptional pathways (GOBP) |
| | ↑ Maternal NK cell activation transcripts (e.g. *Klra18*, *Ncr1*, *Gzma*, *Klrb1c, Ptgds*) |
| | ↑ Pro-inflammatory cytokines (eotaxin, IL-2, LIF, and CXCL9 protein concentrations) |
| | ↑ Maternal angiogenesis (*Pecam1* staining, angiopoietin-2, follistatin, VEGF-A protein concentrations) |
| | ↑ Maternal vascular development transcriptional pathways (GOBP) |
| | ↓ ECM-related transcripts (e.g. *Adamtsl3*, *Eln*, *S100a4*) |
| | ↓ Collagen deposition (Movat-Russel's pentachrome) |
| | ↑ Cellular proliferation of maternal-derived placental tissue (scCODA) |
| | ↑ Tumor suppressor transcription (e.g. *Plac9*, *Mt3*, *Gml*, *Rprm*) |
| **E17.5** | ↓ Maternal immune transcriptional pathways (GOBP) |
| | ↑ Maternal NK cell cytotoxicity (e.g. *Fcrl6*, *Sh2d1b2*, *Prf1, Cst7*) and granzymes (e.g. *Gzmb*, *Gzmg*, *Gzmd*, *Gzmf*, *Gzmc*, *Gzme*) transcripts |
| | ↓ Chemokines recruiting NK and T cells (CXCL9 and CXCL10 protein concentrations) |
| | ↑ Organic acid transport and energy derivation transcriptional pathways (GOBP) |
| | ↑ Nucleic acid-related transcriptional pathways (GOBP) |
| | ↓ Vascular tone (e,g. *Nppb*, *Adgra1*, *Tph1*) |
| | ↓ ECM-related transcripts (e.g. *Elfn1)* |
| | ↓ ECM organization transcriptional pathways (GOBP) |

**Table S4.** Summary of the findings for fetal placenta (*i.e.* decidua and MLAp removed, LZ and JZ concentrated) at E11.5 and E17.5 in aSI-L versus aSI-H groups.

| Reagent | Type | Host/Conjugate/Channel or Fluorophore | Dilution (μg/mL) | Vendor | Catalog no. |
|---|---|---|---|---|---|
| *Pecam1* (CD31) | *ISH* Probe | C3 | - | ACDBio | 316721 |
| Phospho-TGFBR1 (Ser165) Polyclonal Antibody | 1° | Rabbit | 5 | AbboMax | 620-910 |
| Phospho-TGFBR1 (Thr204) Polyclonal Antibody | 1° | Rabbit | 10 | Invitrogen | PA5-106155 |
| TGF beta-1 Recombinant Monoclonal Antibody | 1° | Rabbit | 5 | Invitrogen | MA5-44667 |
| TGF beta-2 Recombinant Monoclonal Antibody | 1° | Rabbit | 0.5 | Abcam | ab323293 |
| Phospho-FGFR2 (Tyr769) Polyclonal Antibody | 1° | Rabbit | 5 | Invitrogen | PA5-105880 |
| Phospho-LRP6 (Thr1479) Polyclonal Antibody | 1° | Rabbit | 10 | Invitrogen | PA5-106063 |
| ZO-1 Polyclonal Antibody | 1° | Rabbit | 0.25 | Invitrogen | 61-7300 |
| Opal 690 | Fluorescent dye | 690 | - | Akoya Biosciences | NC1605064 |
| Streptavidin, Alexa Fluor 647 Conjugate | Fluorescent dye | Alexa Fluor 647 | 2 | Invitrogen | S32357 |
| Alexa Fluor 647 Anti-Rabbit IgG (H+L), Highly Cross-Absorbed | 2° | Goat, Alexa Fluor 647 | 2 | Invitrogen | A21245 |

**Table S12.** Reagents used for *in situ* hybridization or immunohistochemical analyses

| Protein Marker | Fluorophore | Host | Clone | Lot | Vendor | Catalog # | Dilution |
|---|---|---|---|---|---|---|---|
| CD49a | PE-Cy7 | Armenian Hamster | HMa1 | B374852 | BioLegend | 142607 | 1/200 |
| CD45 [All] | Pacific Blue | Mouse | QA17A26 | B435224 | BioLegend | 157620 | 1/500 |
| Eomes | PE | Rat | W17001A | B430797 | BioLegend | 157706 | 1/200 |
| FOXP3 | Alexa Fluor 660 | Rat | FJK-16s | 3117198 | Invitrogen | 606-5773-80 | 1/100 |
| Ly6C | BV786 | Rat | HK1.4 | 5114188 | BD OptiBuild | 755197 | 1/200 |
| CD3ε | Super Bright 600 | Armenian hamster | 145-2C11 | 2939287 | Invitrogen | 63-0031-82 | 1/100 |
| CD45 [IV] | eFluor 506 | Rat | 30-F11 | 3042087 | Invitrogen | 69-0451-82 | IV injection: 1/4 |
| Ly6G | PE/Dazzle 594 | Rat | 1A8 | B439210 | BioLegend | 127647 | 1/100 |
| TCR γ/δ | Spark Blue 550 | Armenian hamster | GL3 | B445902 | BioLegend | 285241 | 1/100 |
| Tbet | PE/Fire 810 | Mouse | 4B10 | B450836 | BioLegend | 644839 | 1/200 |
| RORγt | RB670 | Mouse | Q31-378 | 4261321 | BD Horizons | 571952 | 1/100 |
| NK1.1 | APC | Mouse | PK146 | B341103 | BioLegend | 108710 | 1/200 |
| CD49b | FITC | Armenian hamster | HMa2 | B297130 | BioLegend | 103504 | 3/200 |
| CD19 | Super Bright 645 | Rat | eBio1D3 (1D3) | 2630545 | Invitrogen | 64-0193-82 | 1/200 |
| CD27 | Super Bright 702 | Armenian hamster | LG.7F9 | 2630556 | Invitrogen | 67-0271-82 | 1/200 |
| CD11c | BV421 | Armenian hamster | N418 | 6181749 | BD Biosciences | 565452 | 1/200 |
| CD8a | PerCP-Cy5.5 | Rat | 53-6.7 | E08300-1631 | eBioscience | 45-0081-82 | 1/200 |
| CD4 | APC-Cy7 | Rat | RM4-5 | B208365 | BioLegend | 100526 | 1/100 |
| F4/80 | Alexa Fluor 700 | Rat | BM8 | B338811 | BioLegend | 123130 | 1/200 |
| Zombie Red (Viability) | - | - | - | B407223 | BioLegend | 423110 | 1/1000 |
| CD11b | PerCP | Rat | M1/70 | B204232 | BioLegend | 101229 | 1/100 |

**Table S13.** Antibodies employed for flow cytometry in **Fig. 2**

| Protein Marker | Fluorophore | Host | Clone | Lot | Vendor | Catalog # | Dilution |
|---|---|---|---|---|---|---|---|
| CD49a | PE-Cy7 | Armenian Hamster | HMa1 | B374852 | BioLegend | 142607 | 1/200 |
| CD45 [All] | V450 | Rat | 30-F11 | 1210486 | BD Horizon | 560501 | 1/200 |
| CD45 [IV] | Alexa Fluor 647 | Rat | S18009D | B363360 | BioLegend | 160304 | IV injection: 7/40 |
| CD3ε | PE | Rat | 17A2 | 1005829 | BD Biosciences | 555275 | 1/100 |
| NK1.1 | APC | Mouse | PK146 | B341103 | BioLegend | 108710 | 1/200 |
| CD49b | FITC | Armenian Hamster | HMa2 | B297130 | BioLegend | 103504 | 3/200 |
| CD19 | Super Bright 645 | Rat | eBio1D3 (1D3) | 2630545 | Invitrogen | 64-0193-82 | 1/200 |

**Table S14.** Antibodies employed for flow cytometry in **Fig. 6**
